## Supplementary figures and images for "Convergent Evolution of H4K16ac-mediated Dosage Compensation Shapes Sex-dependent Lifespan in a ZW Species"

### S1 Fig.

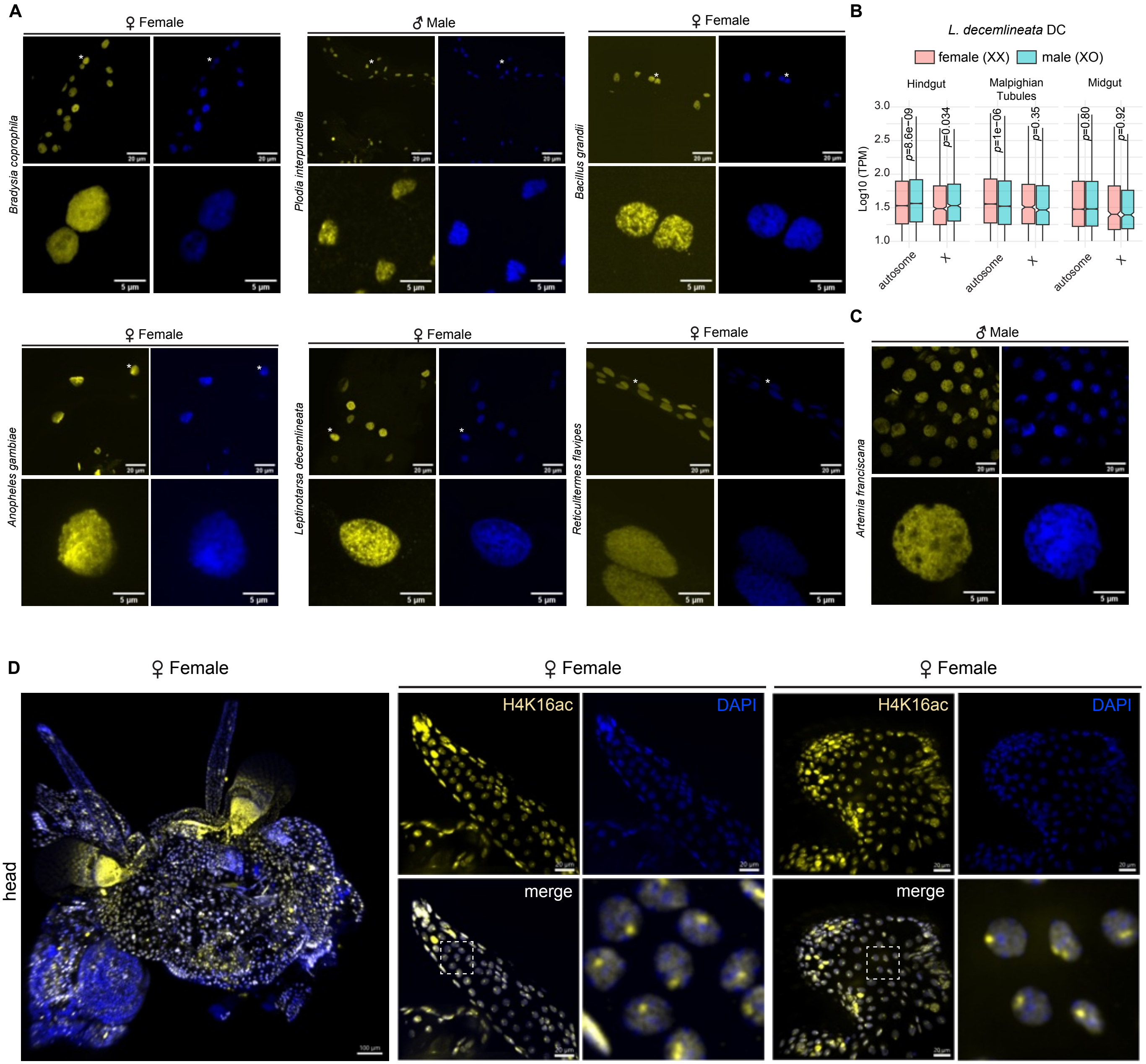

### S2 Fig.

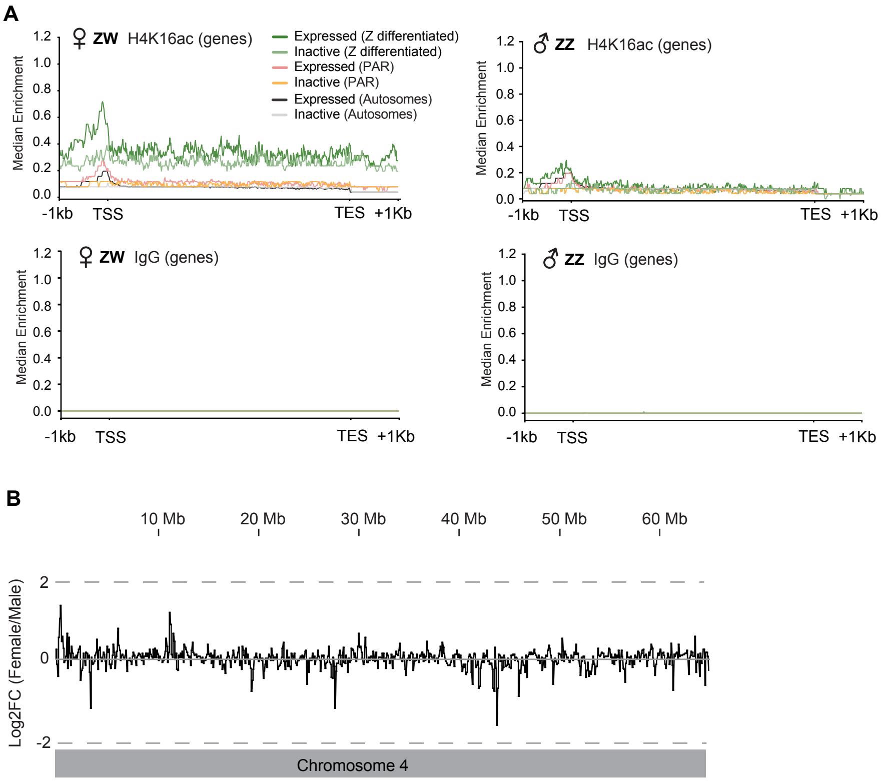

### S3 Fig.

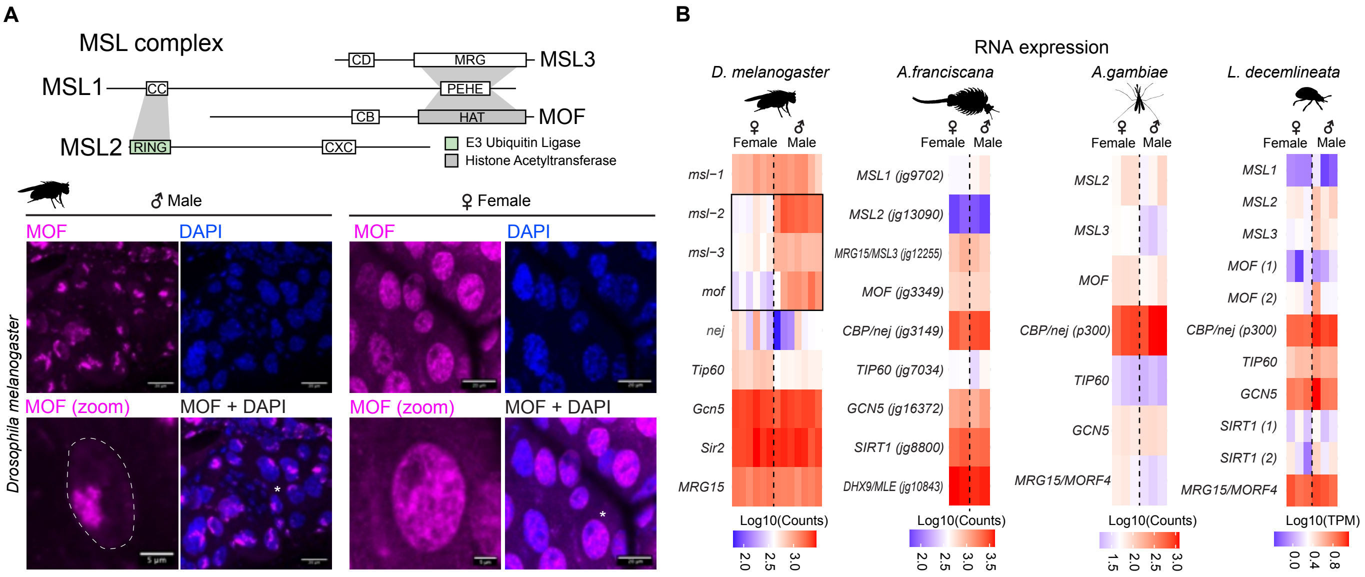

### S4 Fig.

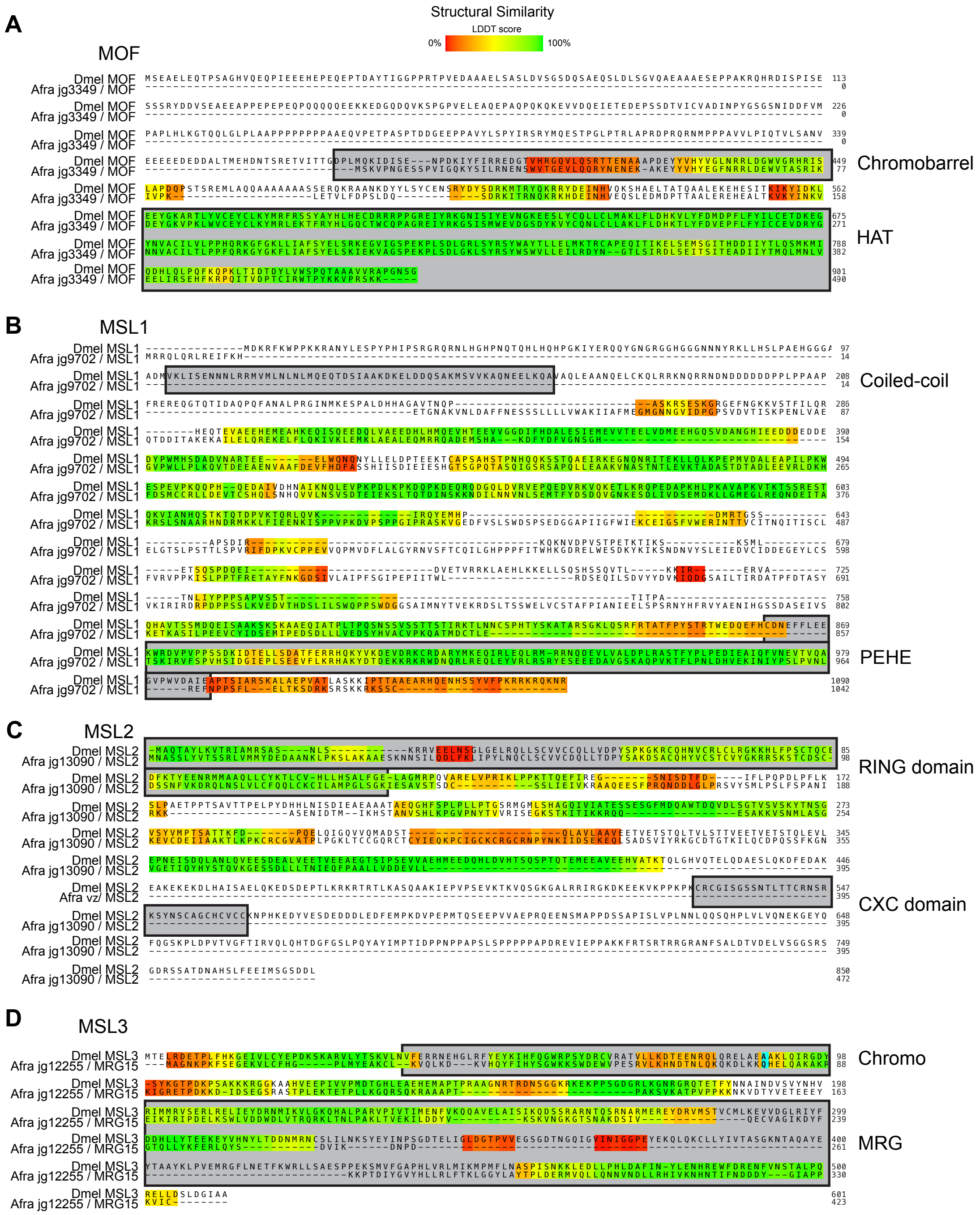

### S5 Fig.

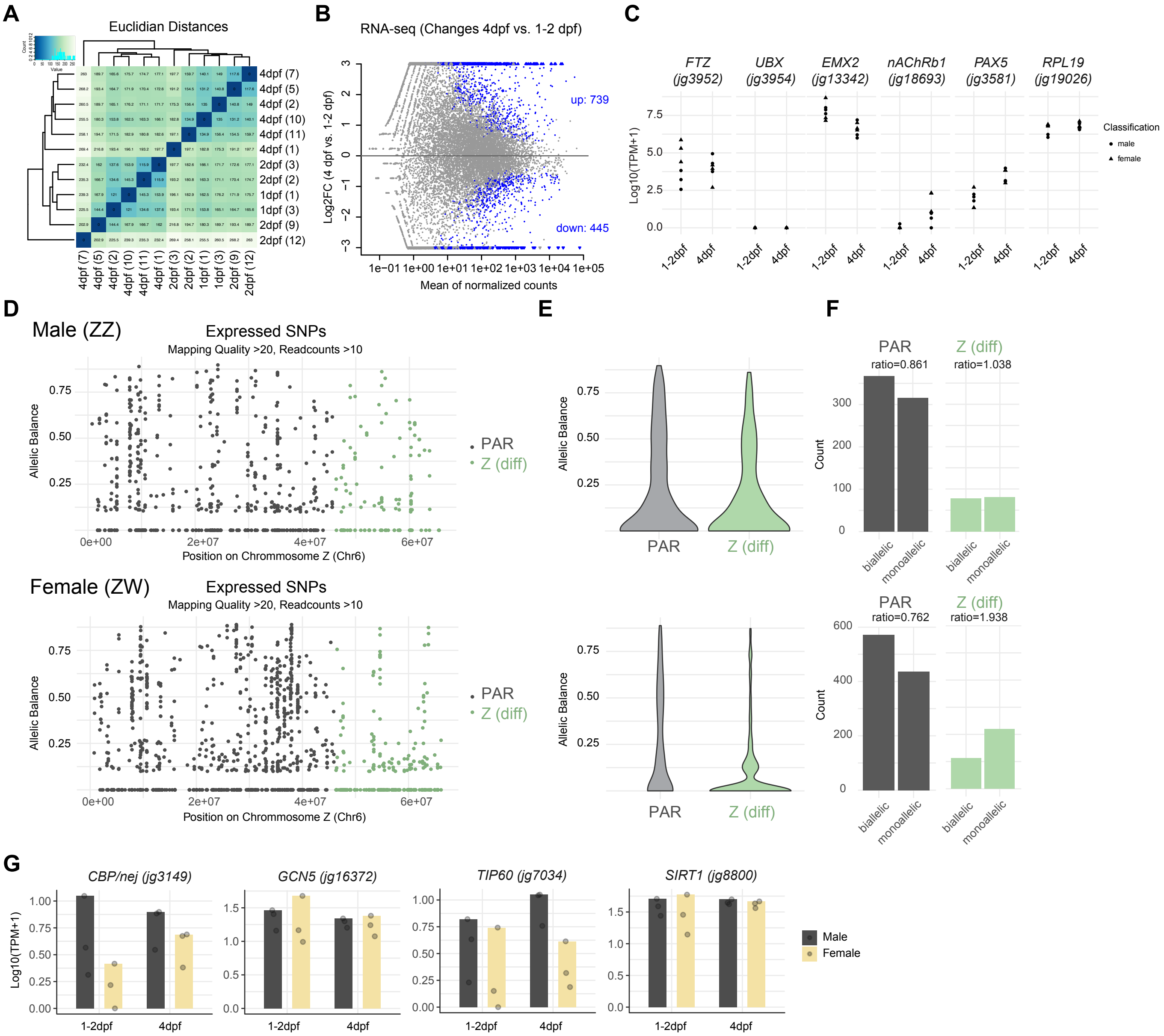

### S6 Fig.

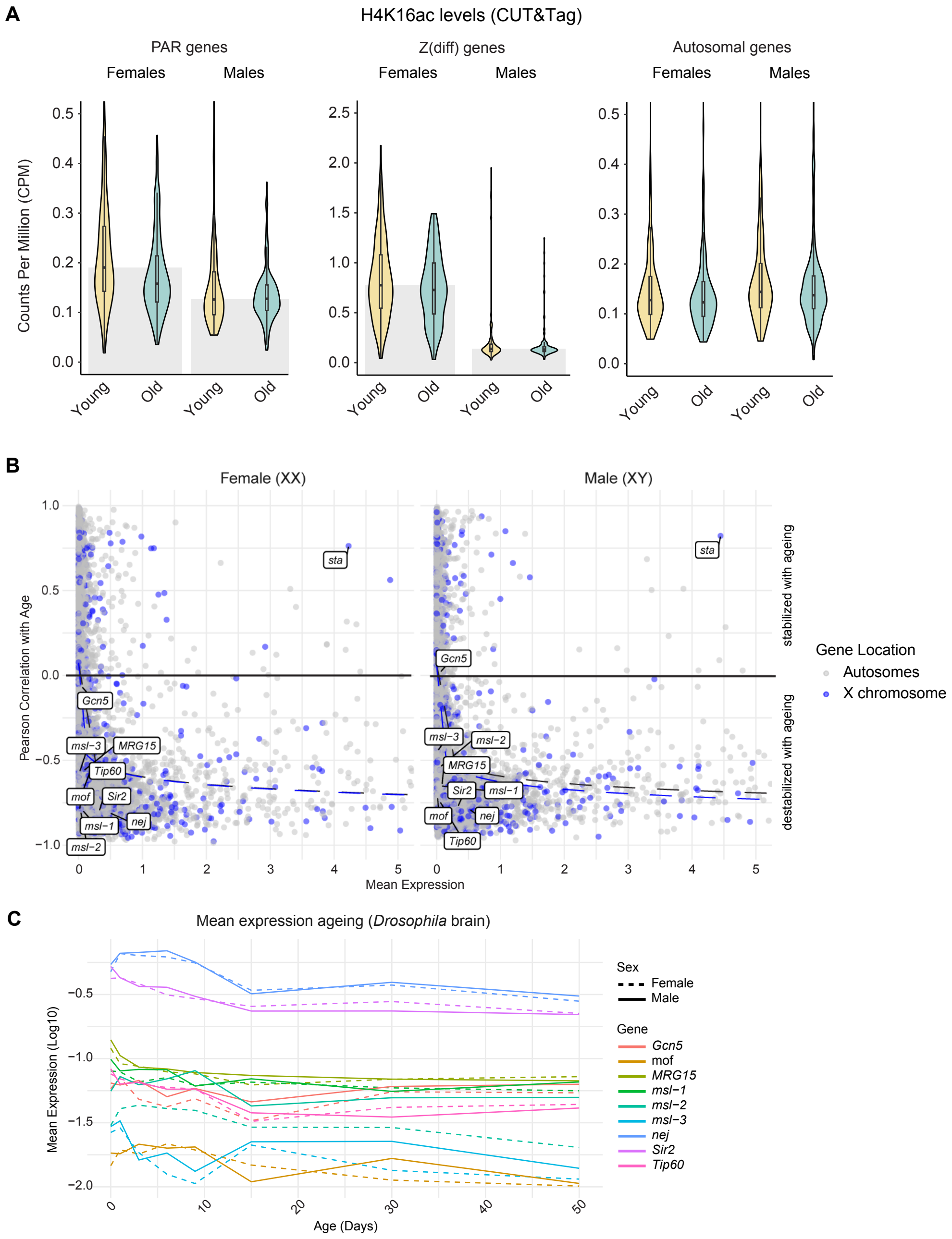

### S7 Fig

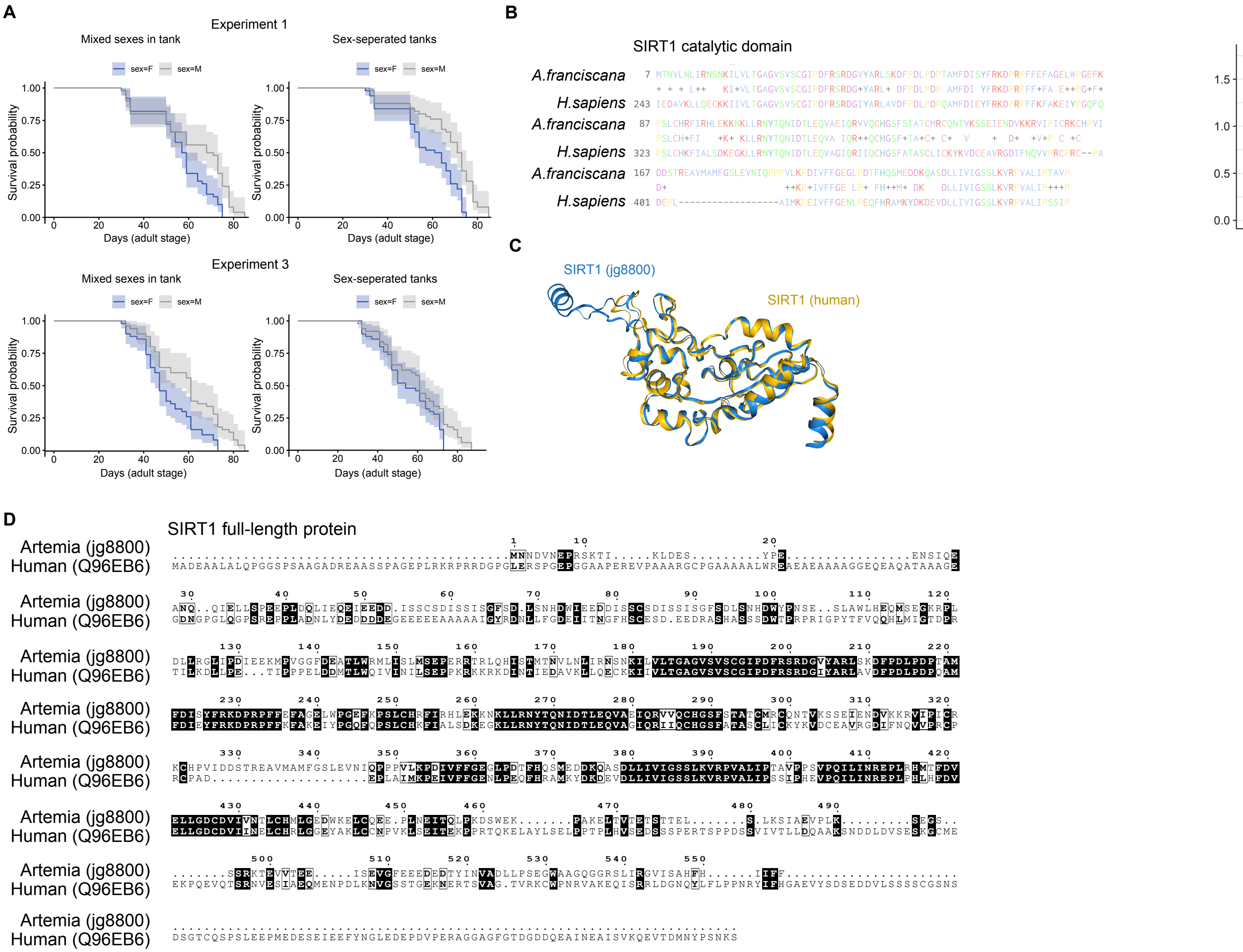
