## Supplementary material for "Convergent Evolution of H4K16ac-mediated Dosage Compensation Shapes Sex-dependent Lifespan in a ZW Species": S2_Data

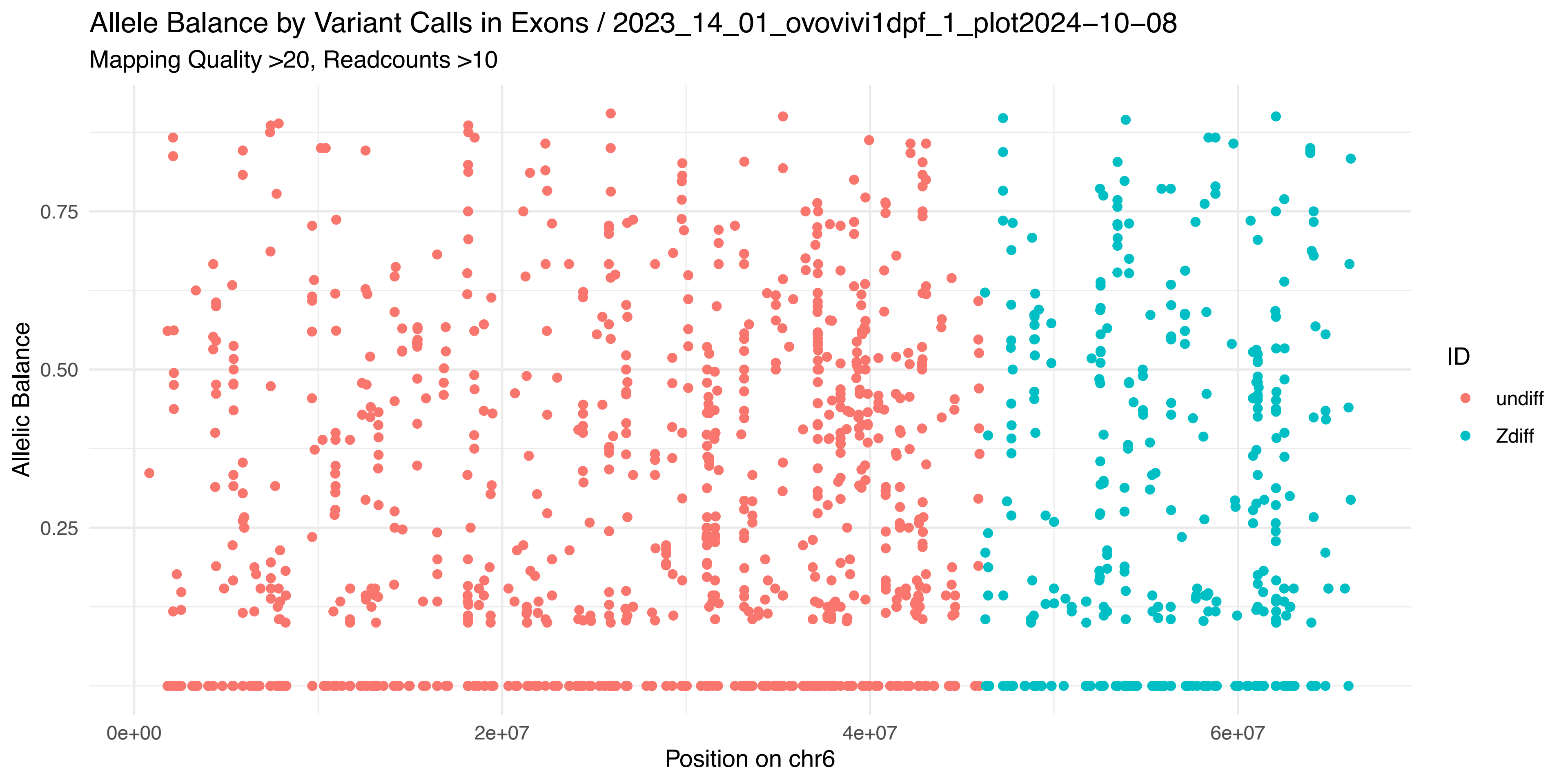
Allele Balance by Variant Calls in Exons / 2023_14_01_ovovivi1dpf_1_plot2024−10−08
Mapping Quality >20, Readcounts >10
Allelic Balance
0.75
0.50
0.25
0e+00
2e+07
4e+07
6e+07
Position on chr6
ID
undiff
Zdiff

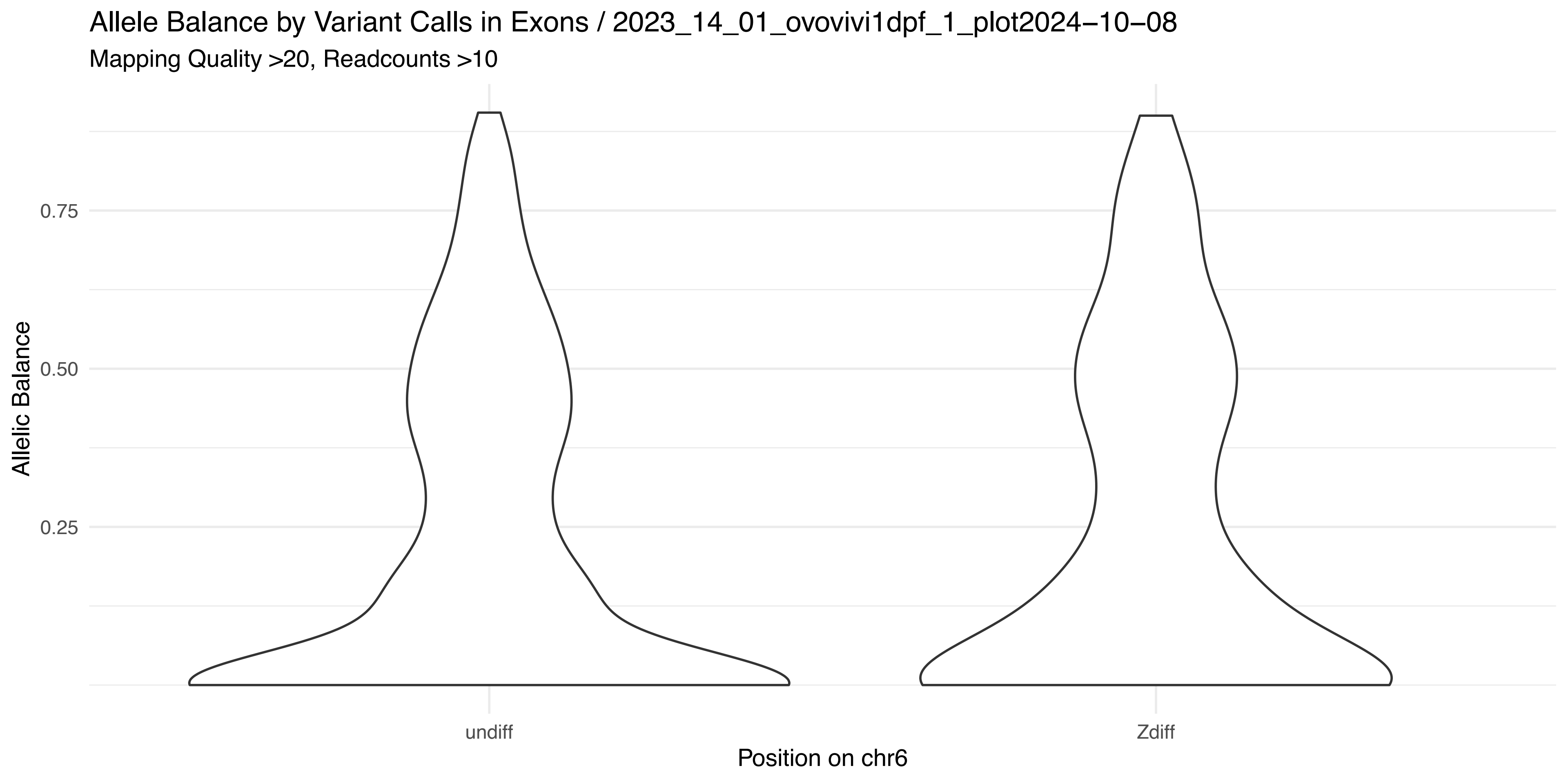
Allele Balance by Variant Calls in Exons / 2023_14_01_ovovivi1dpf_1_plot2024−10−08
Mapping Quality >20, Readcounts >10
Allelic Balance
0.75
0.50
0.25
undiff
Zdiff
Position on chr6

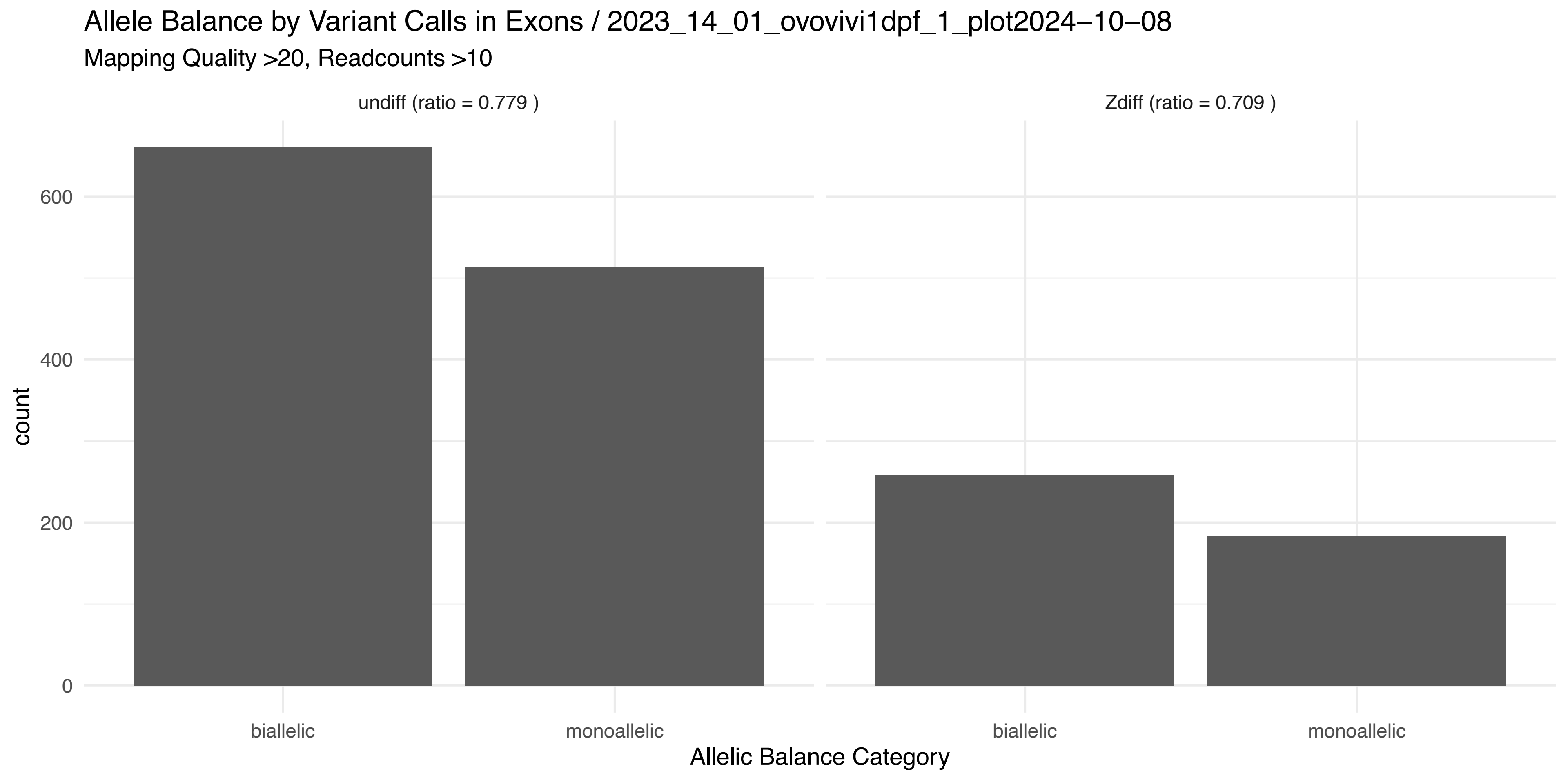
Allele Balance by Variant Calls in Exons / 2023_14_01_ovovivi1dpf_1_plot2024−10−08
Mapping Quality >20, Readcounts >10
undiff (ratio = 0.779 )
Zdiff (ratio = 0.709 )
count
600
400
200
0
biallelic
monoallelic
biallelic
monoallelic
Allelic Balance Category

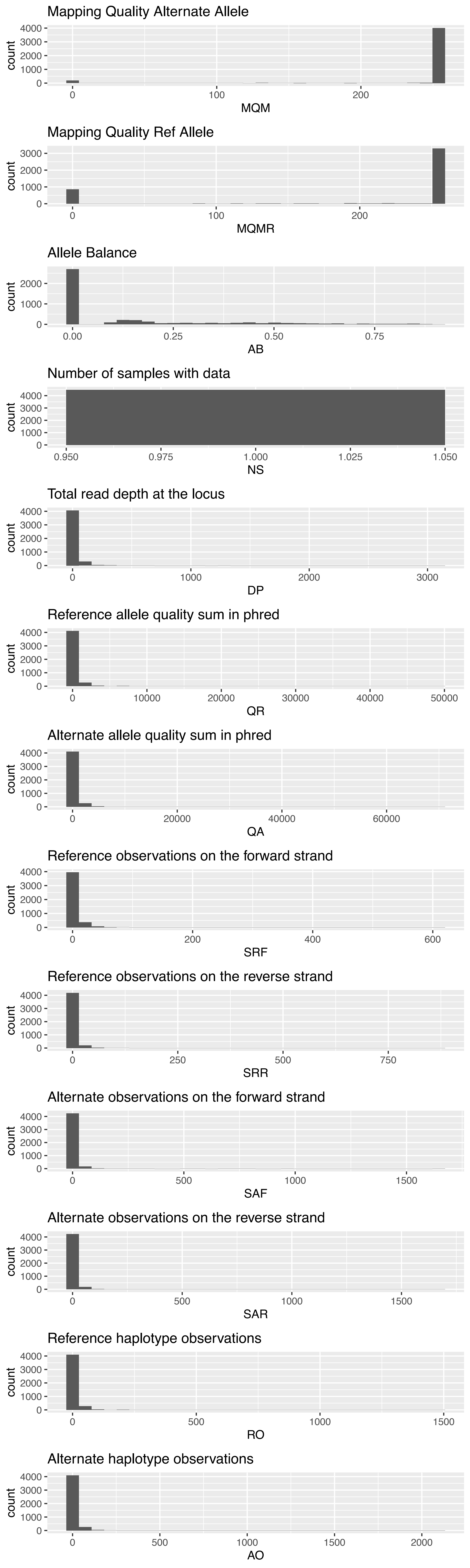
Mapping Quality Alternate Allele
count
MQM
Mapping Quality Ref Allele
count
MQMR
Allele Balance
count
AB
Number of samples with data
count
NS
Total read depth at the locus
count
DP
Reference allele quality sum in phred
count
QR
Alternate allele quality sum in phred
count
QA
Reference observations on the forward strand
count
SRF
Reference observations on the reverse strand
count
SRR
Alternate observations on the forward strand
count
SAF
Alternate observations on the reverse strand
count
SAR
Reference haplotype observations
count
RO
Alternate haplotype observations
count
AO

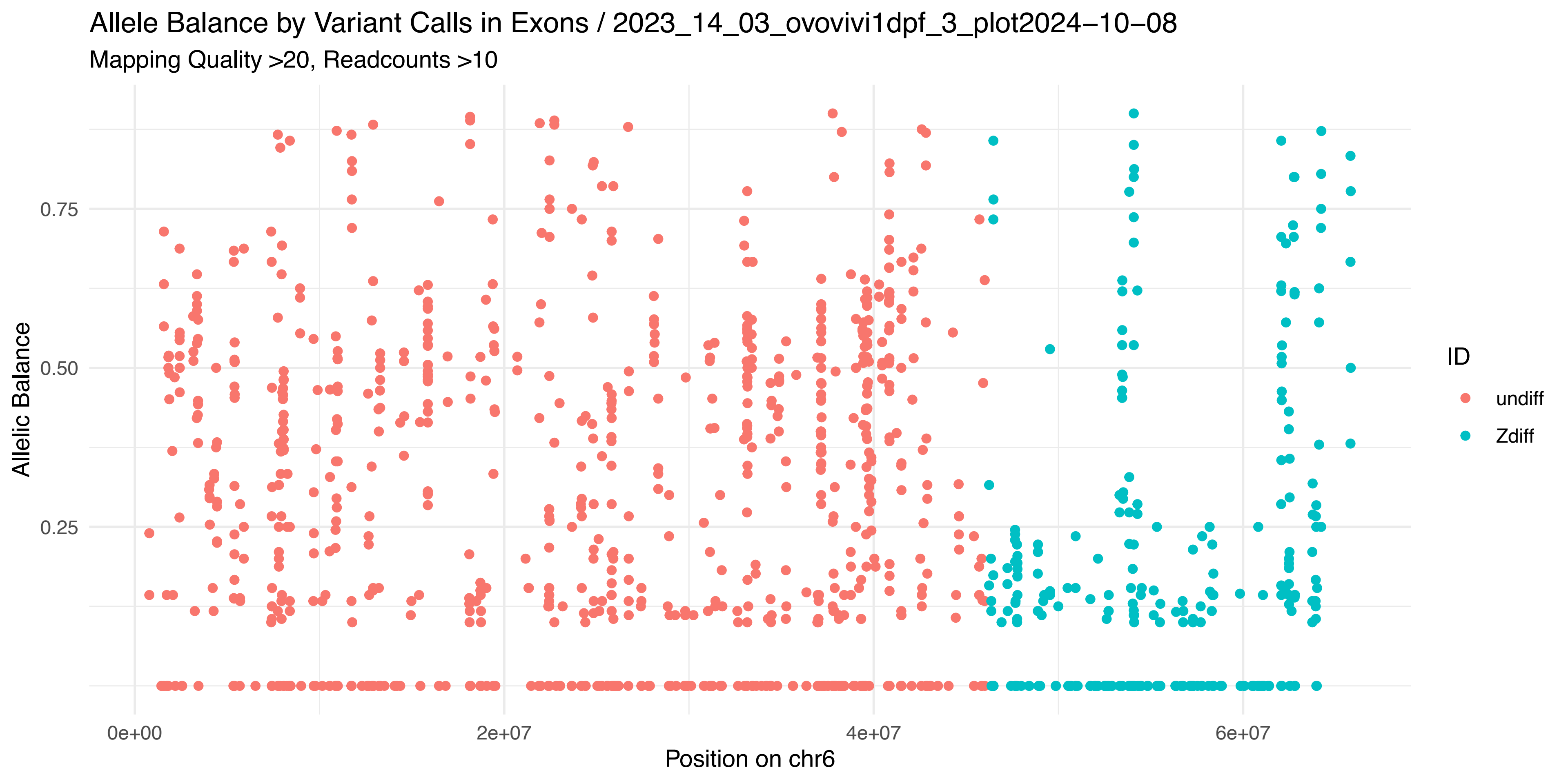

Allele Balance by Variant Calls in Exons / 2023_14_03_ovovivi1dpf_3_plot2024−10−08
Mapping Quality >20, Readcounts >10
Allelic Balance
0.75
0.50
0.25
0e+00
2e+07
4e+07
6e+07
Position on chr6
ID
undiff
Zdiff


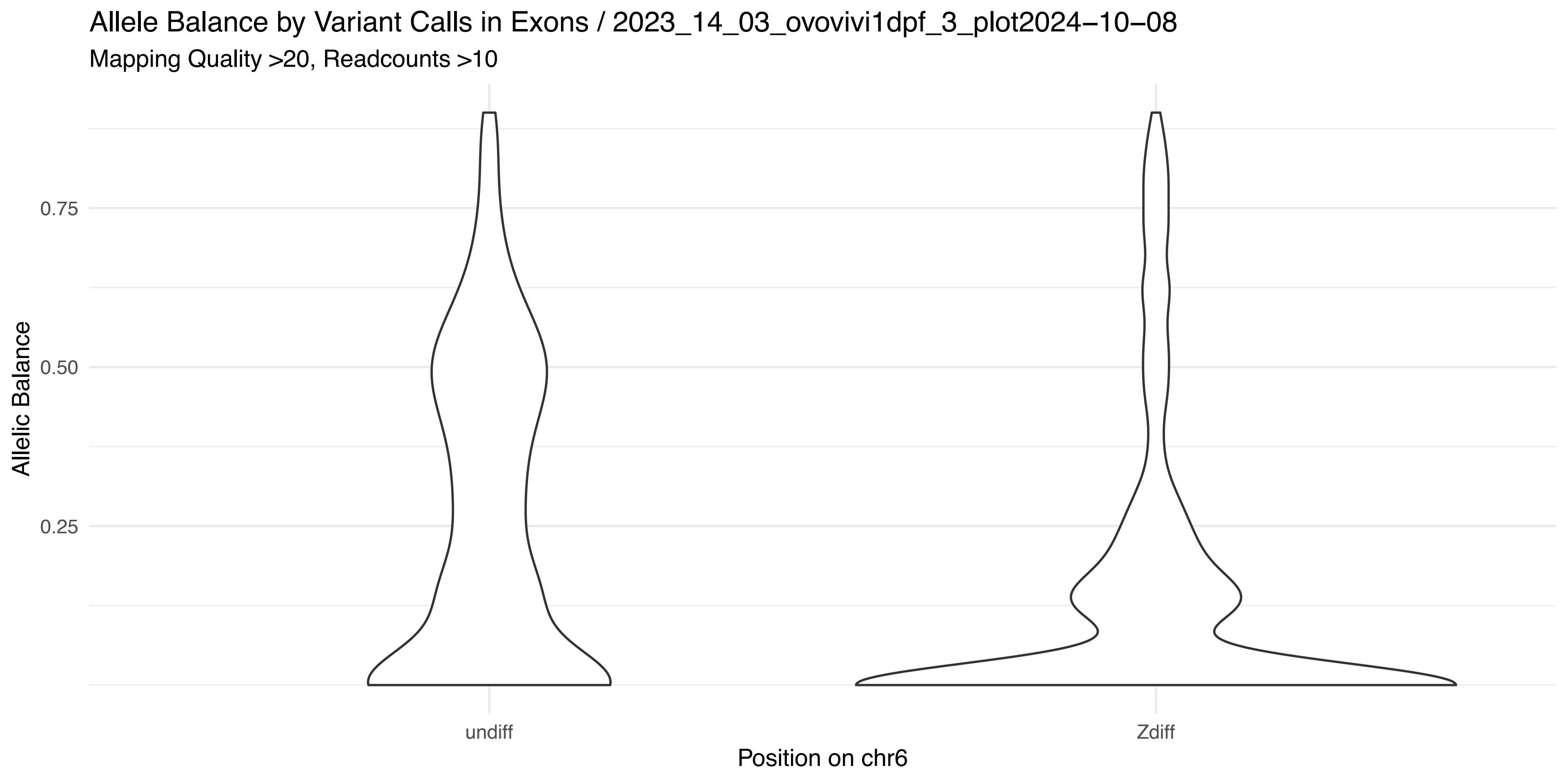

Allele Balance by Variant Calls in Exons / 2023_14_03_ovovivi1dpf_3_plot2024−10−08
Mapping Quality >20, Readcounts >10
Allelic Balance
0.75
0.50
0.25
undiff
Zdiff
Position on chr6


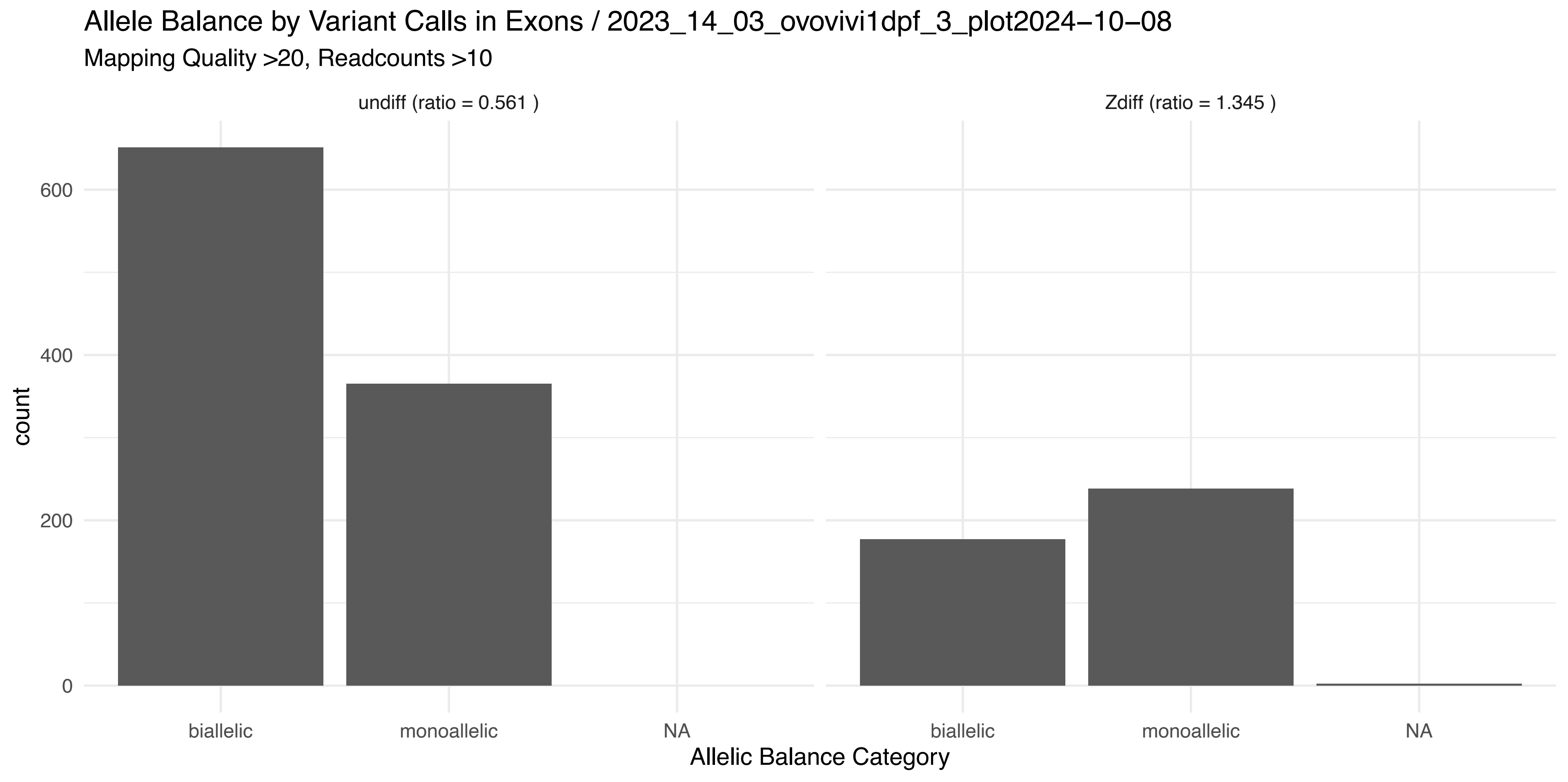

Allele Balance by Variant Calls in Exons / 2023_14_03_ovovivi1dpf_3_plot2024−10−08
Mapping Quality >20, Readcounts >10
undiff (ratio = 0.561 )
Zdiff (ratio = 1.345 )
count
600
400
200
0
biallelic
monoallelic
NA
Allelic Balance Category

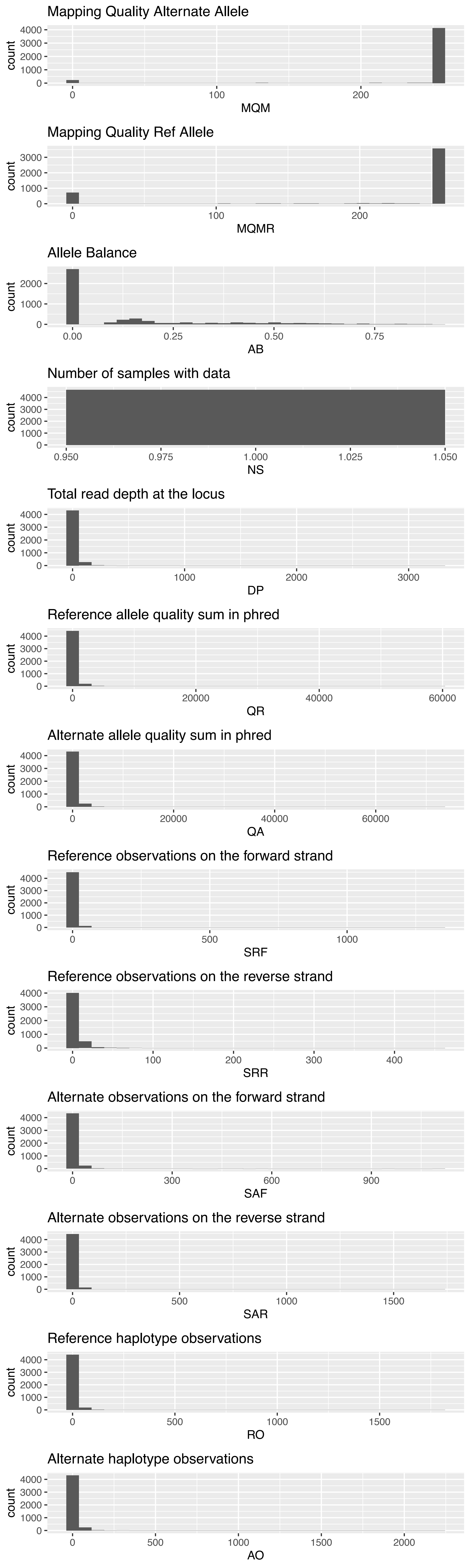

Mapping Quality Alternate Allele
count
0
1000
2000
3000
4000
0
100
200
MQM
Mapping Quality Ref Allele
count
0
1000
2000
3000
0
100
200
MQMR
Allele Balance
count
0
1000
2000
0.00
0.25
0.50
0.75
AB
Number of samples with data
count
0
1000
2000
3000
4000
0.950
0.975
1.000
1.025
1.050
NS
Total read depth at the locus
count
0
1000
2000
3000
4000
0
1000
2000
3000
DP
Reference allele quality sum in phred
count
0
1000
2000
3000
4000
0
20000
40000
60000
QR
Alternate allele quality sum in phred
count
0
1000
2000
3000
4000
0
20000
40000
60000
QA
Reference observations on the forward strand
count
0
1000
2000
3000
4000
0
500
1000
SRF
Reference observations on the reverse strand
count
0
1000
2000
3000
4000
0
100
200
300
400
SRR
Alternate observations on the forward strand
count
0
1000
2000
3000
4000
0
300
600
900
SAF
Alternate observations on the reverse strand
count
0
1000
2000
3000
4000
0
500
1000
1500
SAR
Reference haplotype observations
count
0
1000
2000
3000
4000
0
500
1000
1500
RO
Alternate haplotype observations
count
0
1000
2000
3000
4000
0
500
1000
1500
2000
AO

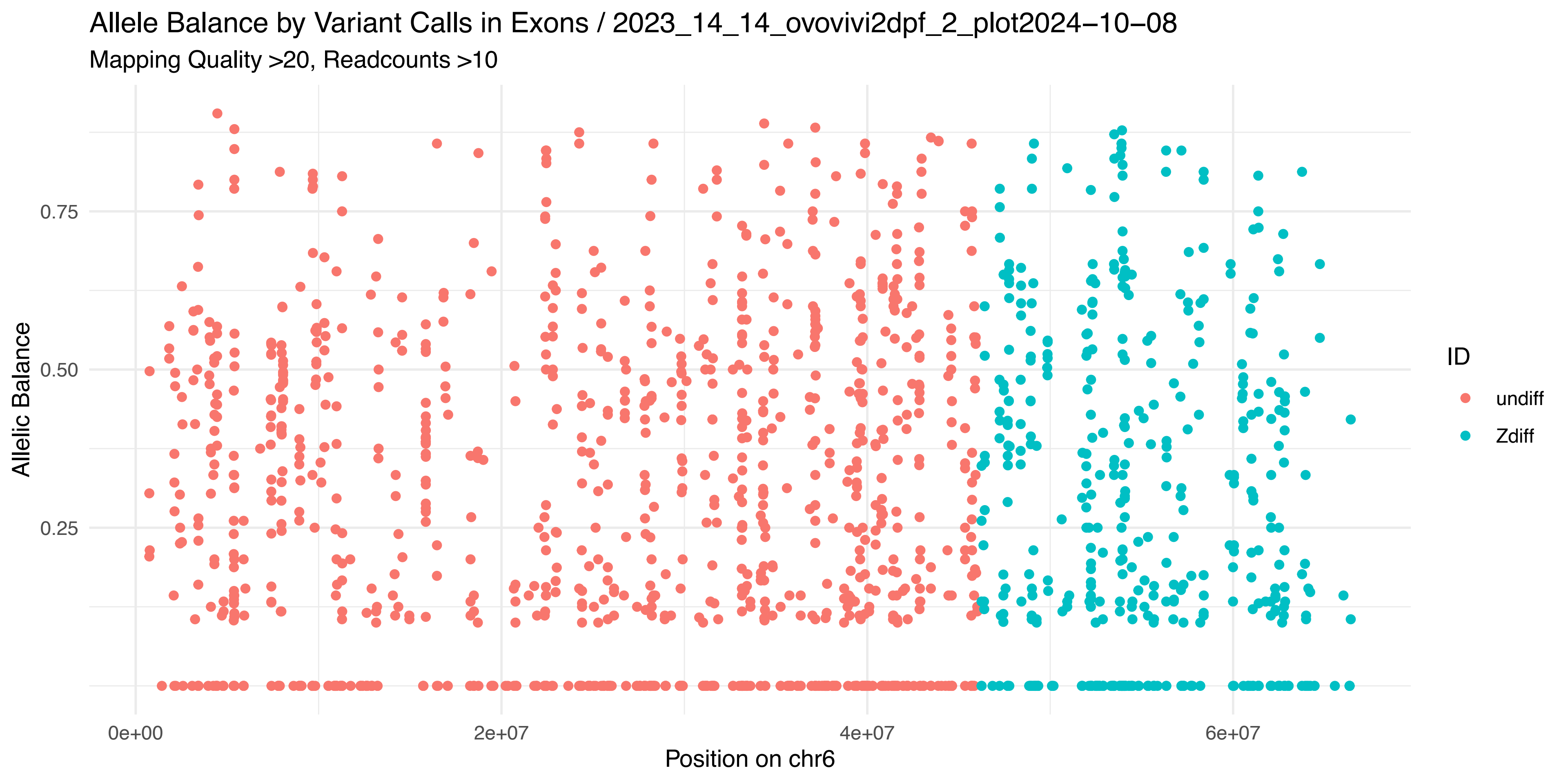
Allele Balance by Variant Calls in Exons / 2023_14_14_ovovivi2dpf_2_plot2024−10−08
Mapping Quality >20, Readcounts >10
Allelic Balance
0.75
0.50
0.25
0e+00
2e+07
4e+07
6e+07
Position on chr6
ID
undiff
Zdiff

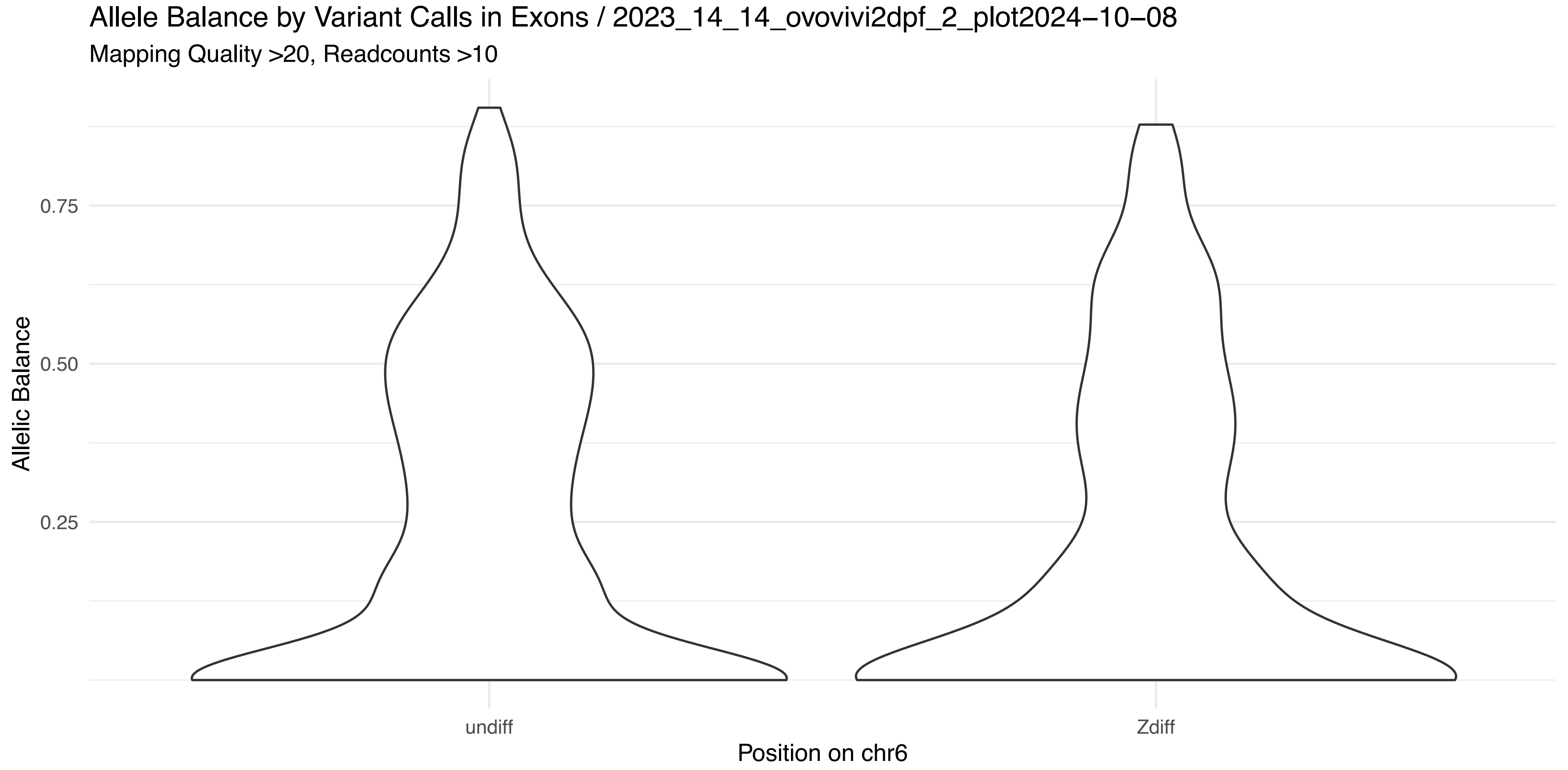
Allele Balance by Variant Calls in Exons / 2023_14_14_ovovivi2dpf_2_plot2024−10−08
Mapping Quality >20, Readcounts >10
Allelic Balance
0.75
0.50
0.25
undiff
Zdiff
Position on chr6

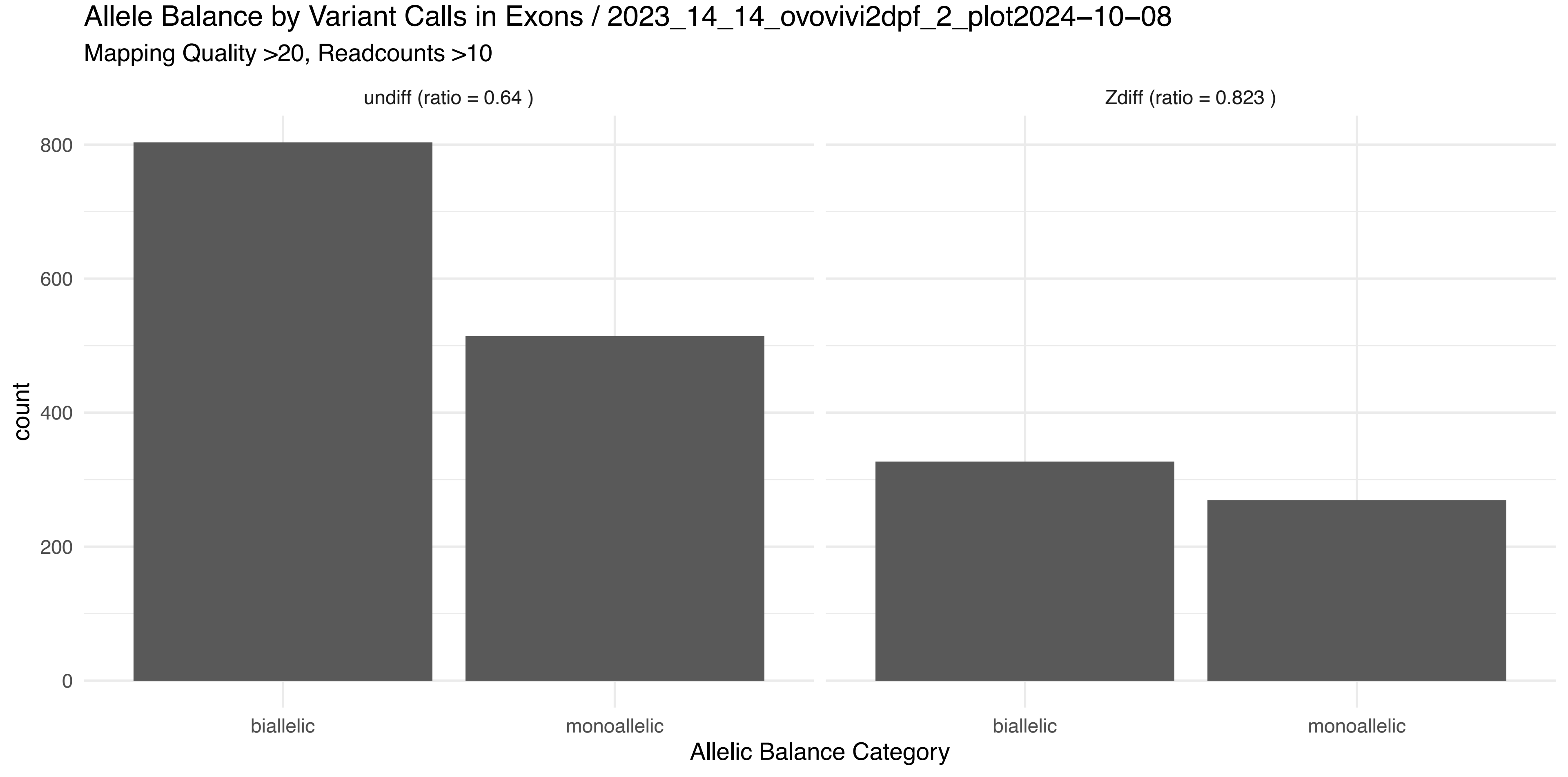
Allele Balance by Variant Calls in Exons / 2023_14_14_ovovivi2dpf_2_plot2024−10−08
Mapping Quality >20, Readcounts >10
undiff (ratio = 0.64 )
Zdiff (ratio = 0.823 )
count
800
600
400
200
0
biallelic
monoallelic
biallelic
monoallelic
Allelic Balance Category

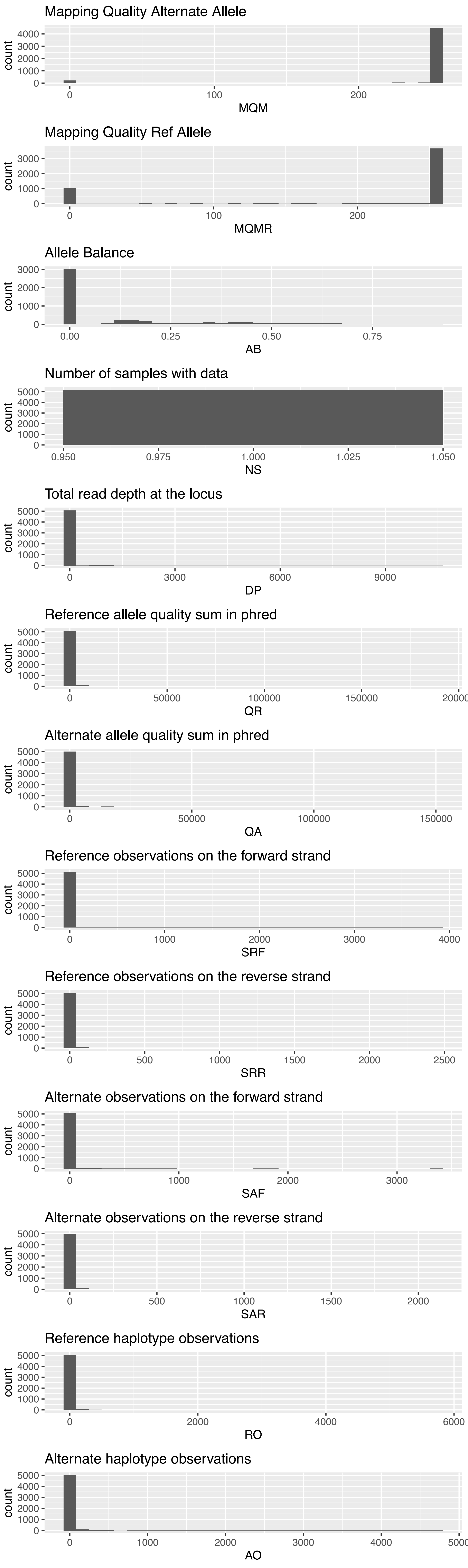
Mapping Quality Alternate Allele
count
0
1000
2000
3000
4000
0
100
200
MQM
Mapping Quality Ref Allele
count
0
1000
2000
3000
0
100
200
MQMR
Allele Balance
count
0
1000
2000
3000
0.00
0.25
0.50
0.75
AB
Number of samples with data
count
0
1000
2000
3000
4000
5000
0.950
0.975
1.000
1.025
1.050
NS
Total read depth at the locus
count
0
1000
2000
3000
4000
5000
0
3000
6000
9000
DP
Reference allele quality sum in phred
count
0
1000
2000
3000
4000
5000
0
50000
100000
150000
20000
QR
Alternate allele quality sum in phred
count
0
1000
2000
3000
4000
5000
0
50000
100000
150000
QA
Reference observations on the forward strand
count
0
1000
2000
3000
4000
5000
0
1000
2000
3000
4000
SRF
Reference observations on the reverse strand
count
0
1000
2000
3000
4000
5000
0
500
1000
1500
2000
2500
SRR
Alternate observations on the forward strand
count
0
1000
2000
3000
4000
5000
0
1000
2000
3000
SAF
Alternate observations on the reverse strand
count
0
1000
2000
3000
4000
5000
0
500
1000
1500
2000
SAR
Reference haplotype observations
count
0
1000
2000
3000
4000
5000
0
2000
4000
6000
RO
Alternate haplotype observations
count
0
1000
2000
3000
4000
5000
0
1000
2000
3000
4000
5000
AO

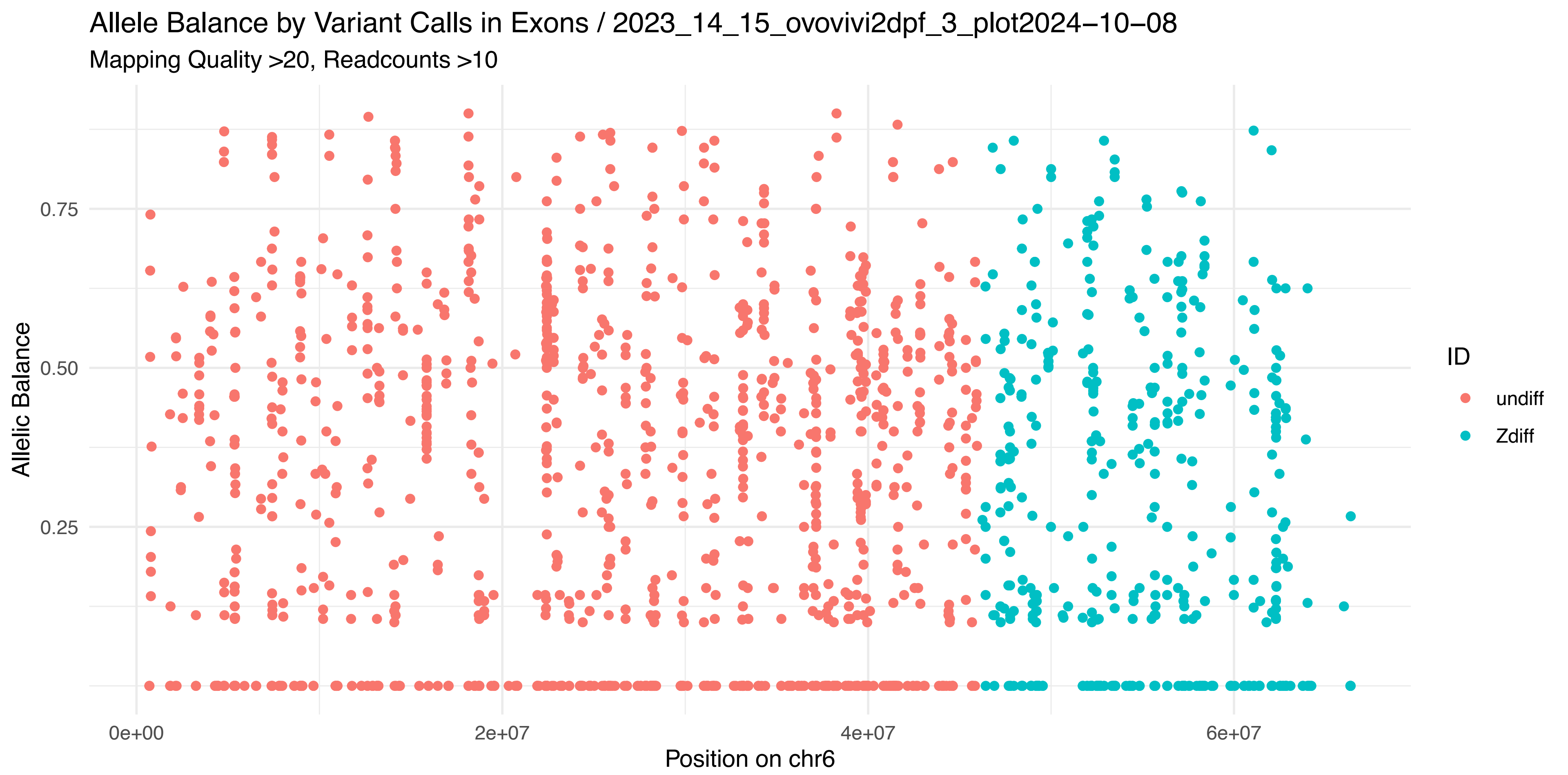
Allele Balance by Variant Calls in Exons / 2023_14_15_ovovivi2dpf_3_plot2024−10−08
Mapping Quality >20, Readcounts >10
Allelic Balance
0.75
0.50
0.25
0e+00
2e+07
4e+07
6e+07
Position on chr6
ID
undiff
Zdiff

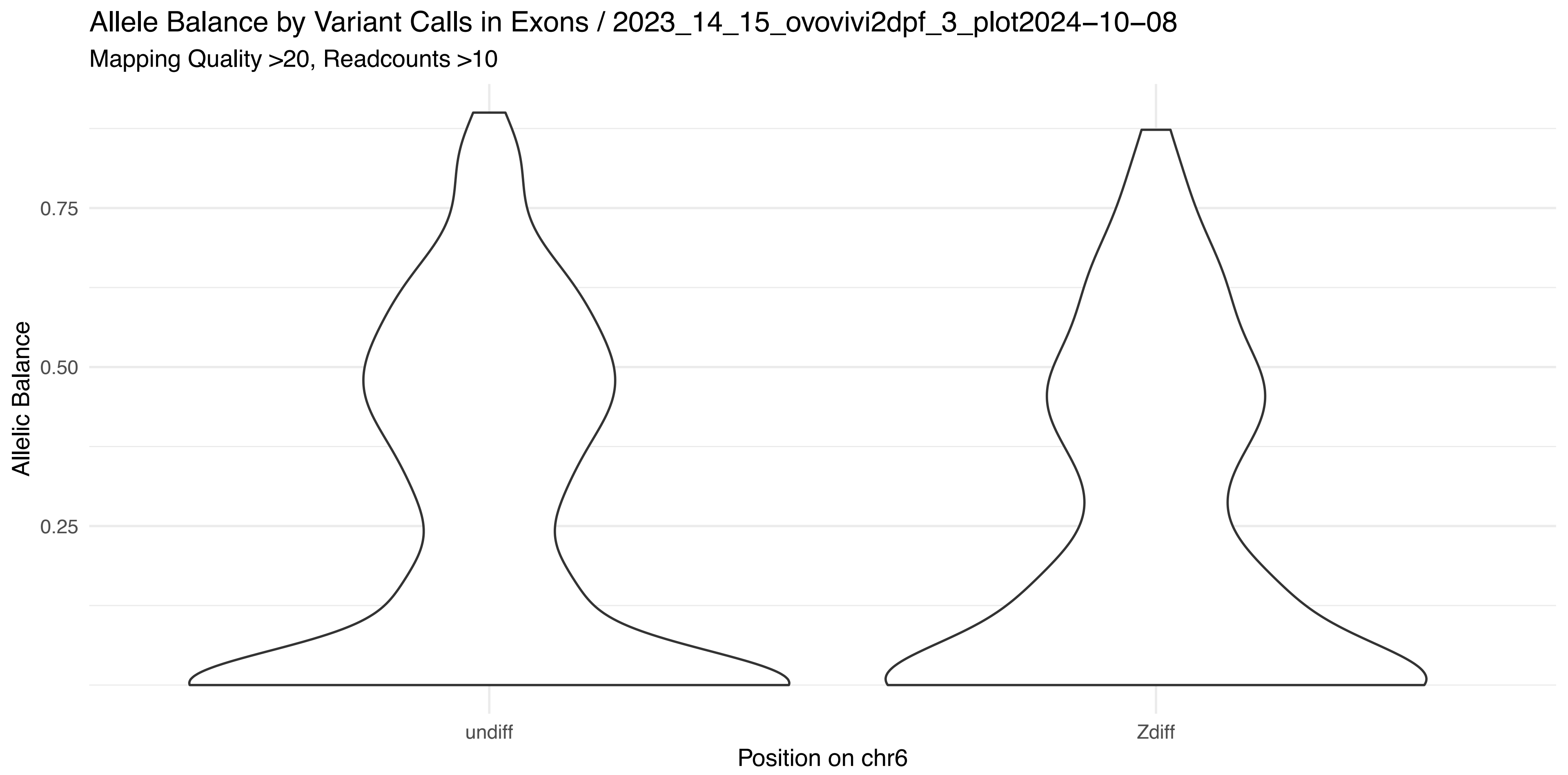
Allele Balance by Variant Calls in Exons / 2023_14_15_ovovivi2dpf_3_plot2024−10−08
Mapping Quality >20, Readcounts >10
Allelic Balance
0.75
0.50
0.25
undiff
Zdiff
Position on chr6

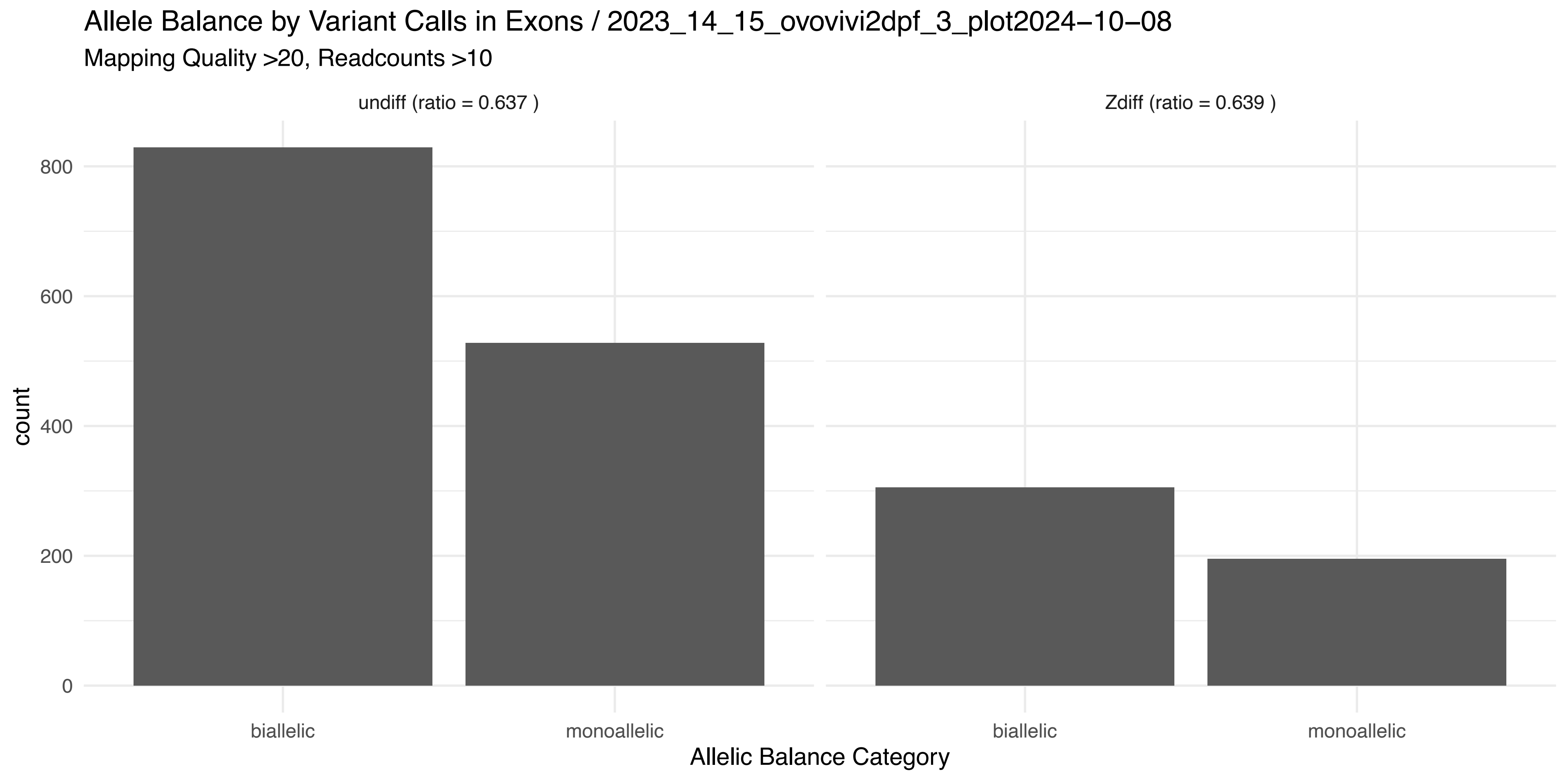
Allele Balance by Variant Calls in Exons / 2023_14_15_ovovivi2dpf_3_plot2024−10−08
Mapping Quality >20, Readcounts >10
undiff (ratio = 0.637 )
Zdiff (ratio = 0.639 )
count
800
600
400
200
0
biallelic
monoallelic
biallelic
monoallelic
Allelic Balance Category

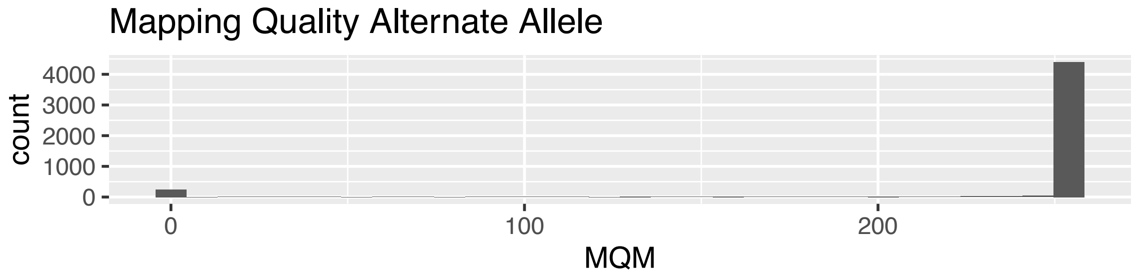
Mapping Quality Alternate Allele
count
4000
3000
2000
1000
0
0
100
200
MQM

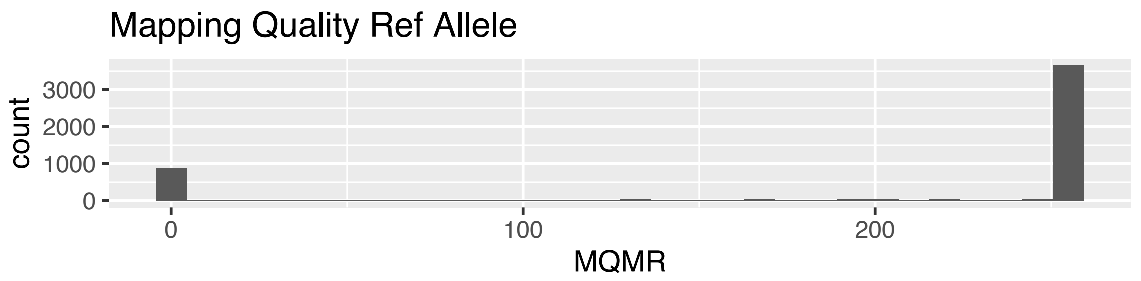
Mapping Quality Ref Allele
count
3000
2000
1000
0
0
100
200
MQMR

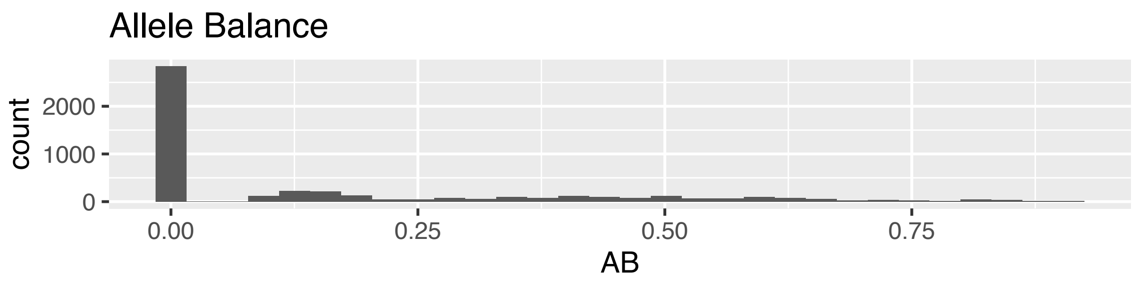
Allele Balance
count
2000
1000
0
0.00
0.25
0.50
0.75
AB

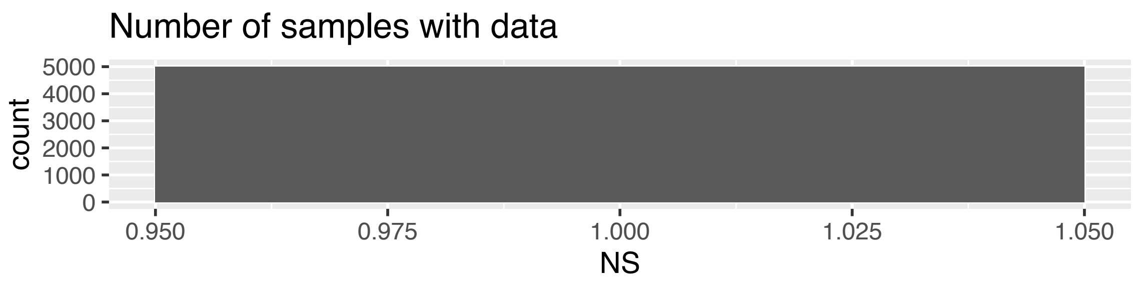
Number of samples with data
count
5000
4000
3000
2000
1000
0
0.950
0.975
1.000
1.025
1.050
NS

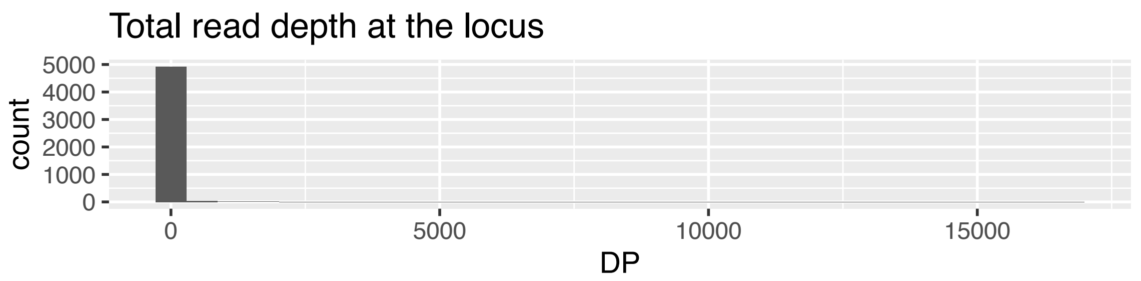
Total read depth at the locus
count
5000
4000
3000
2000
1000
0
0
5000
10000
15000
DP

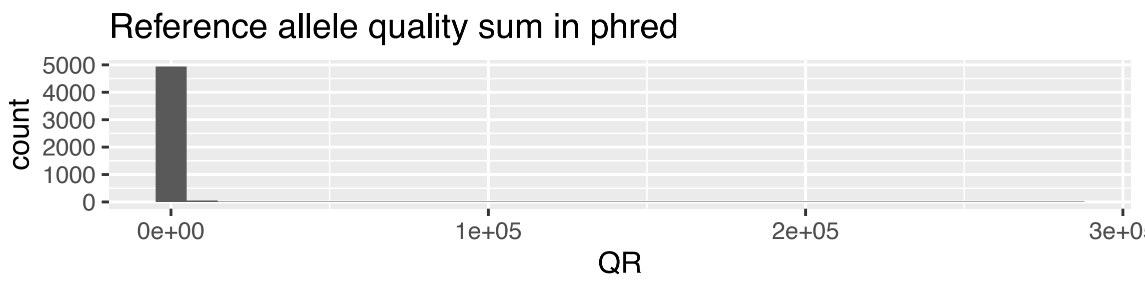
Reference allele quality sum in phred
count
5000
4000
3000
2000
1000
0
0e+00
1e+05
2e+05
QR

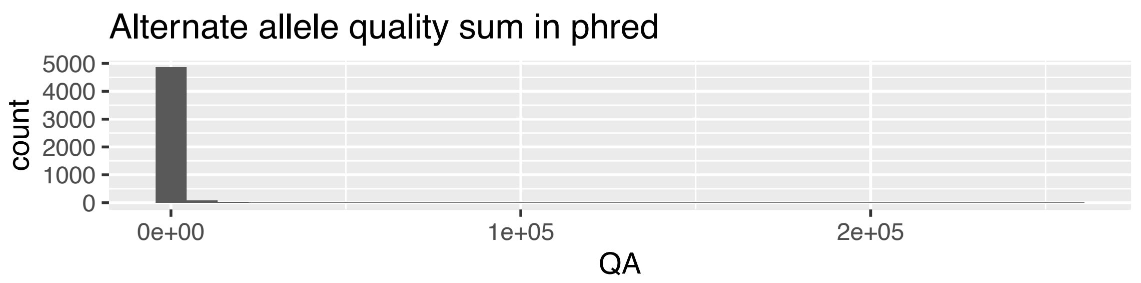
Alternate allele quality sum in phred
count
5000
4000
3000
2000
1000
0
0e+00
1e+05
2e+05
QA

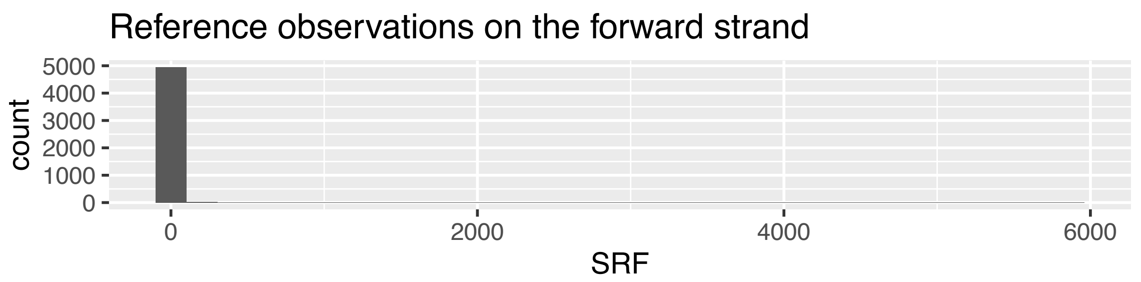
Reference observations on the forward strand
count
5000
4000
3000
2000
1000
0
0
2000
4000
6000
SRF

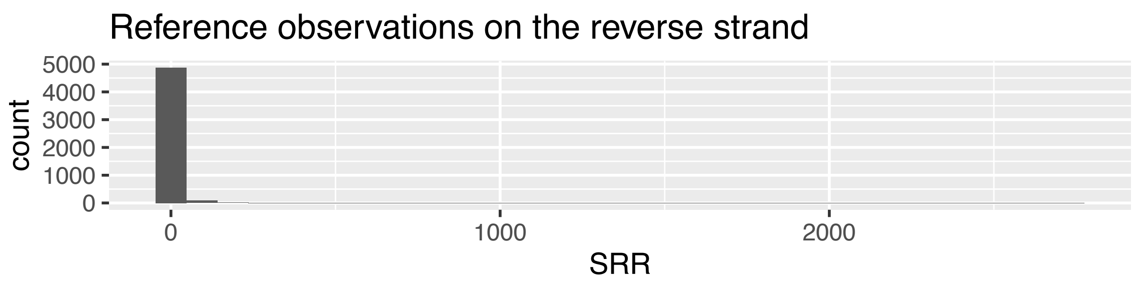
Reference observations on the reverse strand
count
5000
4000
3000
2000
1000
0
0
1000
2000
SRR

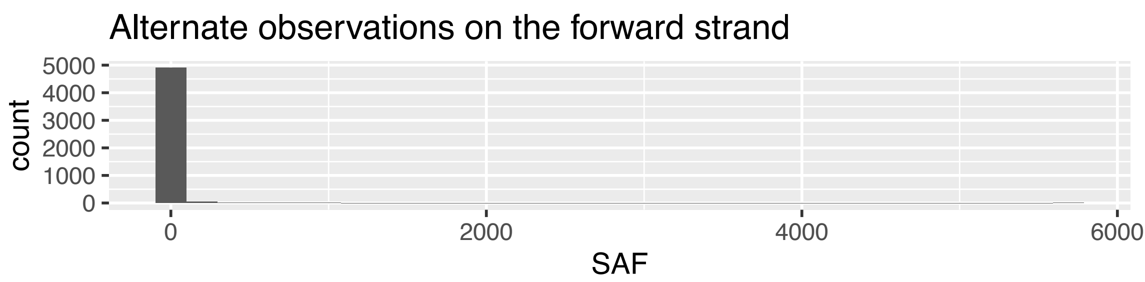
Alternate observations on the forward strand
count
5000
4000
3000
2000
1000
0
0
2000
4000
6000
SAF

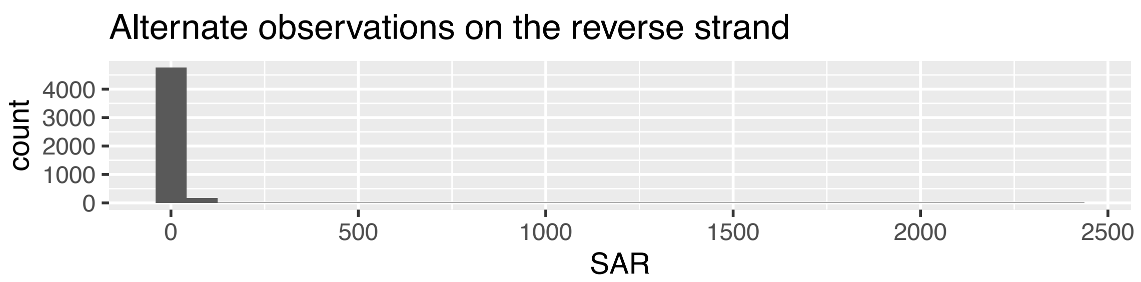
Alternate observations on the reverse strand
count
4000
3000
2000
1000
0
0
500
1000
1500
2000
2500
SAR

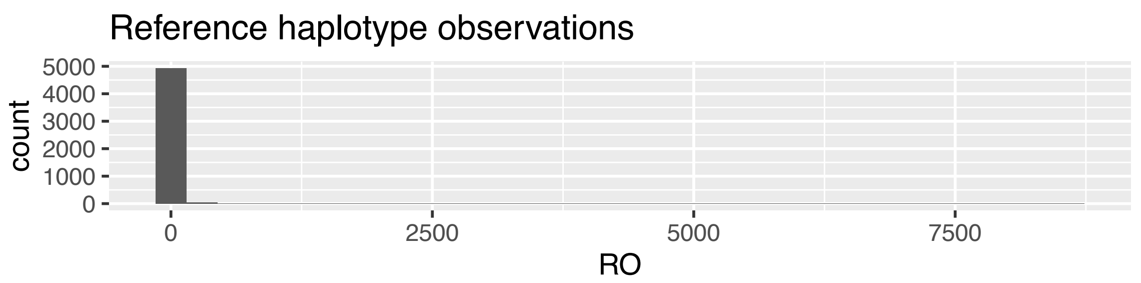
Reference haplotype observations
count
5000
4000
3000
2000
1000
0
0
2500
5000
7500
RO

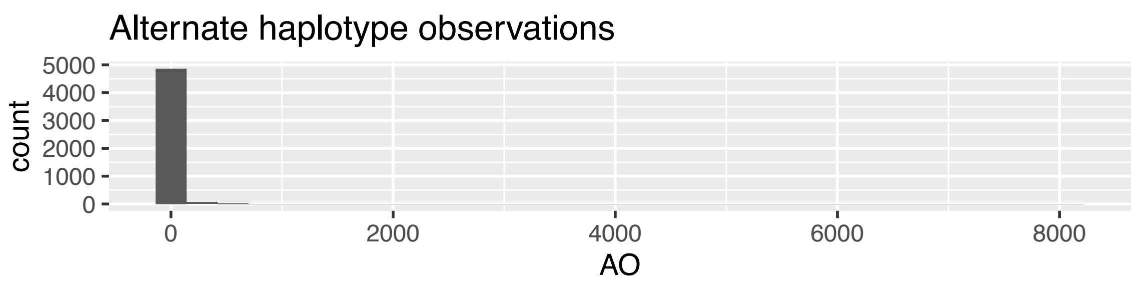
Alternate haplotype observations
count
5000
4000
3000
2000
1000
0
0
2000
4000
6000
8000
AO

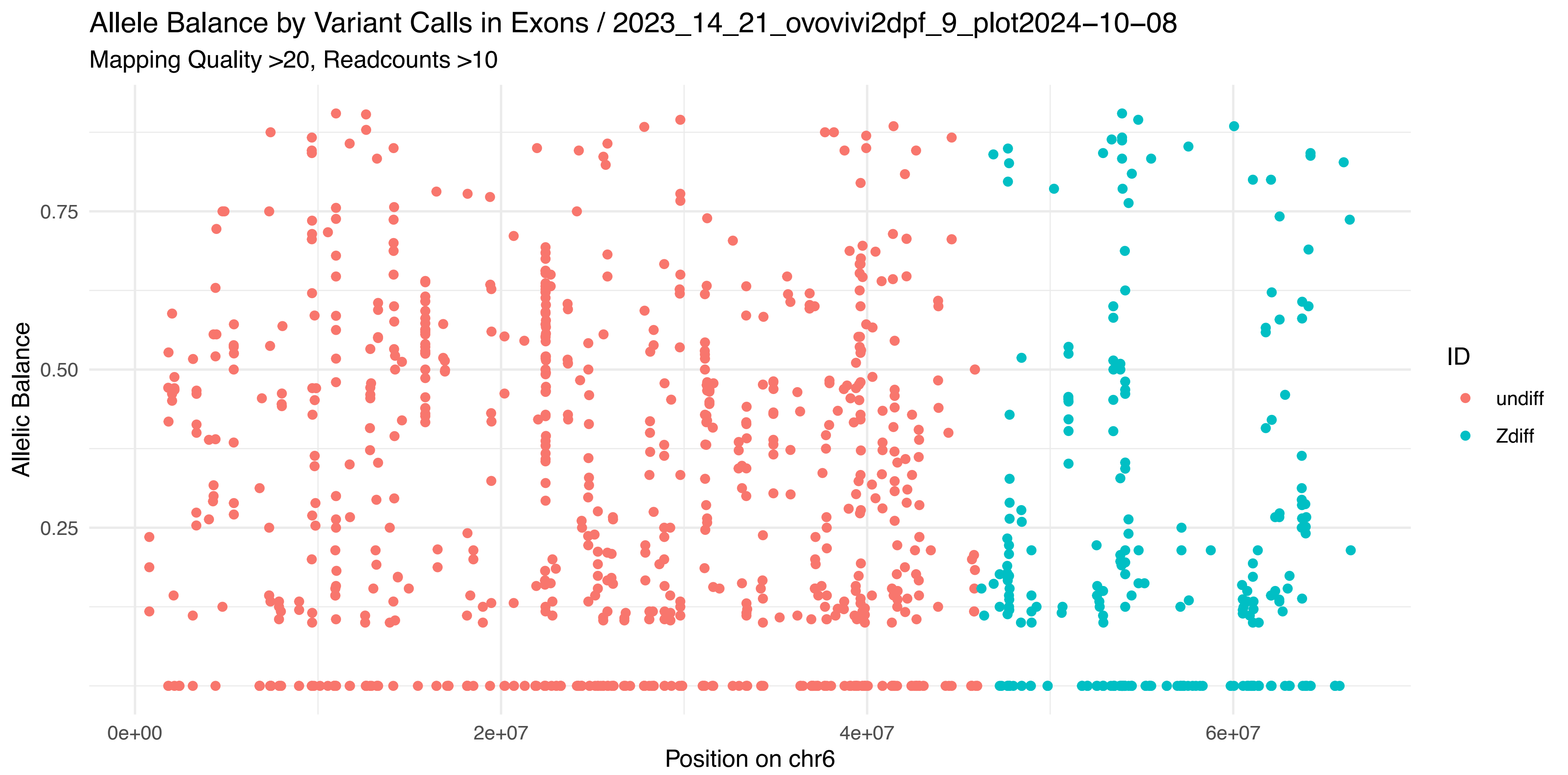

Allele Balance by Variant Calls in Exons / 2023_14_21_ovovivi2dpf_9_plot2024−10−08
Mapping Quality >20, Readcounts >10
Allelic Balance
0.75
0.50
0.25
0e+00
2e+07
4e+07
6e+07
Position on chr6
ID
undiff
Zdiff


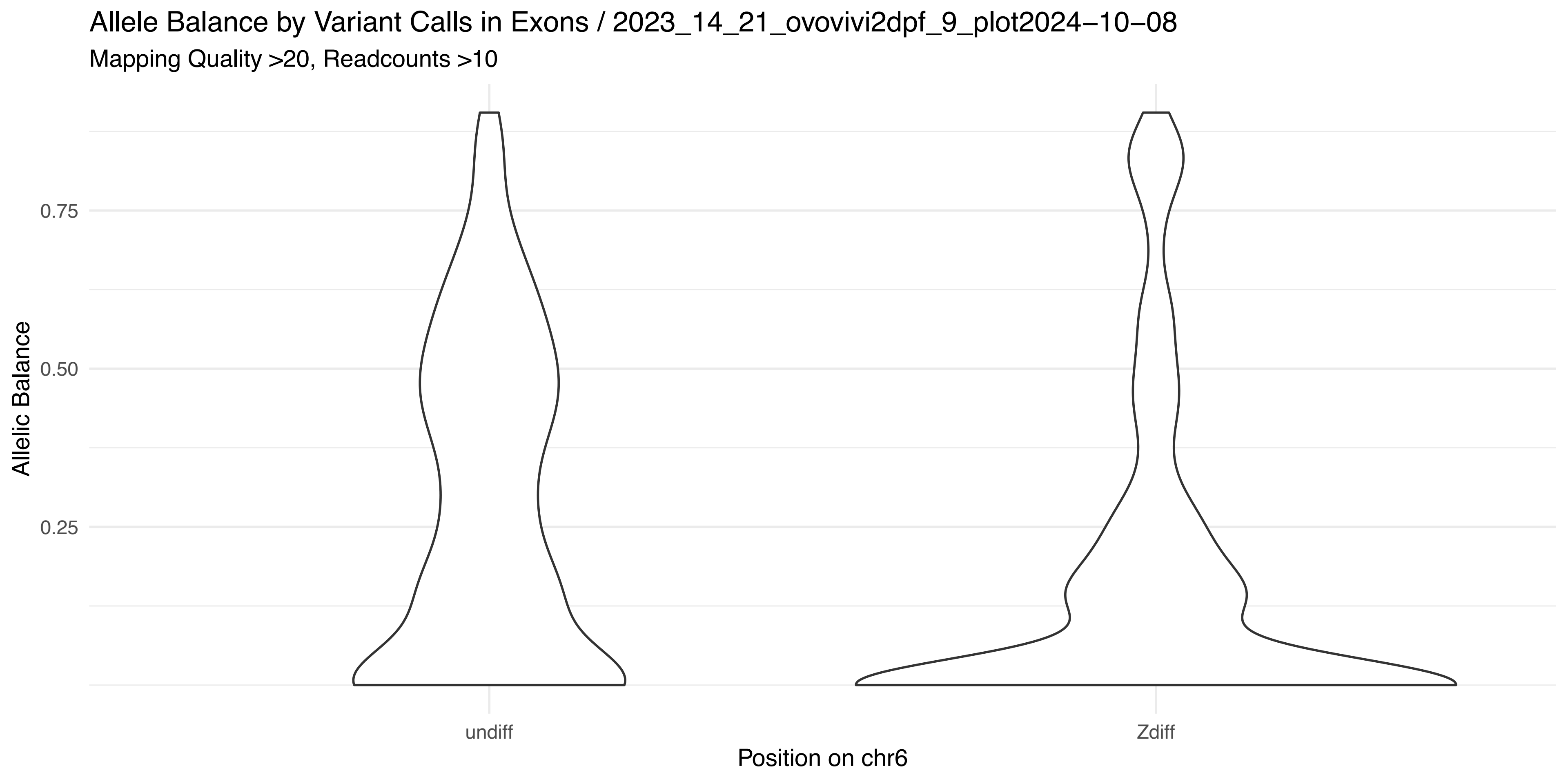

Allele Balance by Variant Calls in Exons / 2023_14_21_ovovivi2dpf_9_plot2024−10−08
Mapping Quality >20, Readcounts >10
Allelic Balance
0.75
0.50
0.25
undiff
Zdiff
Position on chr6


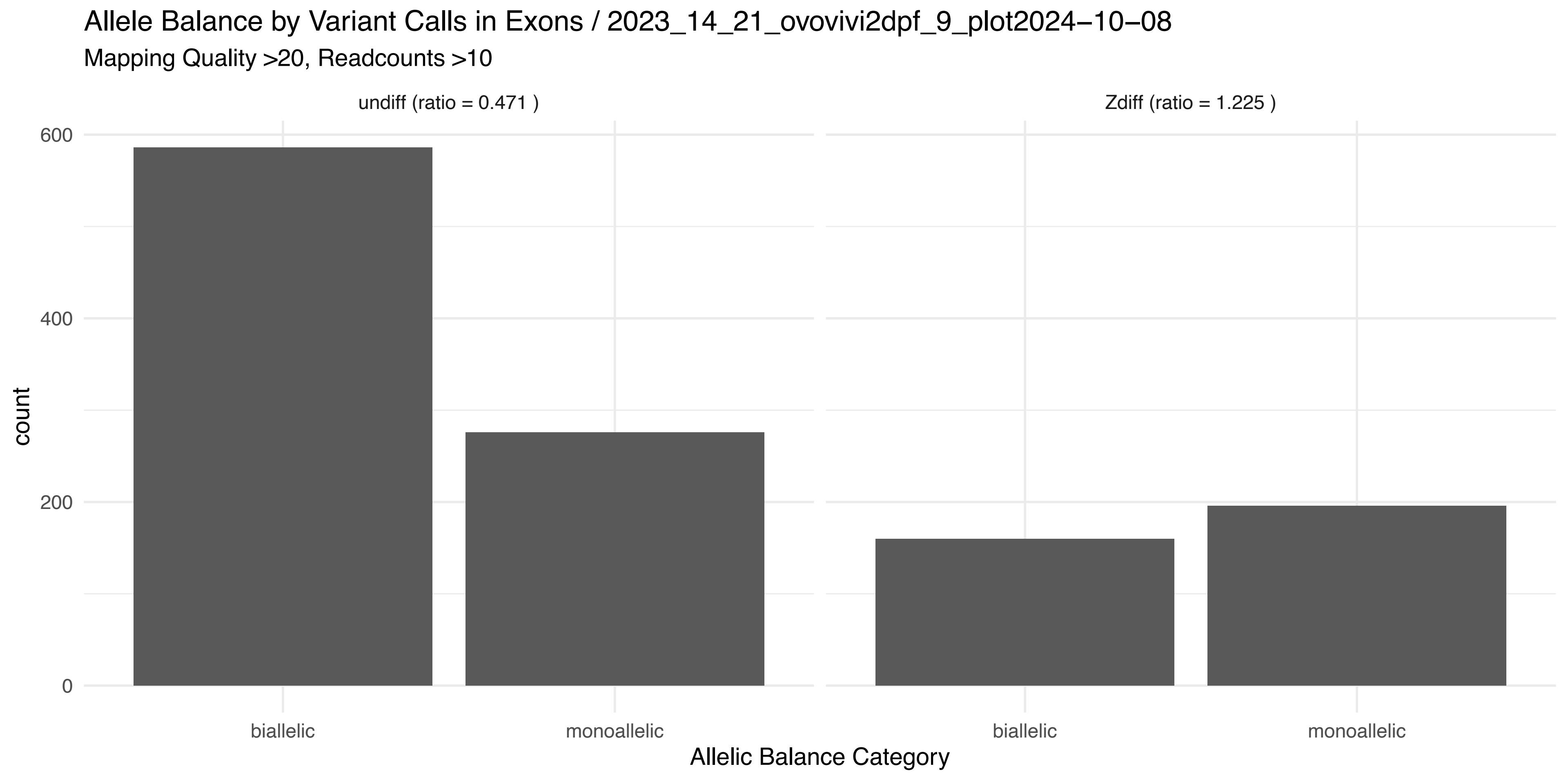

Allele Balance by Variant Calls in Exons / 2023_14_21_ovovivi2dpf_9_plot2024−10−08
Mapping Quality >20, Readcounts >10
undiff (ratio = 0.471 )
Zdiff (ratio = 1.225 )
count
600
400
200
0
biallelic
monoallelic
biallelic
monoallelic
Allelic Balance Category

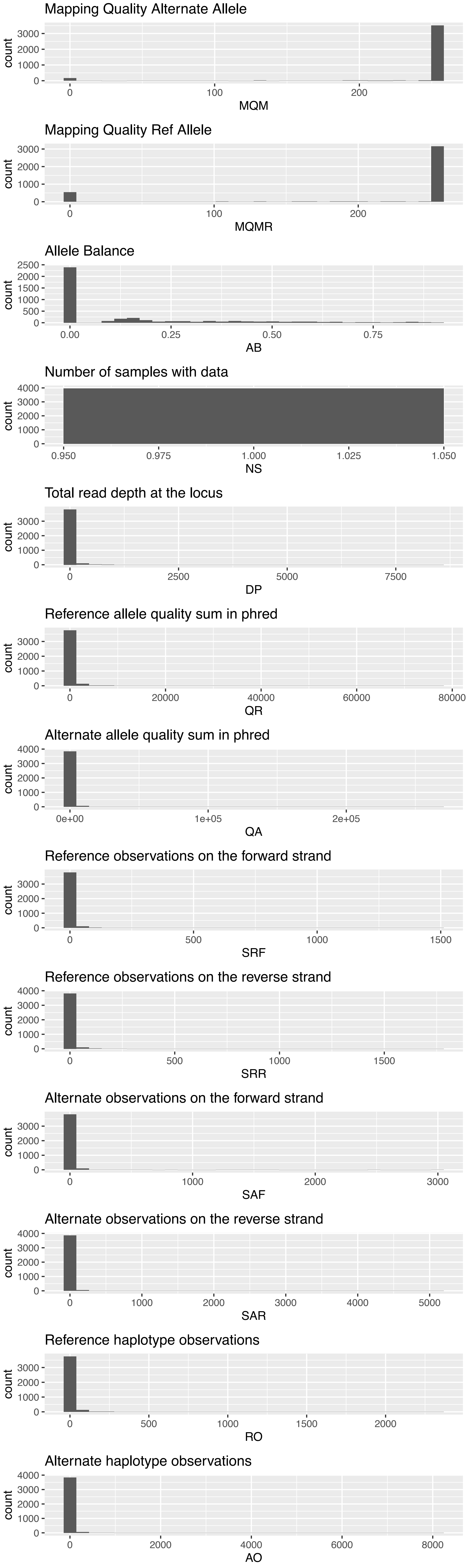
Mapping Quality Alternate Allele
count
0
1000
2000
3000
0
100
200
MQM
Mapping Quality Ref Allele
count
MQMR
Allele Balance
0
500
1000
1500
2000
2500
0.00
0.25
0.50
0.75
AB
Number of samples with data
4000
0.950
0.975
1.000
1.025
1.050
NS
Total read depth at the locus
2500
5000
7500
DP
Reference allele quality sum in phred
20000
40000
60000
80000
QR
Alternate allele quality sum in phred
0e+00
1e+05
2e+05
QA
Reference observations on the forward strand
500
1000
1500
SRF
Reference observations on the reverse strand
SRR
Alternate observations on the forward strand
3000
SAF
Alternate observations on the reverse strand
4000
5000
SAR
Reference haplotype observations
RO
Alternate haplotype observations
6000
8000
AO

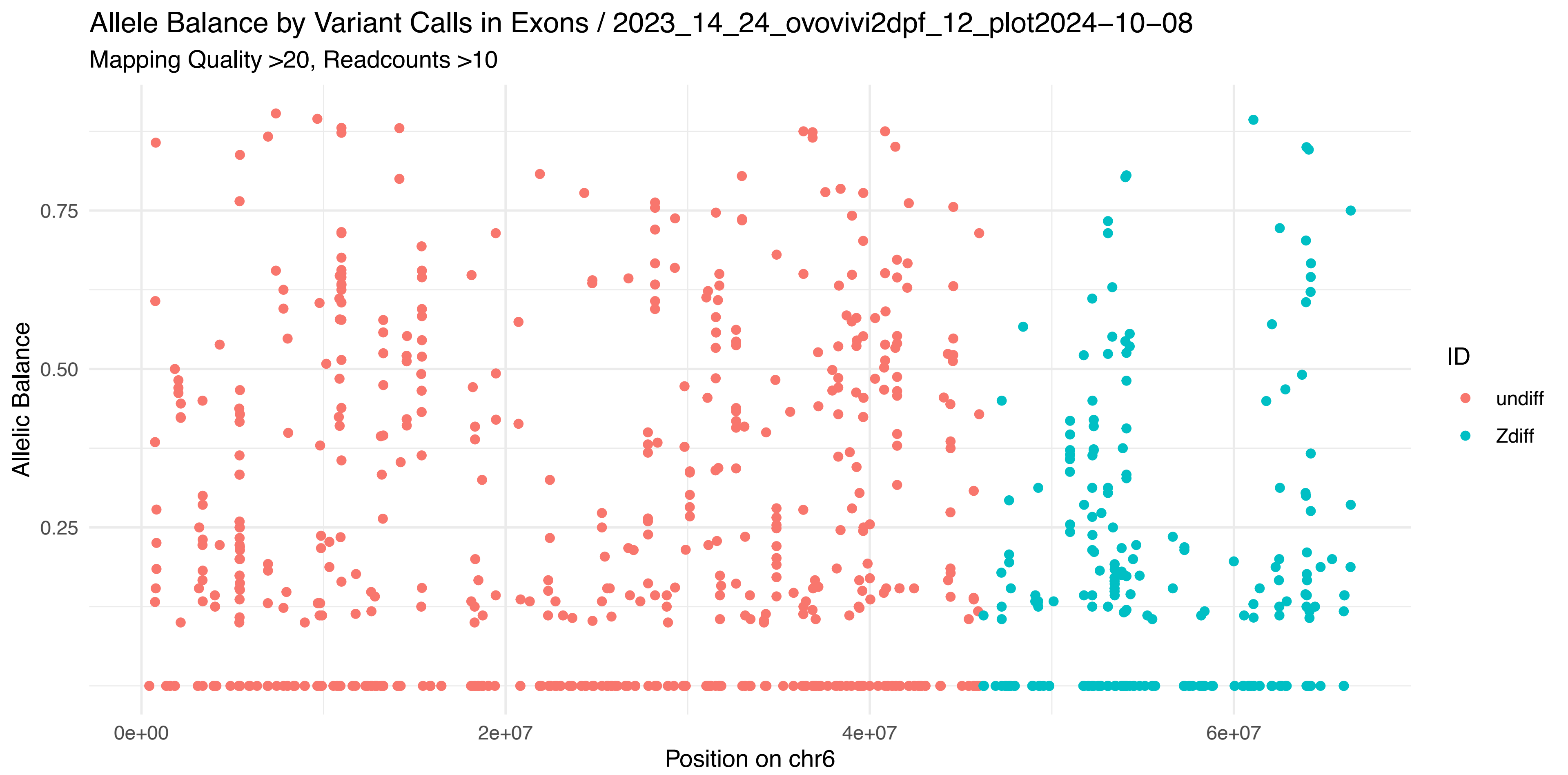
Allele Balance by Variant Calls in Exons / 2023_14_24_ovovivi2dpf_12_plot2024−10−08
Mapping Quality >20, Readcounts >10
Allelic Balance
0.75
0.50
0.25
0e+00
2e+07
4e+07
6e+07
Position on chr6
ID
undiff
Zdiff

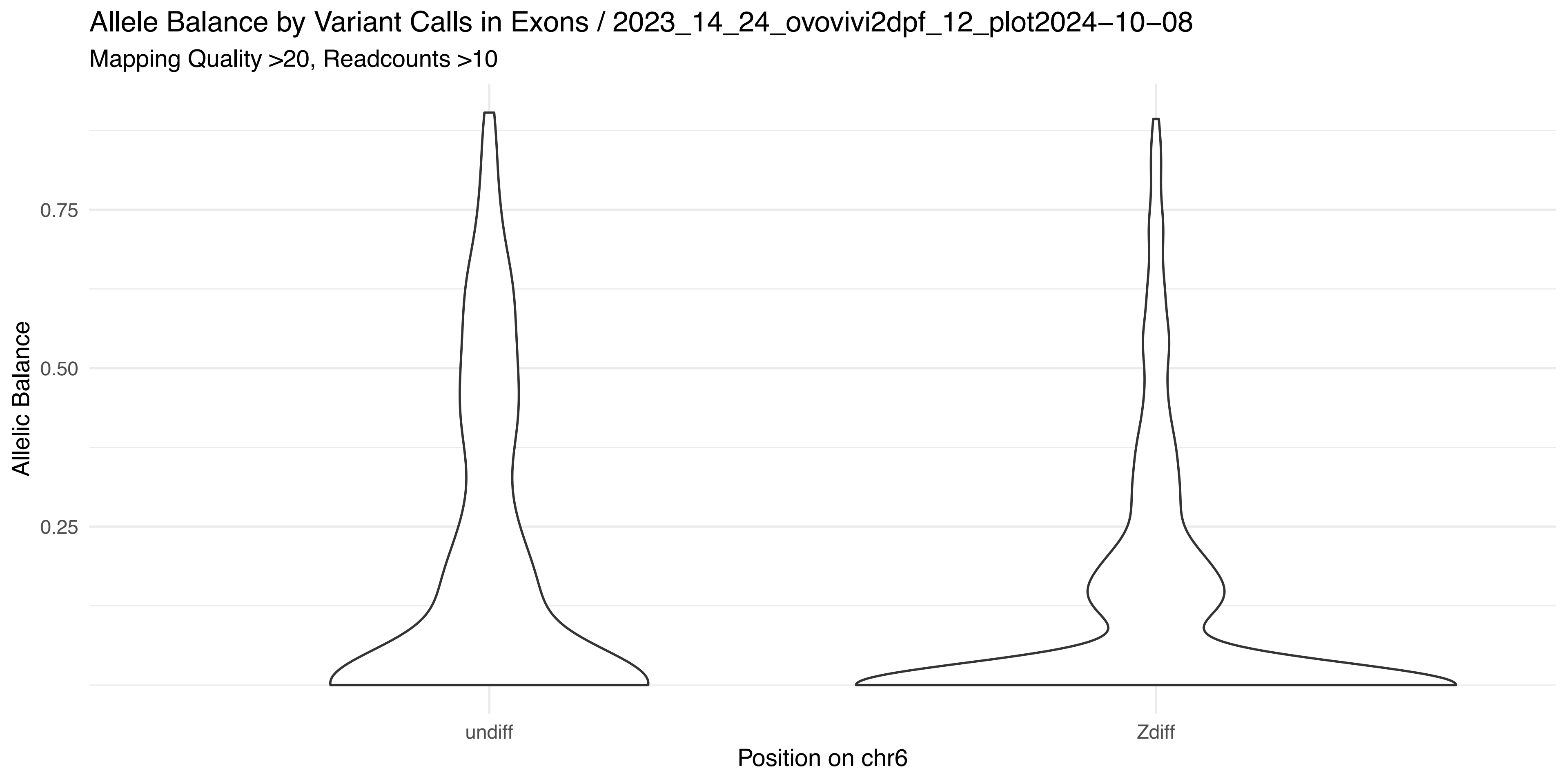
Allele Balance by Variant Calls in Exons / 2023_14_24_ovovivi2dpf_12_plot2024−10−08
Mapping Quality >20, Readcounts >10
Allelic Balance
0.75
0.50
0.25
undiff
Zdiff
Position on chr6

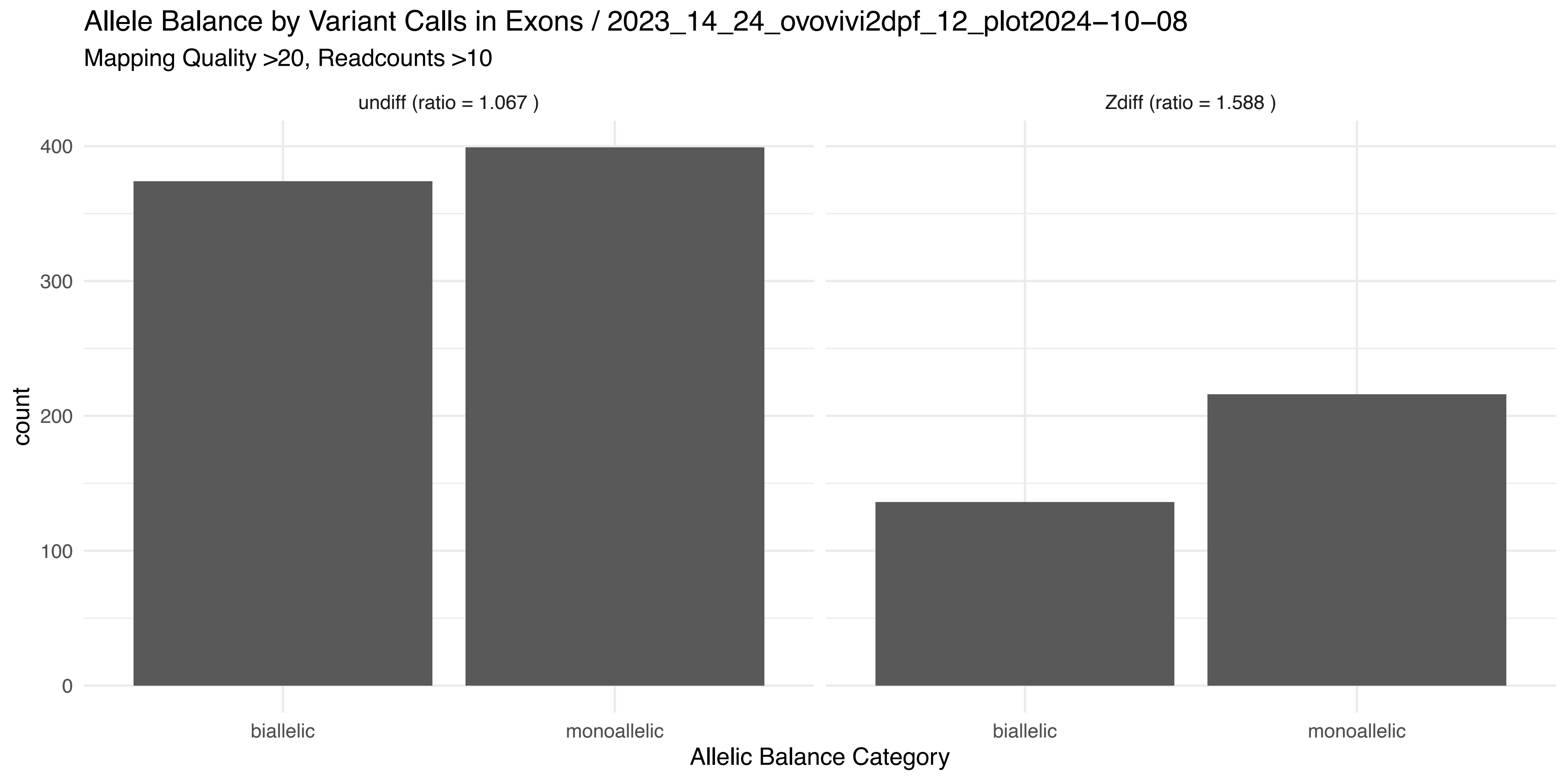
Allele Balance by Variant Calls in Exons / 2023_14_24_ovovivi2dpf_12_plot2024−10−08
Mapping Quality >20, Readcounts >10
undiff (ratio = 1.067 )
Zdiff (ratio = 1.588 )
count
400
300
200
100
0
biallelic
monoallelic
biallelic
monoallelic
Allelic Balance Category

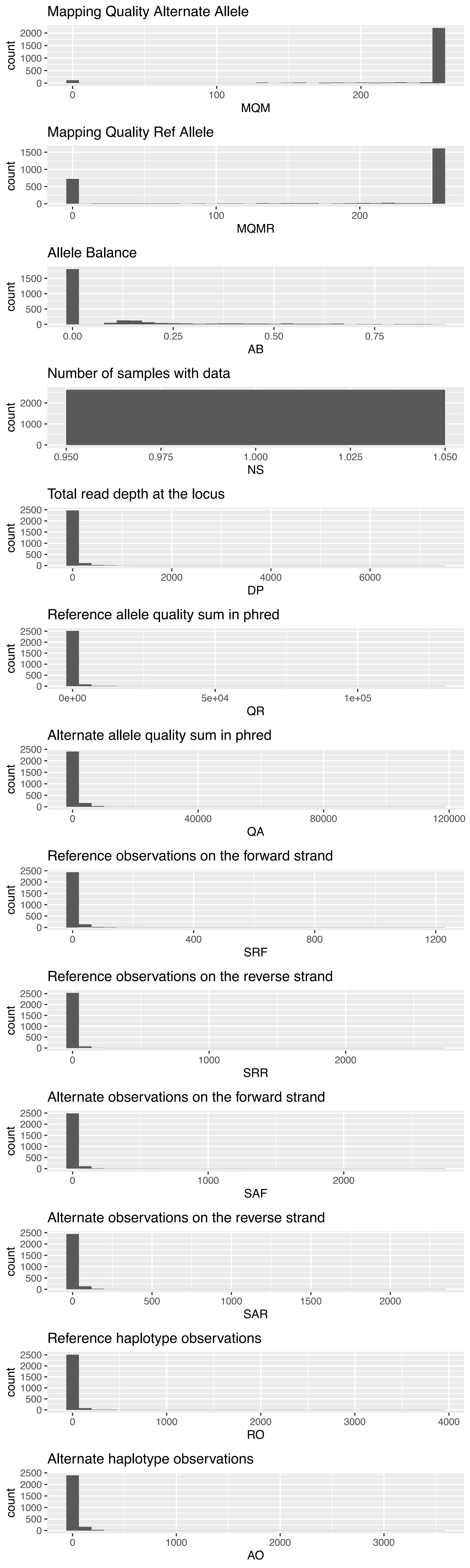
Mapping Quality Alternate Allele
count
MQM
Mapping Quality Ref Allele
count
MQMR
Allele Balance
count
AB
Number of samples with data
count
NS
Total read depth at the locus
count
DP
Reference allele quality sum in phred
count
QR
Alternate allele quality sum in phred
count
QA
Reference observations on the forward strand
count
SRF
Reference observations on the reverse strand
count
SRR
Alternate observations on the forward strand
count
SAF
Alternate observations on the reverse strand
count
SAR
Reference haplotype observations
count
RO
Alternate haplotype observations
count
AO

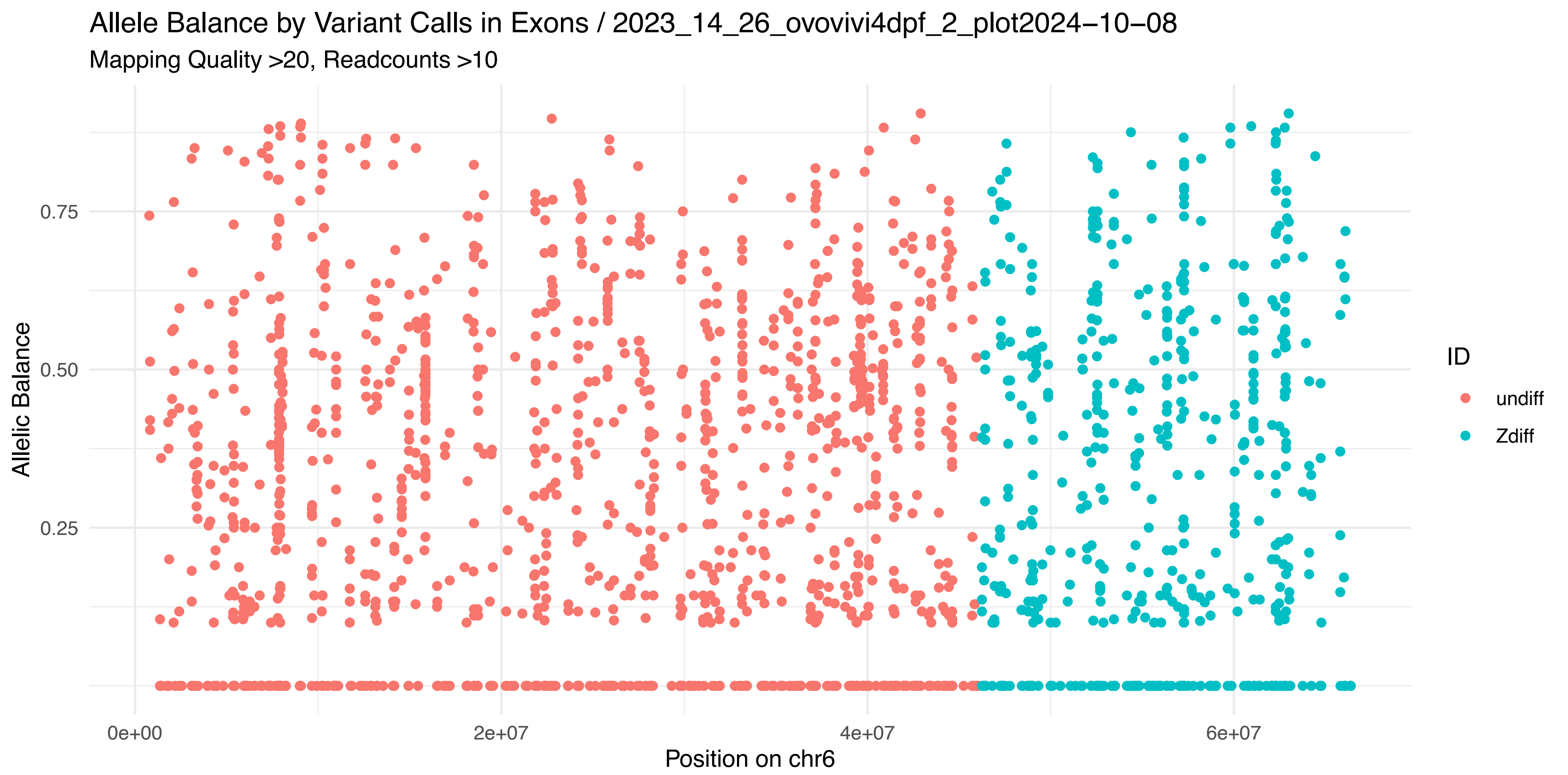
Allele Balance by Variant Calls in Exons / 2023_14_26_ovovivi4dpf_2_plot2024−10−08
Mapping Quality >20, Readcounts >10
Allelic Balance
0.75
0.50
0.25
0e+00
2e+07
4e+07
6e+07
Position on chr6
ID
undiff
Zdiff

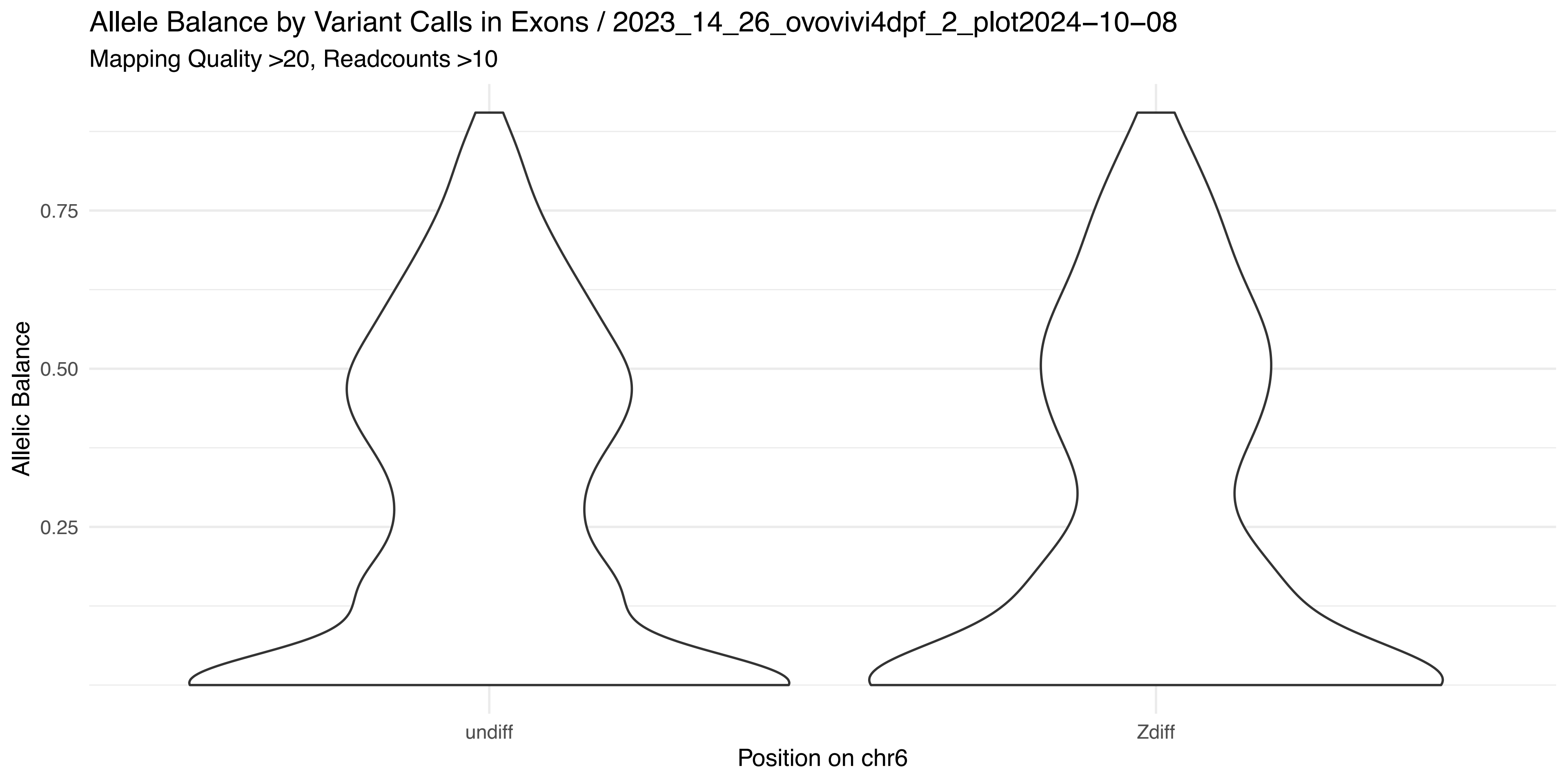
Allele Balance by Variant Calls in Exons / 2023_14_26_ovovivi4dpf_2_plot2024−10−08
Mapping Quality >20, Readcounts >10
Allelic Balance
0.75
0.50
0.25
undiff
Zdiff
Position on chr6

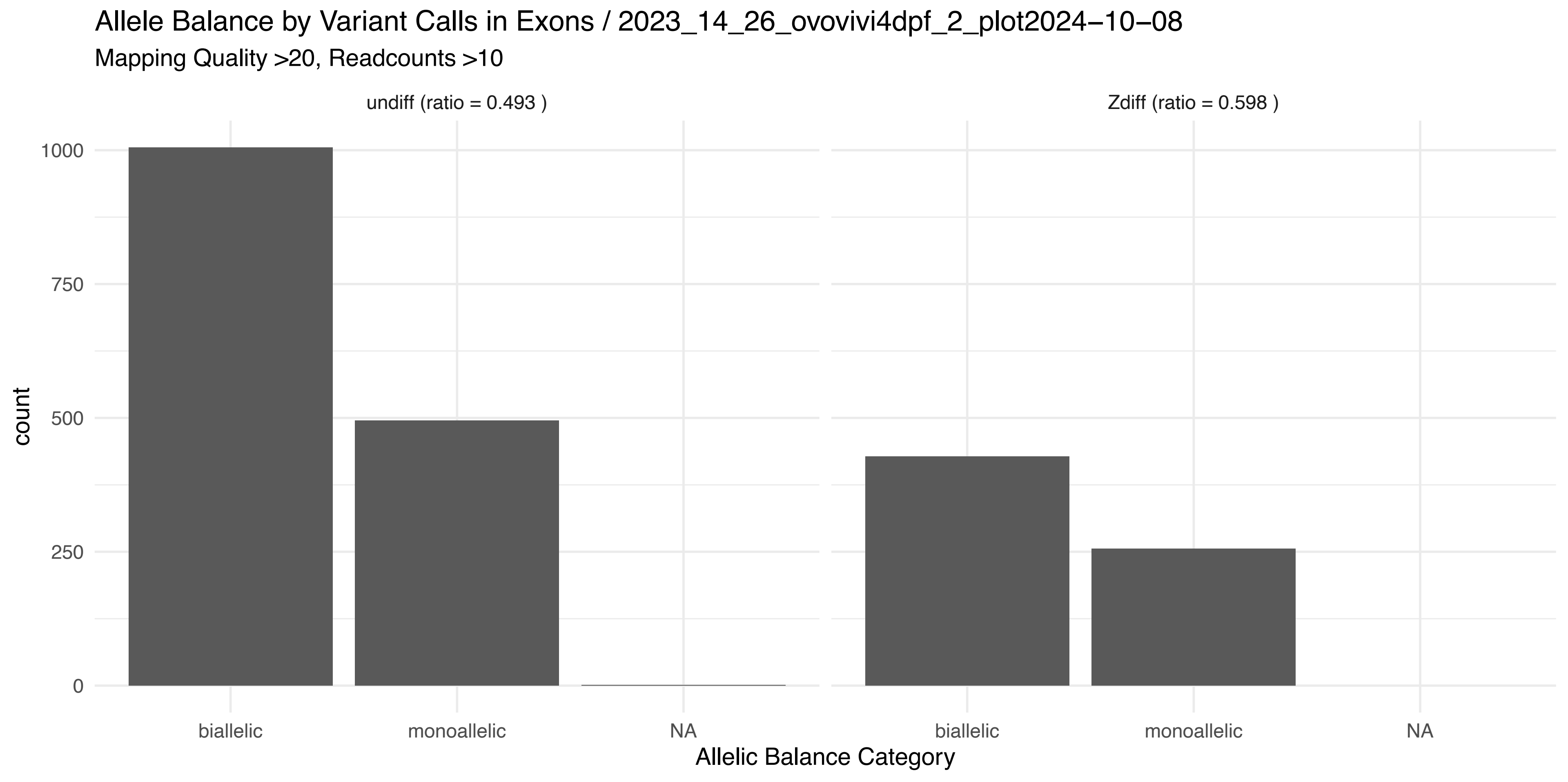
Allele Balance by Variant Calls in Exons / 2023_14_26_ovovivi4dpf_2_plot2024−10−08
Mapping Quality >20, Readcounts >10
undiff (ratio = 0.493 )
Zdiff (ratio = 0.598 )
count
1000
750
500
250
0
biallelic
monoallelic
NA
biallelic
monoallelic
NA
Allelic Balance Category

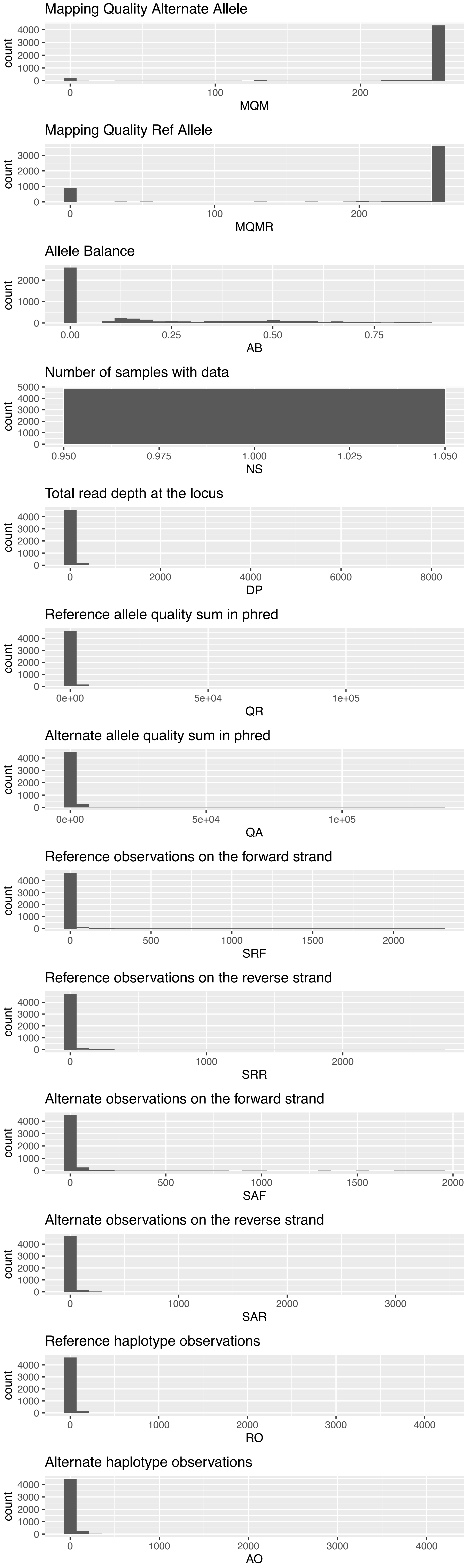

Mapping Quality Alternate Allele
count
0 100 200
MQM
Mapping Quality Ref Allele
count
0 100 200
MQMR
Allele Balance
count
0.00 0.25 0.50 0.75
AB
Number of samples with data
count
0.950 0.975 1.000 1.025 1.050
NS
Total read depth at the locus
count
0 2000 4000 6000 8000
DP
Reference allele quality sum in phred
count
0e+00 5e+04 1e+05
QR
Alternate allele quality sum in phred
count
0e+00 5e+04 1e+05
QA
Reference observations on the forward strand
count
0 500 1000 1500 2000
SRF
Reference observations on the reverse strand
count
0 1000 2000
SRR
Alternate observations on the forward strand
count
0 500 1000 1500 2000
SAF
Alternate observations on the reverse strand
count
0 1000 2000 3000
SAR
Reference haplotype observations
count
0 1000 2000 3000 4000
RO
Alternate haplotype observations
count
0 1000 2000 3000 4000
AO

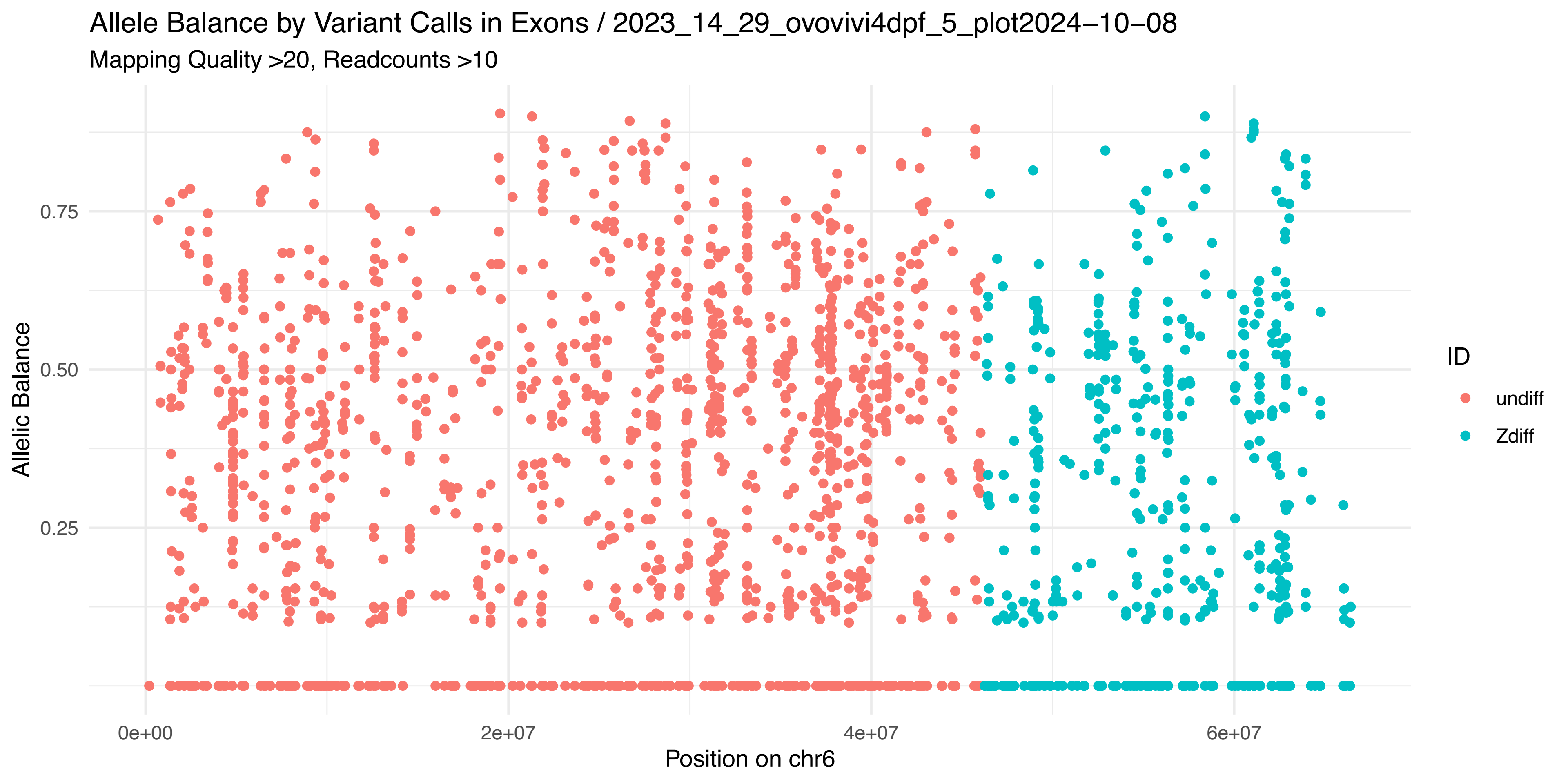
Allele Balance by Variant Calls in Exons / 2023_14_29_ovovivi4dpf_5_plot2024−10−08
Mapping Quality >20, Readcounts >10
Allelic Balance
0.75
0.50
0.25
0e+00
2e+07
4e+07
6e+07
Position on chr6
ID
undiff
Zdiff

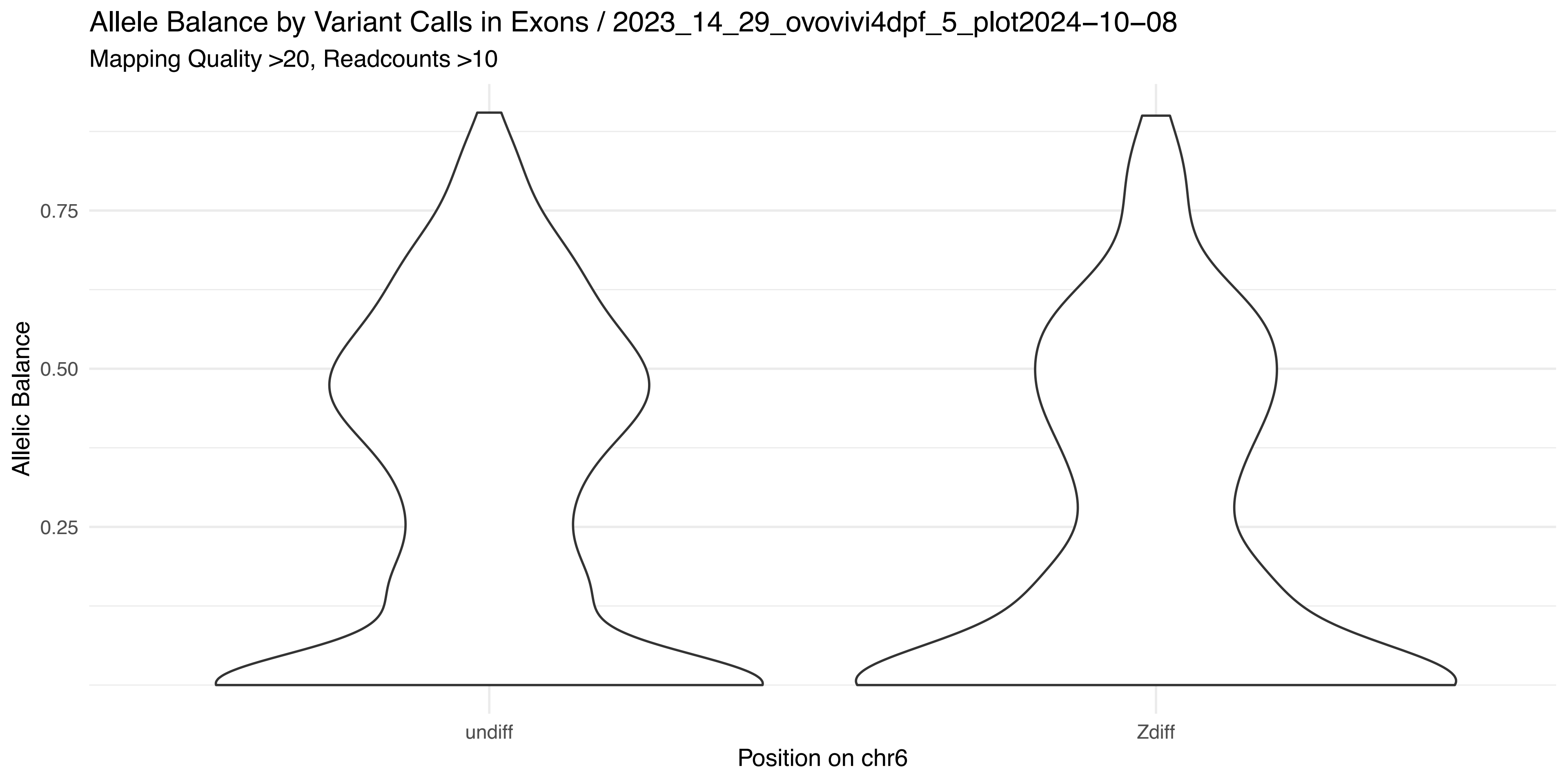
Allele Balance by Variant Calls in Exons / 2023_14_29_ovovivi4dpf_5_plot2024−10−08
Mapping Quality >20, Readcounts >10
Allelic Balance
0.75
0.50
0.25
undiff
Zdiff
Position on chr6

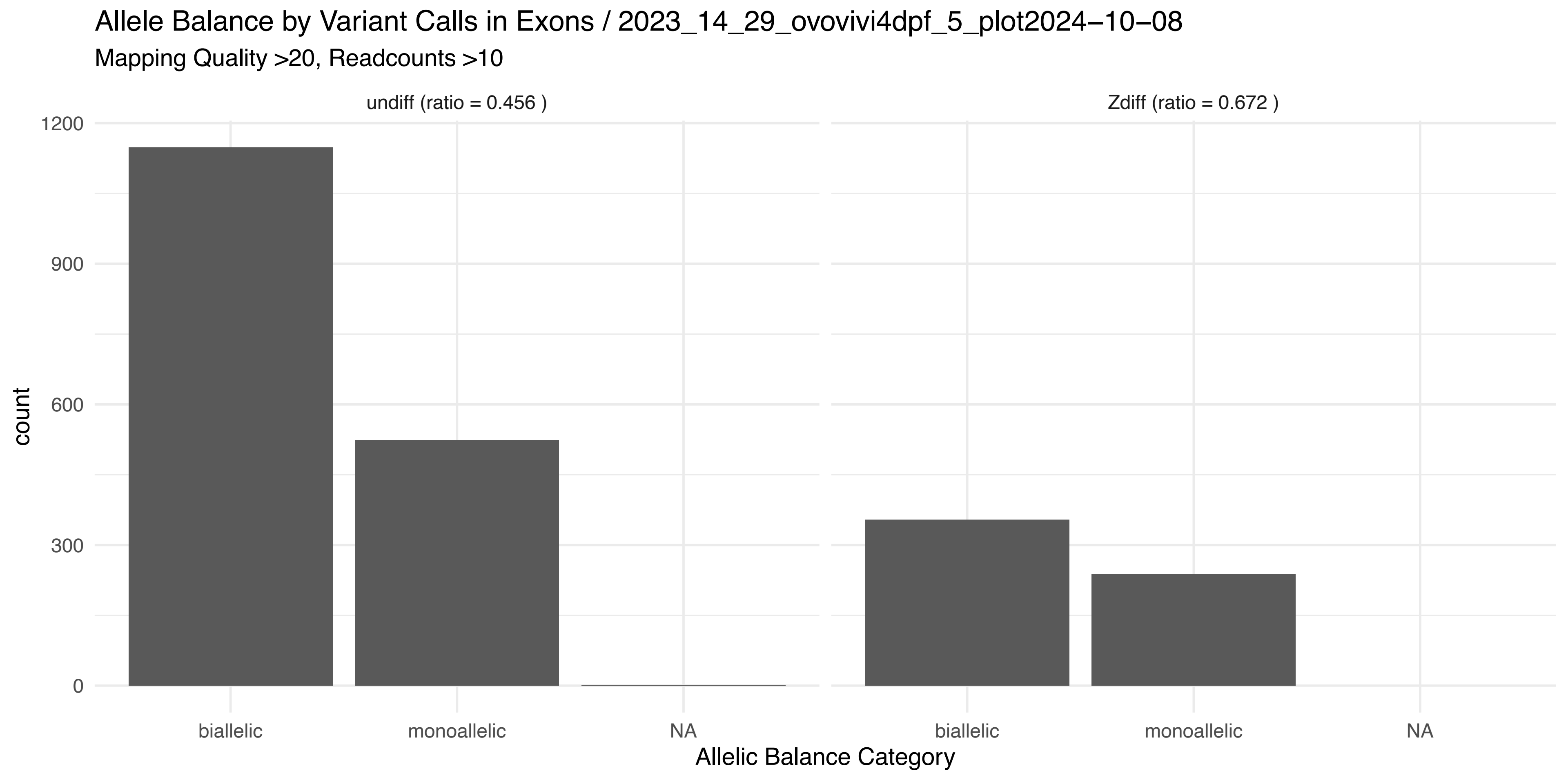
Allele Balance by Variant Calls in Exons / 2023_14_29_ovovivi4dpf_5_plot2024−10−08
Mapping Quality >20, Readcounts >10
undiff (ratio = 0.456 )
Zdiff (ratio = 0.672 )
count
1200
900
600
300
0
biallelic
monoallelic
NA
biallelic
monoallelic
NA
Allelic Balance Category

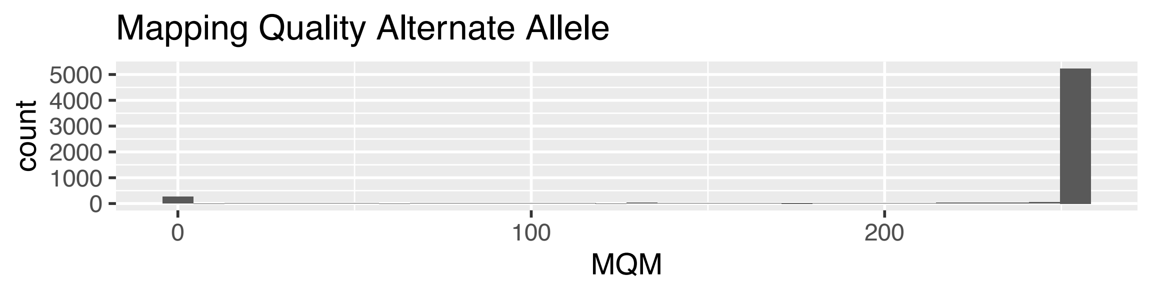
Mapping Quality Alternate Allele
count
0 1000 2000 3000 4000 5000
0 100 200
MQM

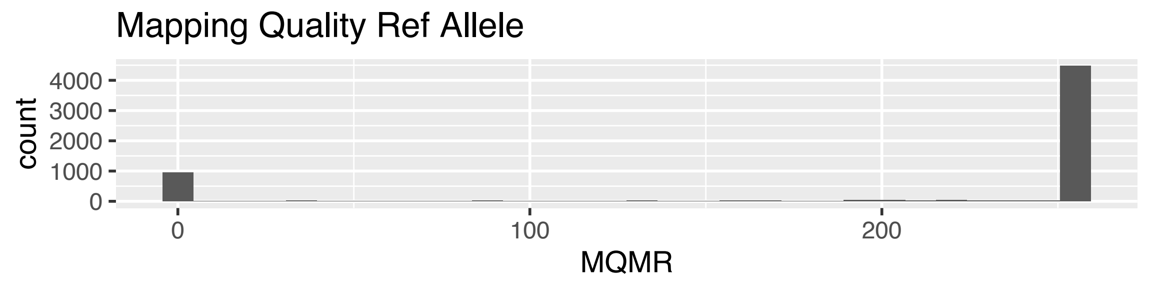
Mapping Quality Ref Allele
count
0 1000 2000 3000 4000
0 100 200
MQMR

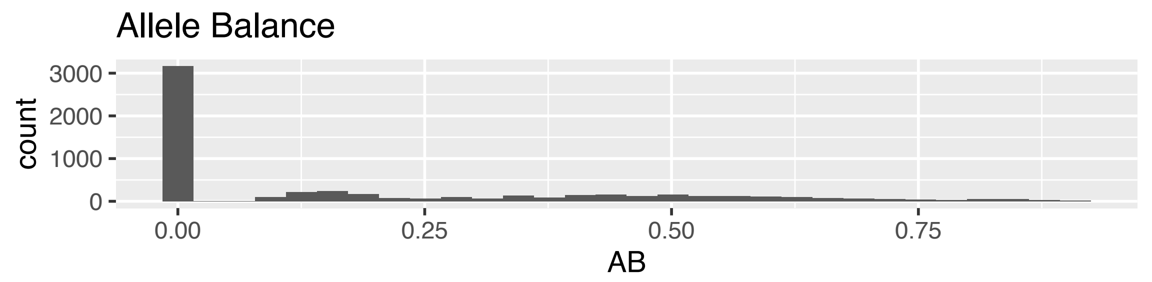
Allele Balance
count
0 1000 2000 3000
0.00 0.25 0.50 0.75
AB

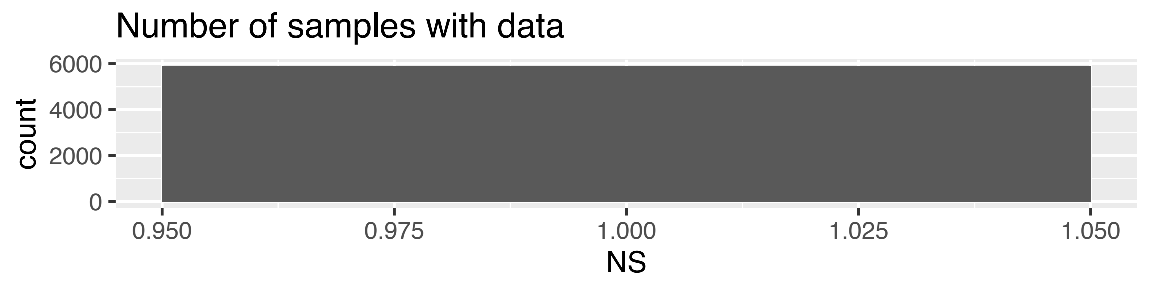
Number of samples with data
count
0 2000 4000 6000
0.950 0.975 1.000 1.025 1.050
NS

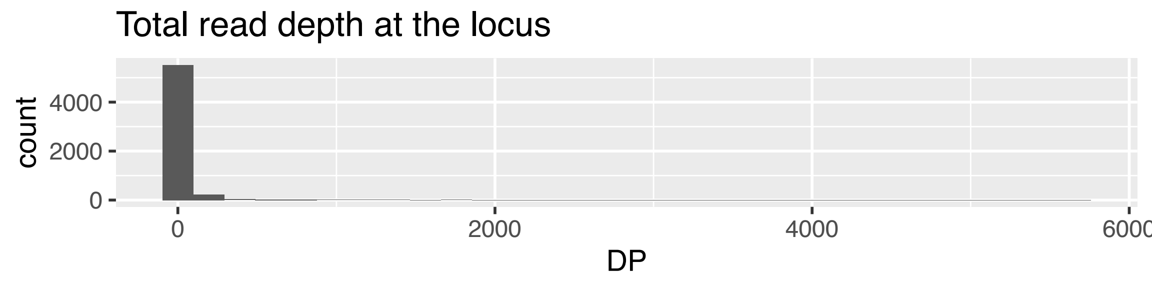
Total read depth at the locus
count
0 2000 4000
0 2000 4000 6000
DP

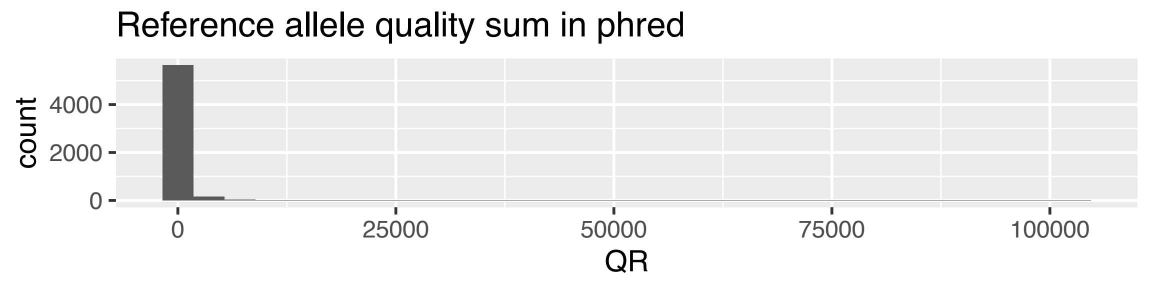
Reference allele quality sum in phred
count
0 2000 4000
0 25000 50000 75000 100000
QR

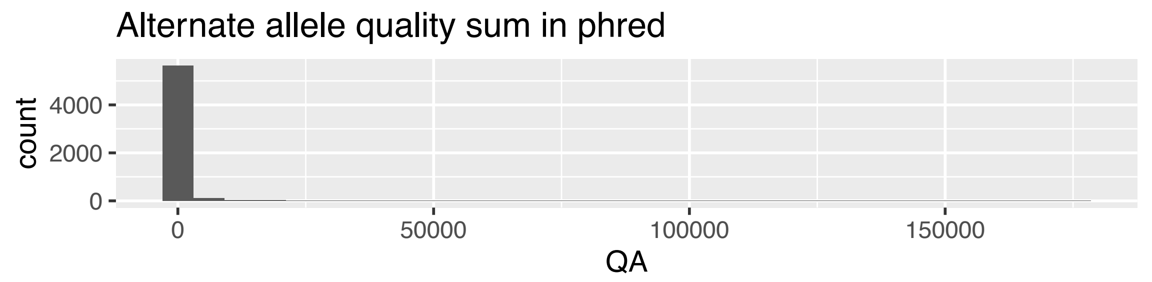
Alternate allele quality sum in phred
count
0 2000 4000
0 50000 100000 150000
QA

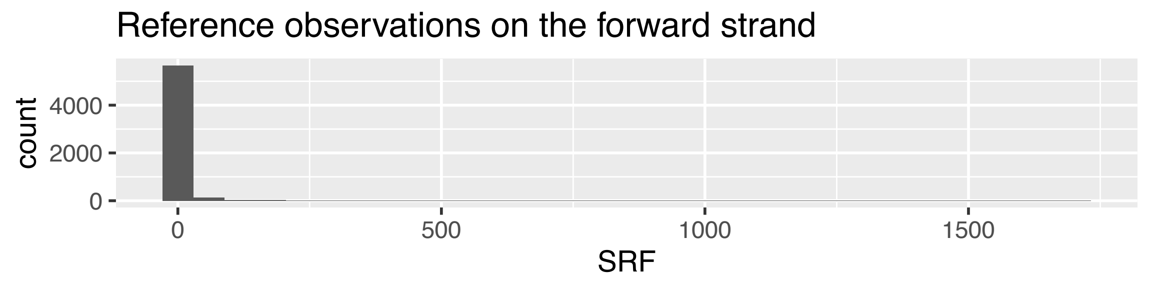
Reference observations on the forward strand
count
0 2000 4000
0 500 1000 1500
SRF

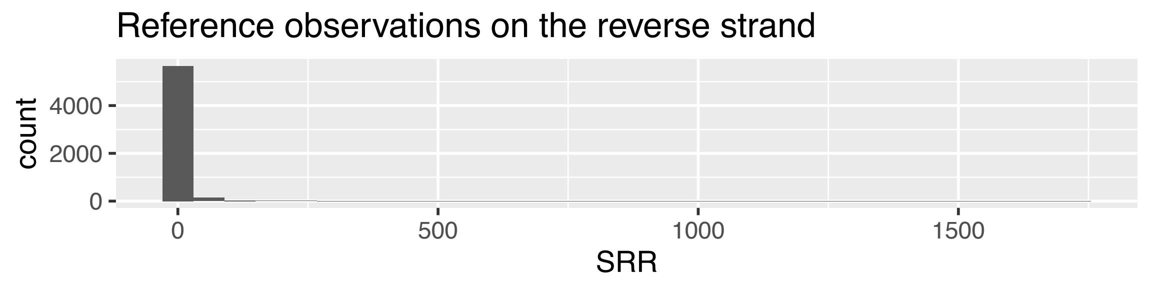
Reference observations on the reverse strand
count
0 2000 4000
0 500 1000 1500
SRR

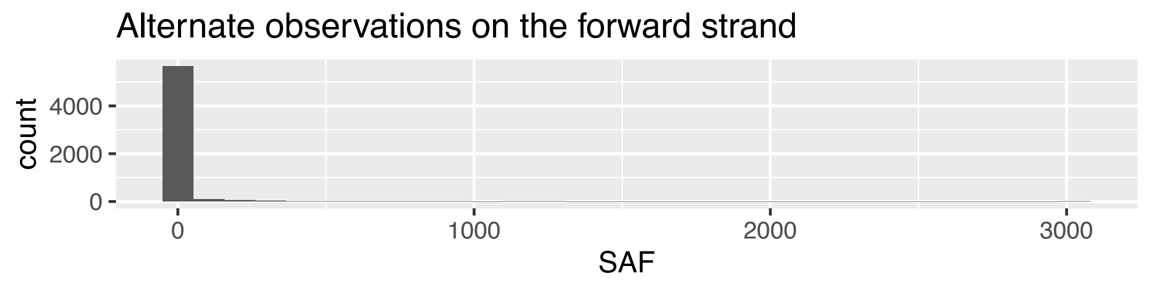
Alternate observations on the forward strand
count
0 2000 4000
0 1000 2000 3000
SAF

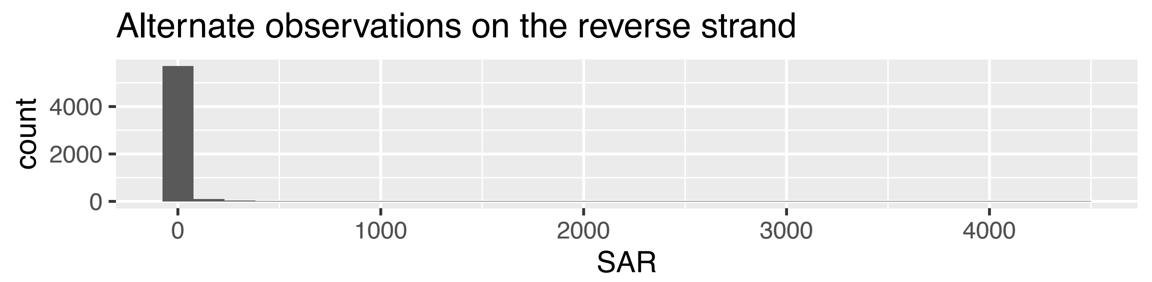
Alternate observations on the reverse strand
count
0 2000 4000
0 1000 2000 3000 4000
SAR

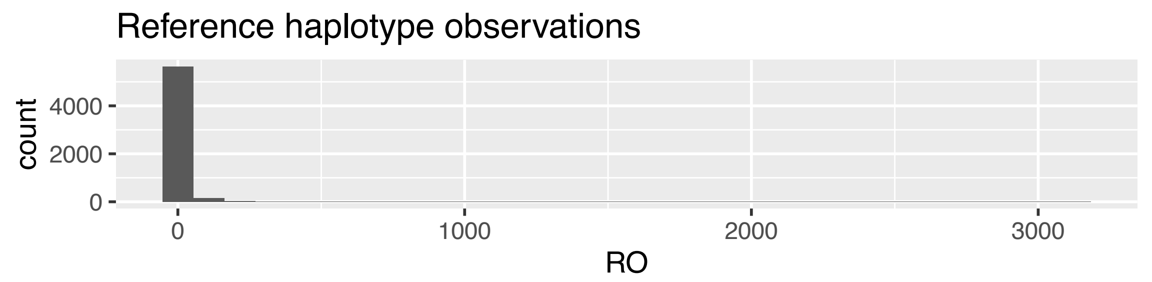
Reference haplotype observations
count
0 2000 4000
0 1000 2000 3000
RO

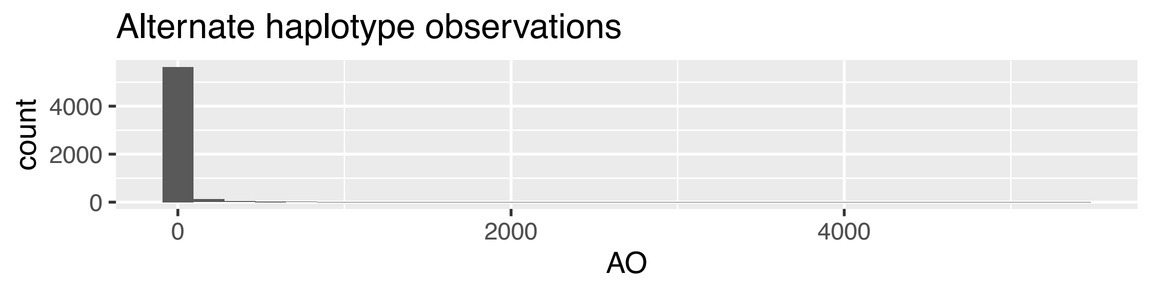
Alternate haplotype observations
count
0 2000 4000
0 2000 4000
AO

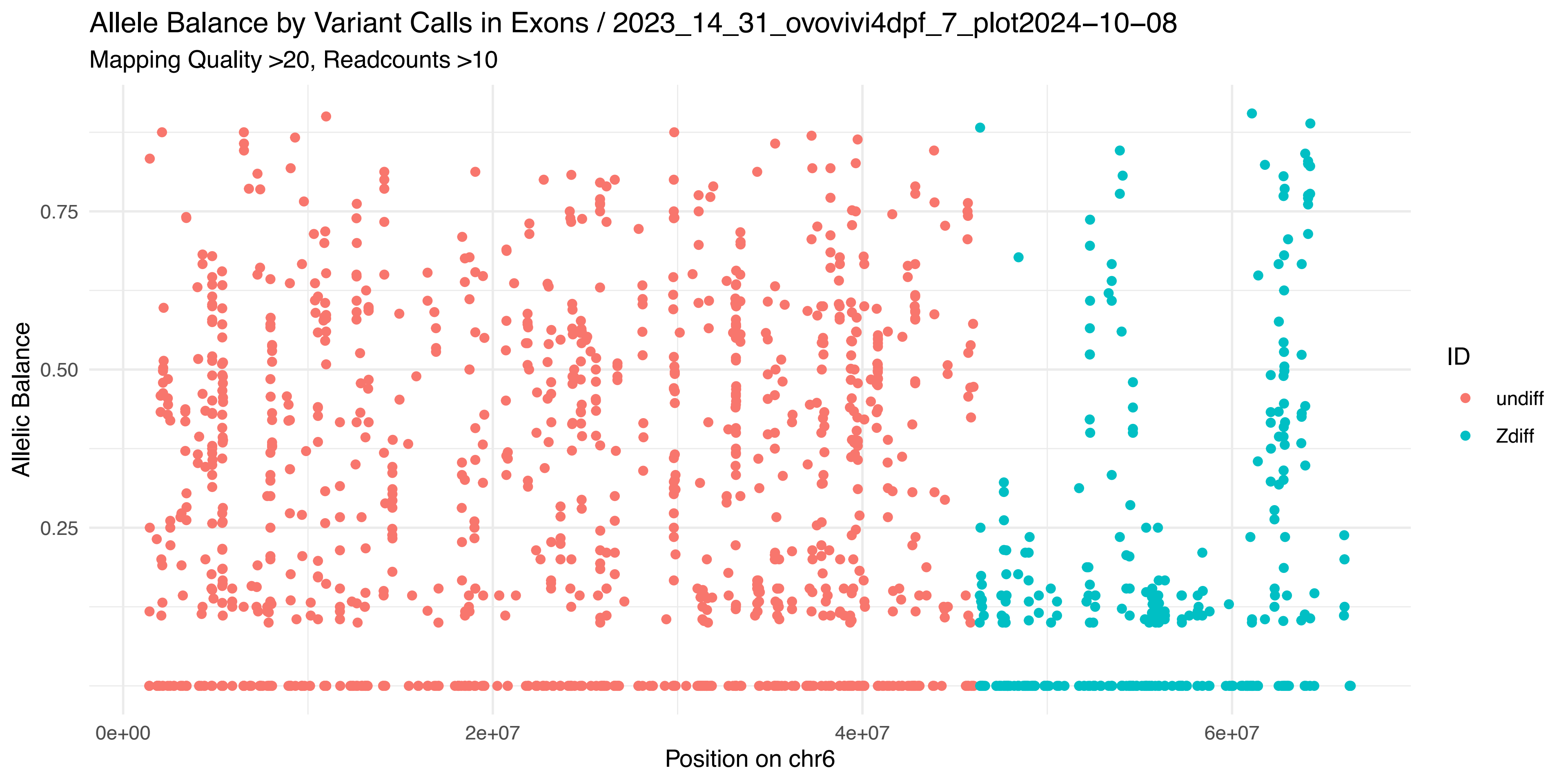

Allele Balance by Variant Calls in Exons / 2023_14_31_ovovivi4dpf_7_plot2024−10−08
Mapping Quality >20, Readcounts >10
Allelic Balance
0.25
0.50
0.75
0e+00
2e+07
4e+07
6e+07
Position on chr6
ID
undiff
Zdiff


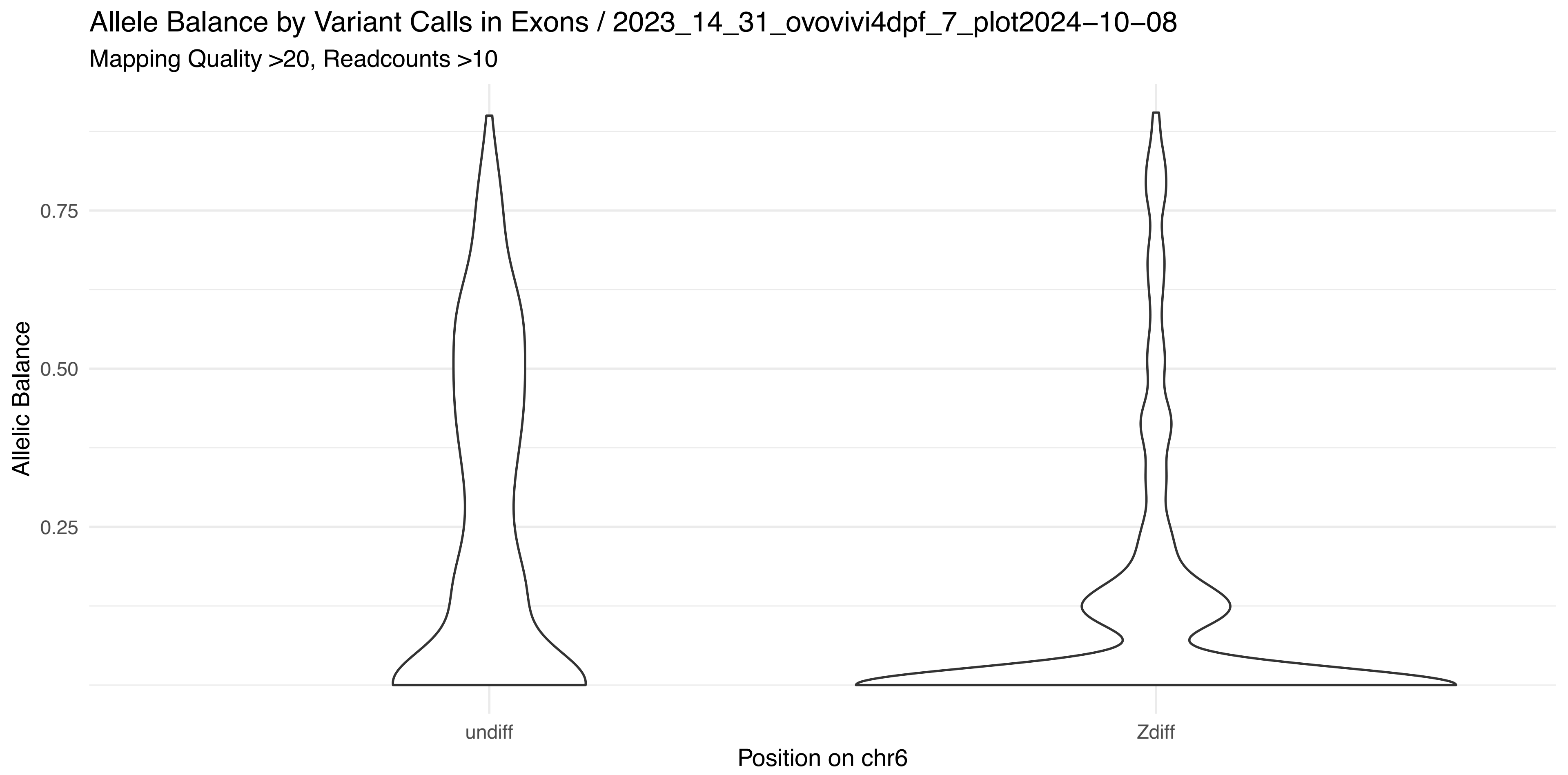

Allele Balance by Variant Calls in Exons / 2023_14_31_ovovivi4dpf_7_plot2024−10−08
Mapping Quality >20, Readcounts >10
Allelic Balance
0.25
0.50
0.75
undiff
Zdiff
Position on chr6


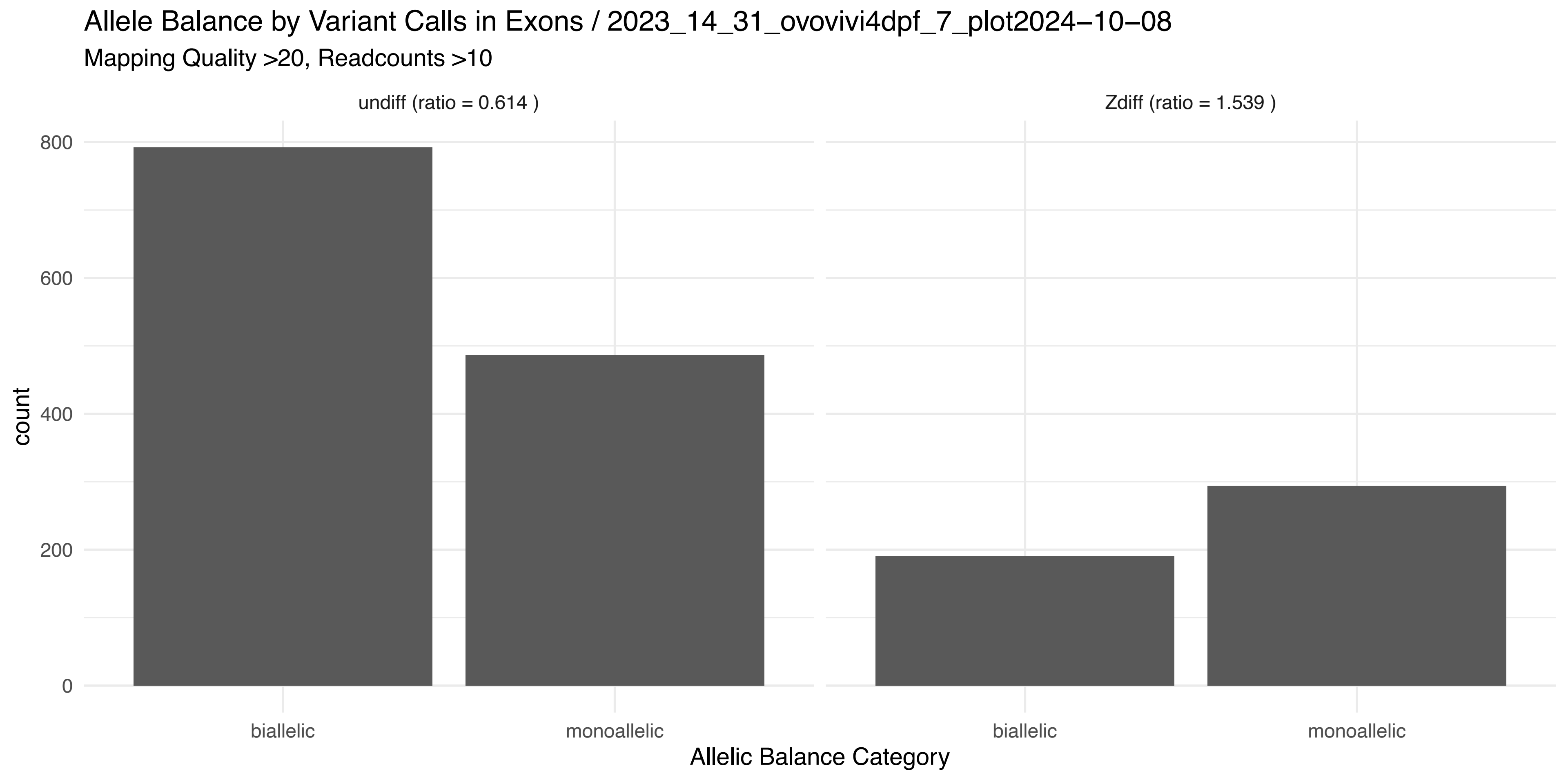

Allele Balance by Variant Calls in Exons / 2023_14_31_ovovivi4dpf_7_plot2024−10−08
Mapping Quality >20, Readcounts >10
undiff (ratio = 0.614 )
Zdiff (ratio = 1.539 )
count
0
200
400
600
800
biallelic
monoallelic
biallelic
monoallelic
Allelic Balance Category

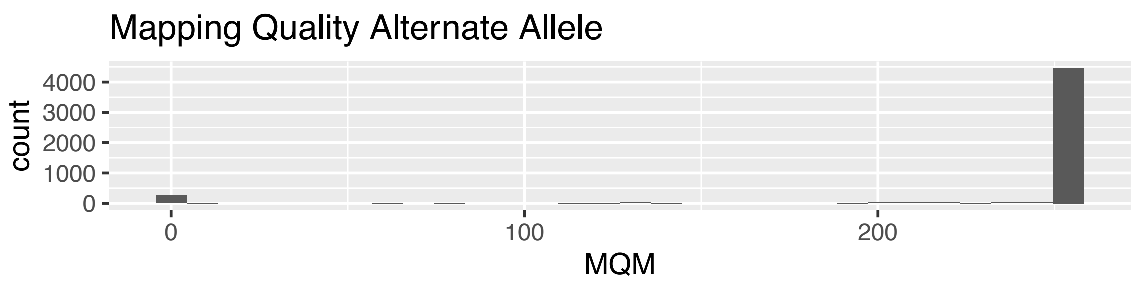
Mapping Quality Alternate Allele
count
0
1000
2000
3000
4000
0
100
200
MQM

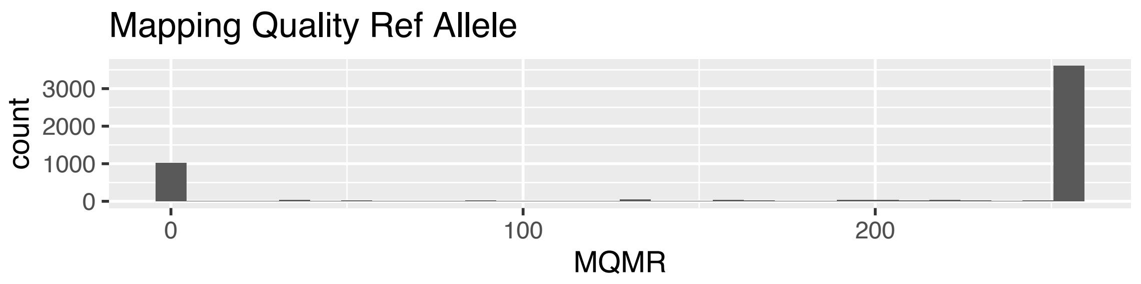
Mapping Quality Ref Allele
count
0
1000
2000
3000
0
100
200
MQMR

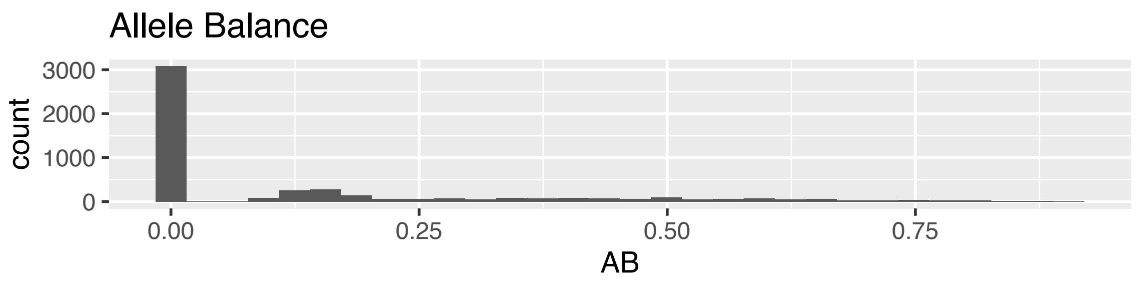
Allele Balance
count
0
1000
2000
3000
0.00
0.25
0.50
0.75
AB

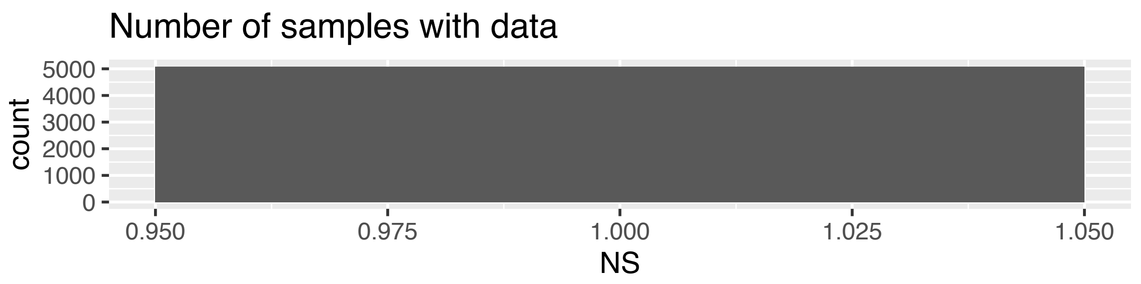
Number of samples with data
count
0
1000
2000
3000
4000
5000
0.950
0.975
1.000
1.025
1.050
NS

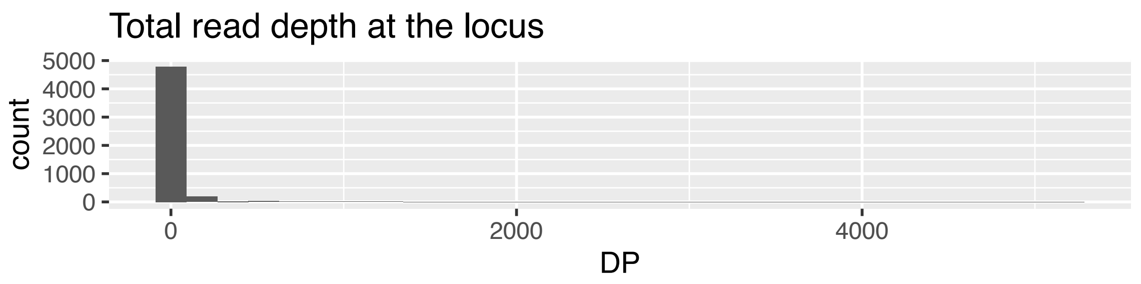
Total read depth at the locus
count
0
1000
2000
3000
4000
5000
0
2000
4000
DP

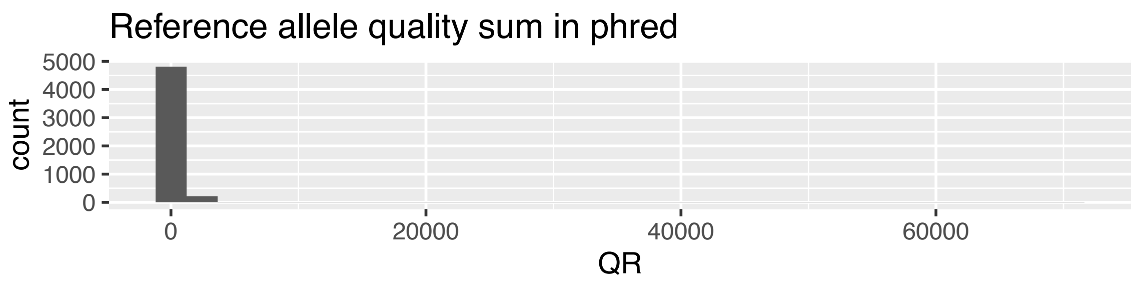
Reference allele quality sum in phred
count
0
1000
2000
3000
4000
5000
0
20000
40000
60000
QR

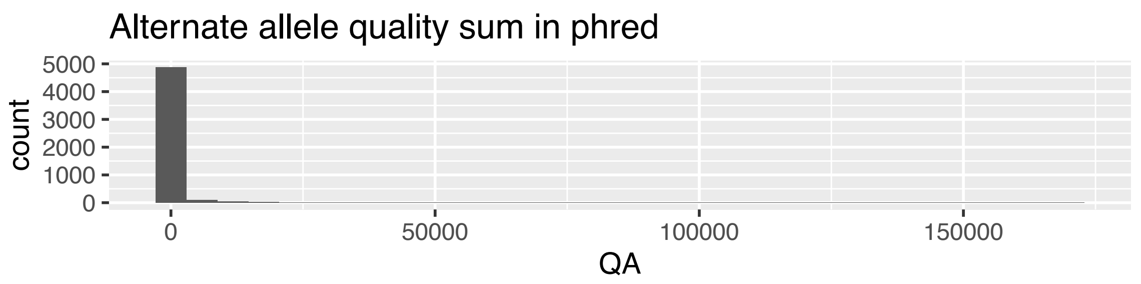
Alternate allele quality sum in phred
count
0
1000
2000
3000
4000
5000
0
50000
100000
150000
QA

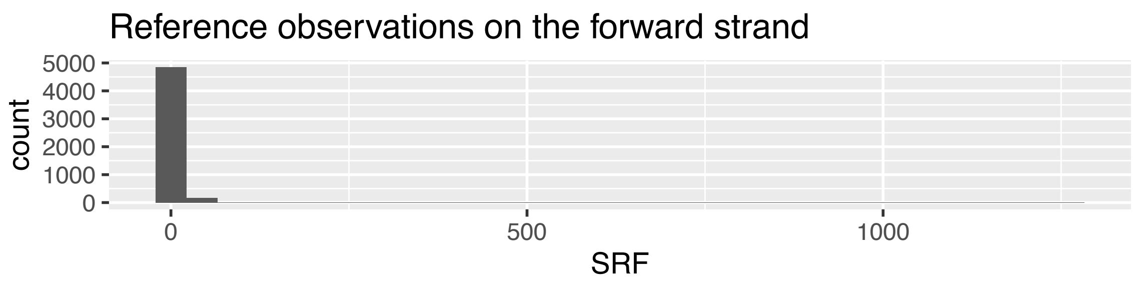
Reference observations on the forward strand
count
0
1000
2000
3000
4000
5000
0
500
1000
SRF

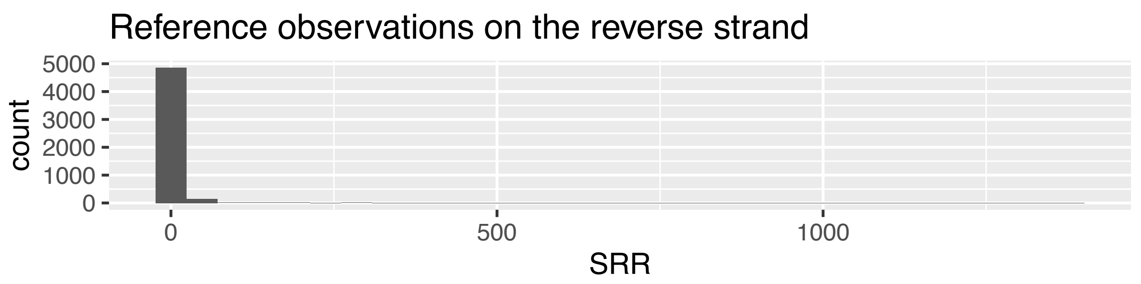
Reference observations on the reverse strand
count
0
1000
2000
3000
4000
5000
0
500
1000
SRR

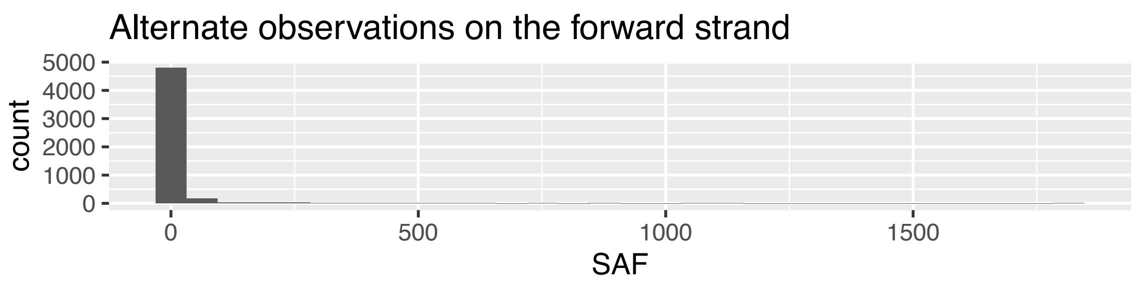
Alternate observations on the forward strand
count
0
1000
2000
3000
4000
5000
0
500
1000
1500
SAF

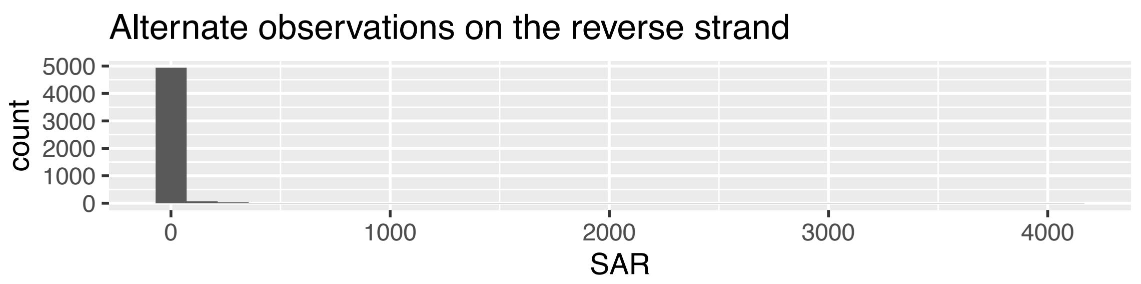
Alternate observations on the reverse strand
count
0
1000
2000
3000
4000
5000
0
1000
2000
3000
4000
SAR

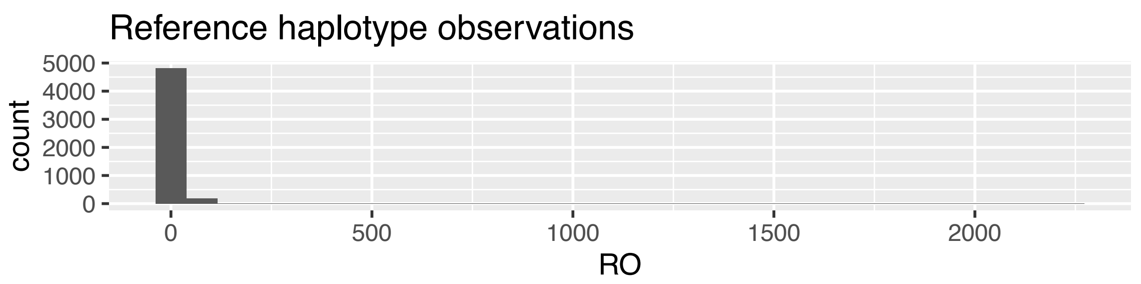
Reference haplotype observations
count
0
1000
2000
3000
4000
5000
0
500
1000
1500
2000
RO

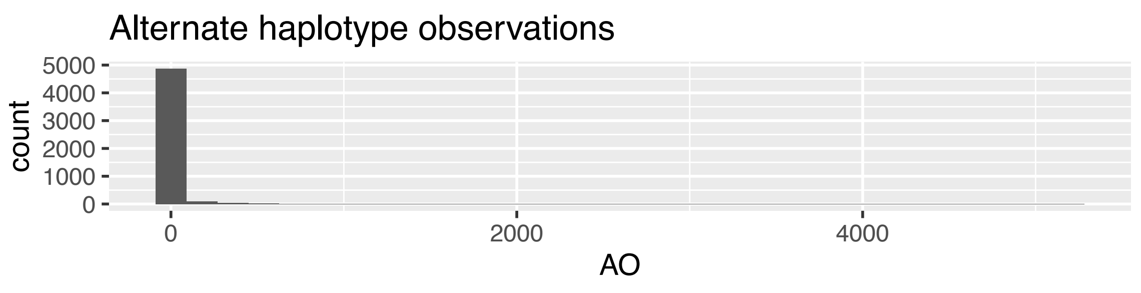
Alternate haplotype observations
count
0
1000
2000
3000
4000
5000
0
2000
4000
AO

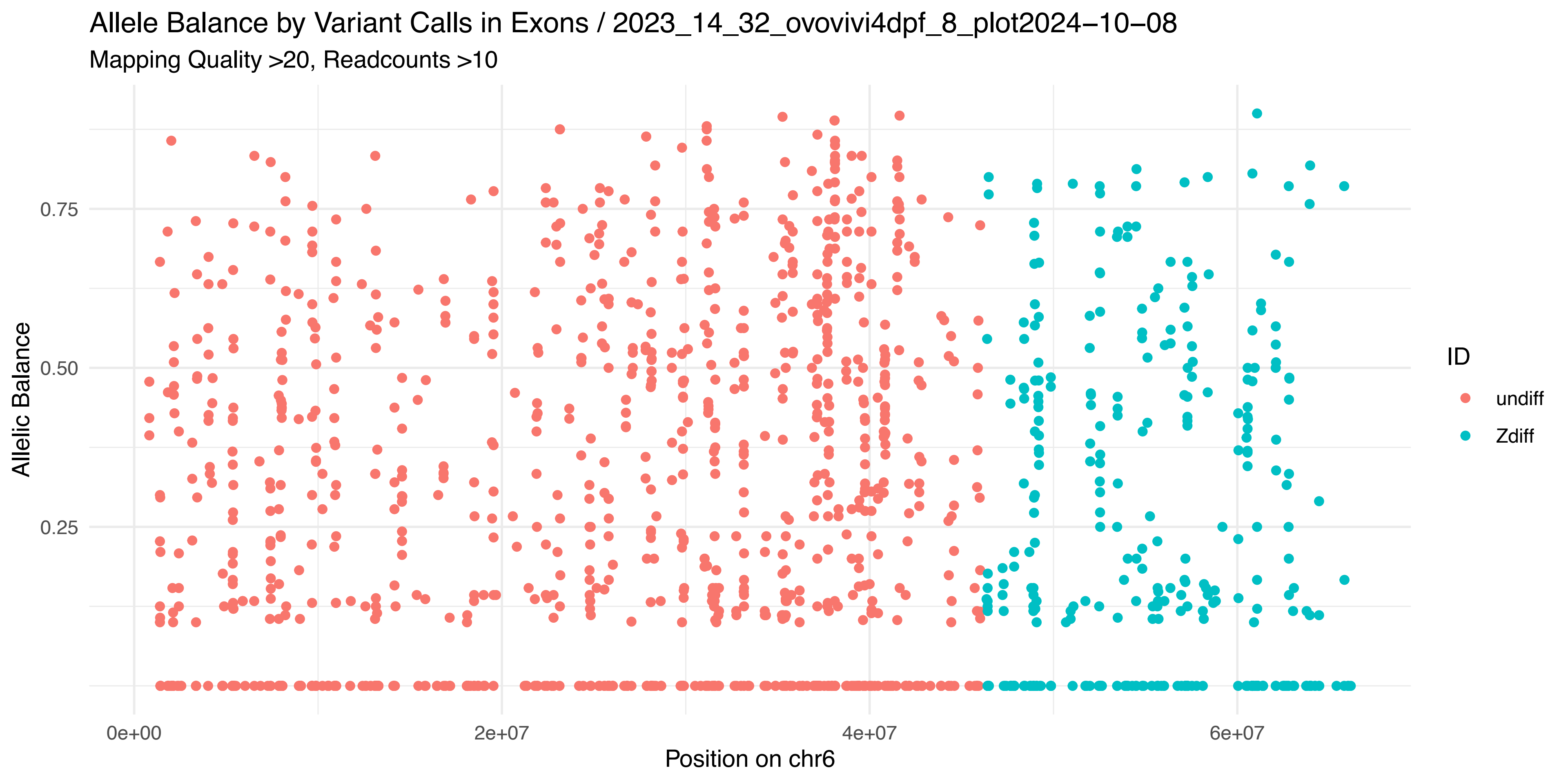
Allele Balance by Variant Calls in Exons / 2023_14_32_ovovivi4dpf_8_plot2024−10−08
Mapping Quality >20, Readcounts >10
Allelic Balance
0.25
0.50
0.75
0e+00
2e+07
4e+07
6e+07
Position on chr6
ID
undiff
Zdiff

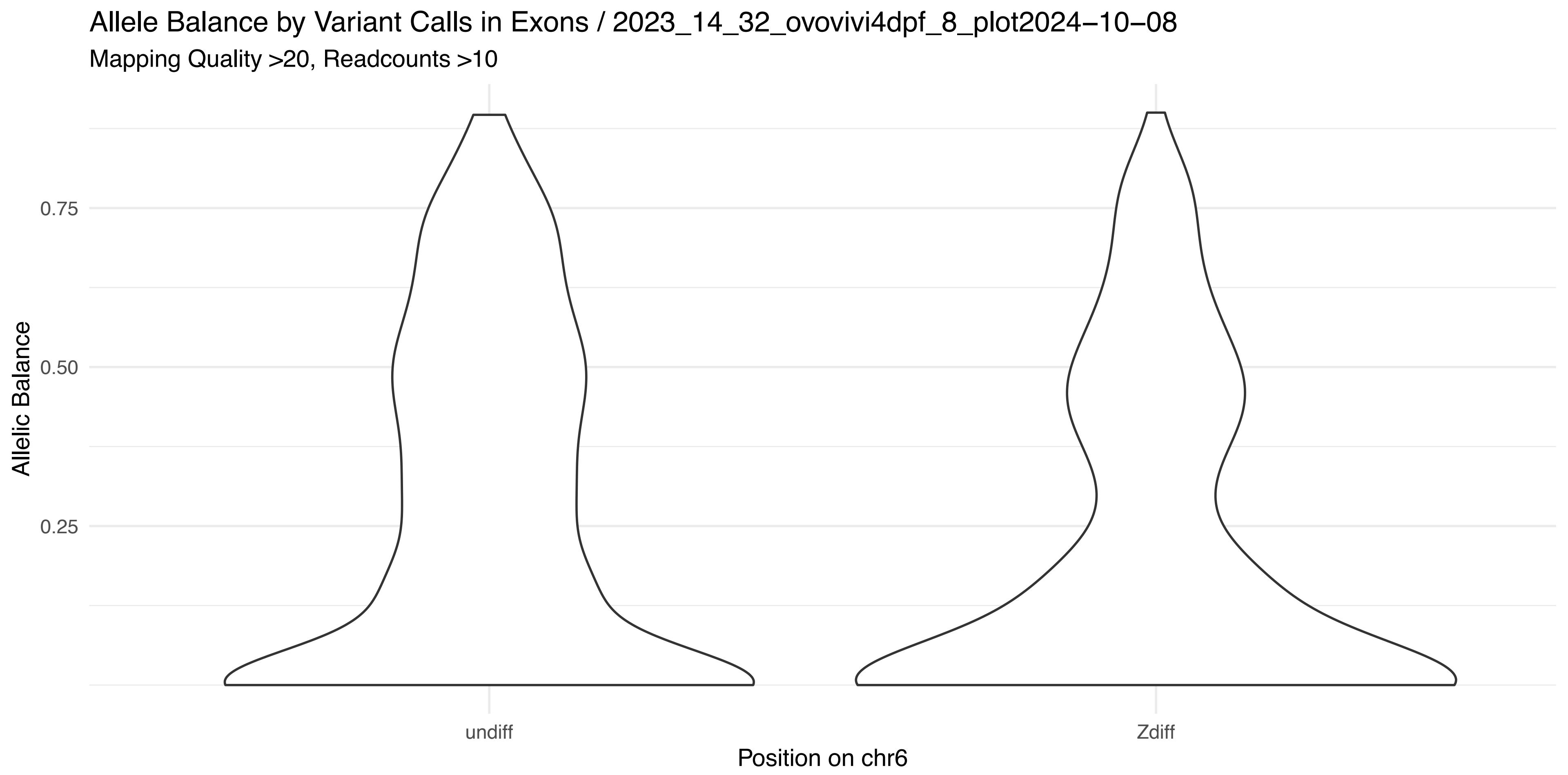
Allele Balance by Variant Calls in Exons / 2023_14_32_ovovivi4dpf_8_plot2024−10−08
Mapping Quality >20, Readcounts >10
Allelic Balance
0.25
0.50
0.75
undiff
Zdiff
Position on chr6

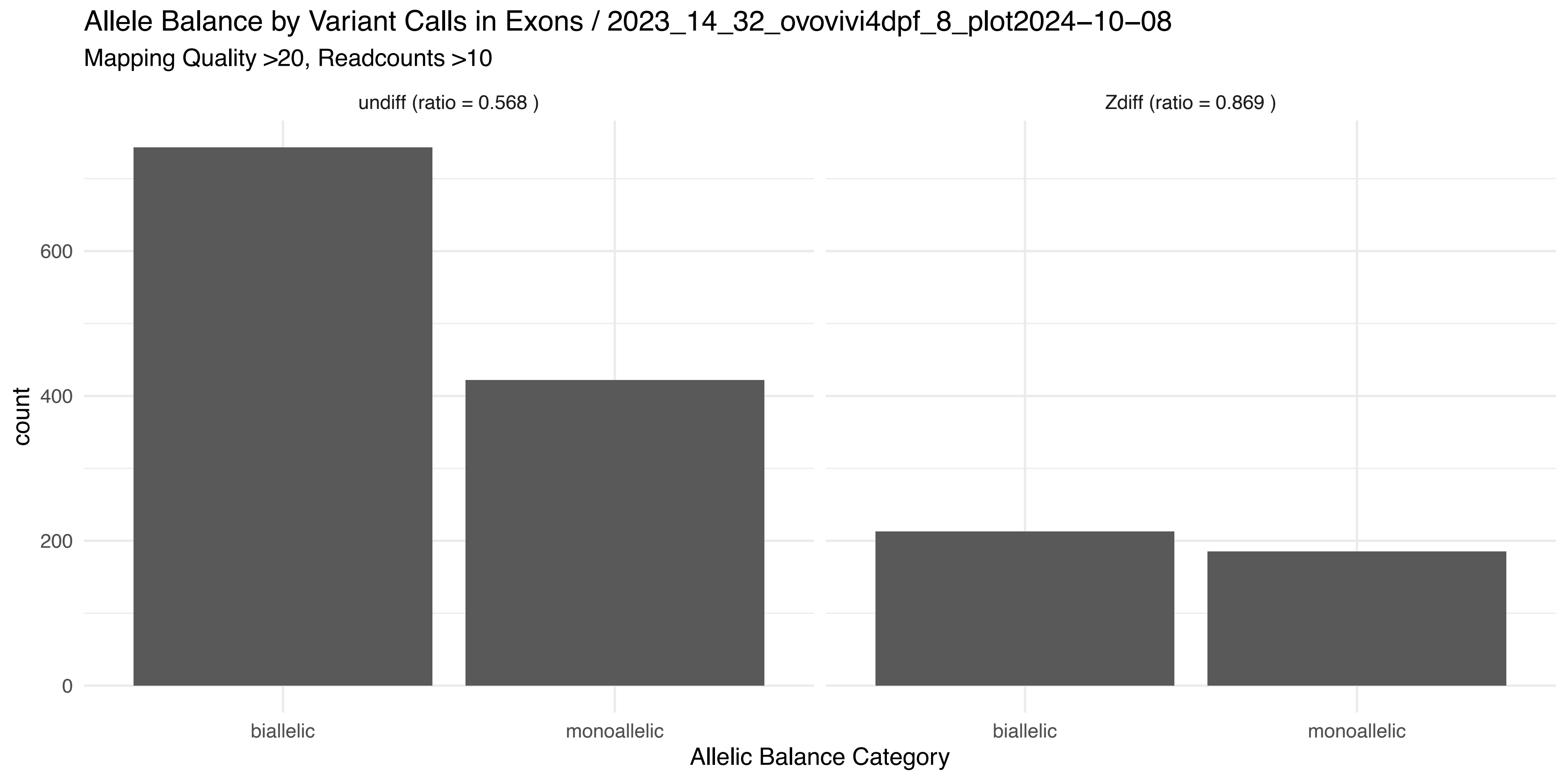
Allele Balance by Variant Calls in Exons / 2023_14_32_ovovivi4dpf_8_plot2024−10−08
Mapping Quality >20, Readcounts >10
undiff (ratio = 0.568 )
Zdiff (ratio = 0.869 )
count
0
200
400
600
biallelic
monoallelic
biallelic
monoallelic
Allelic Balance Category

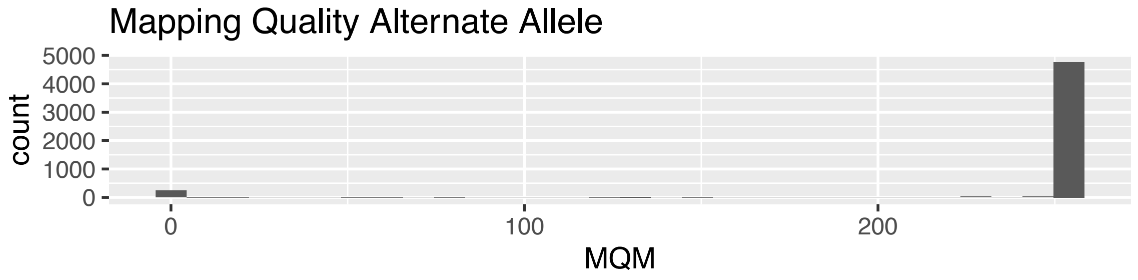
Mapping Quality Alternate Allele
count
5000
4000
3000
2000
1000
0
0
100
200
MQM

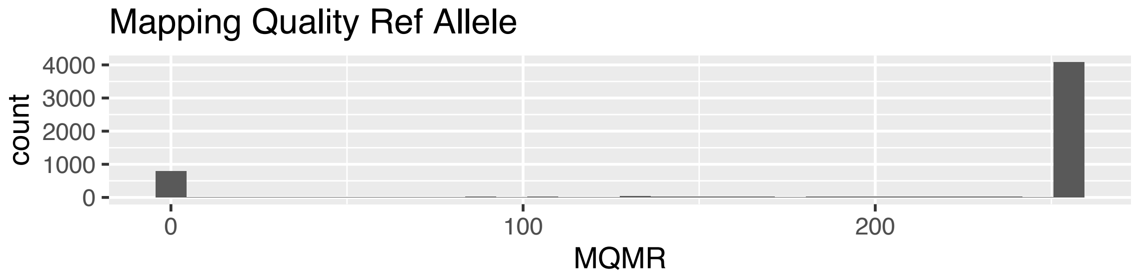
Mapping Quality Ref Allele
count
4000
3000
2000
1000
0
0
100
200
MQMR

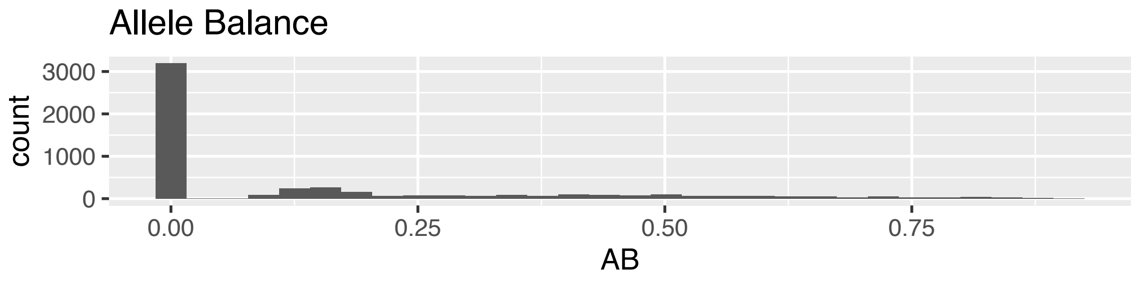
Allele Balance
count
3000
2000
1000
0
0.00
0.25
0.50
0.75
AB

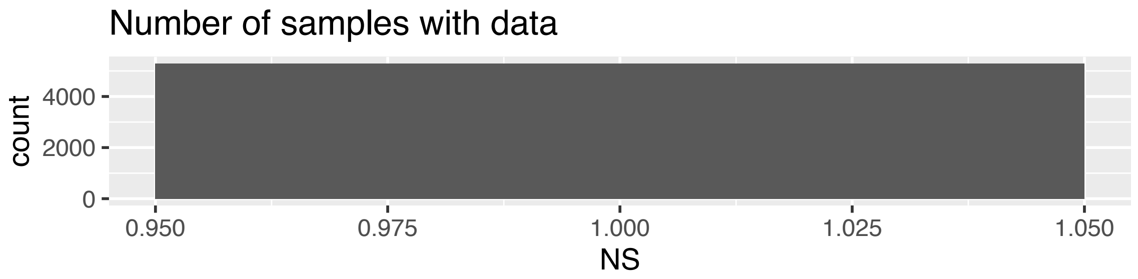
Number of samples with data
count
4000
2000
0
0.950
0.975
1.000
1.025
1.050
NS

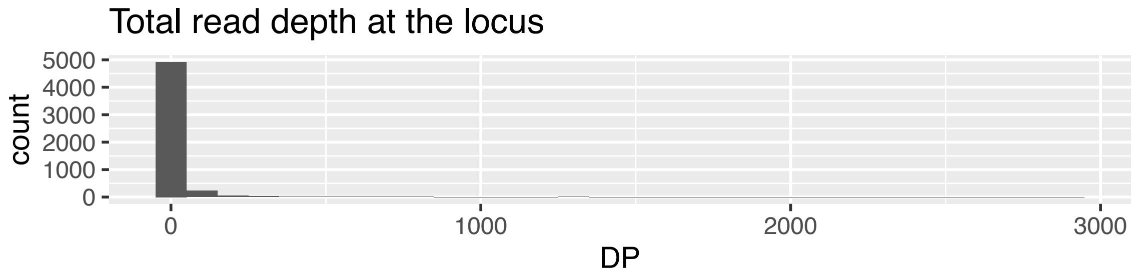
Total read depth at the locus
count
5000
4000
3000
2000
1000
0
0
1000
2000
3000
DP

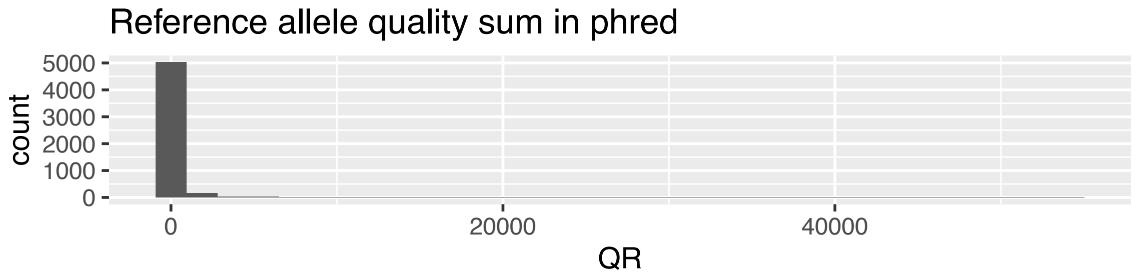
Reference allele quality sum in phred
count
5000
4000
3000
2000
1000
0
0
20000
40000
QR

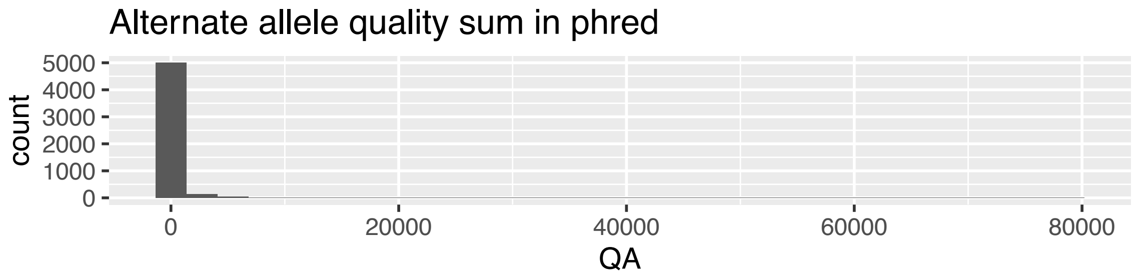
Alternate allele quality sum in phred
count
5000
4000
3000
2000
1000
0
0
20000
40000
60000
80000
QA

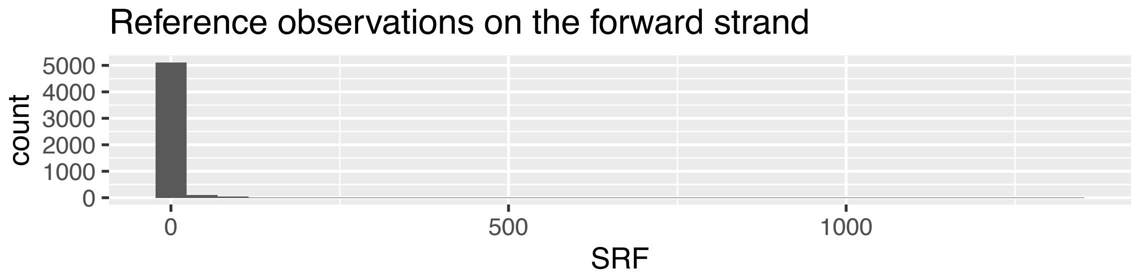
Reference observations on the forward strand
count
5000
4000
3000
2000
1000
0
0
500
1000
SRF

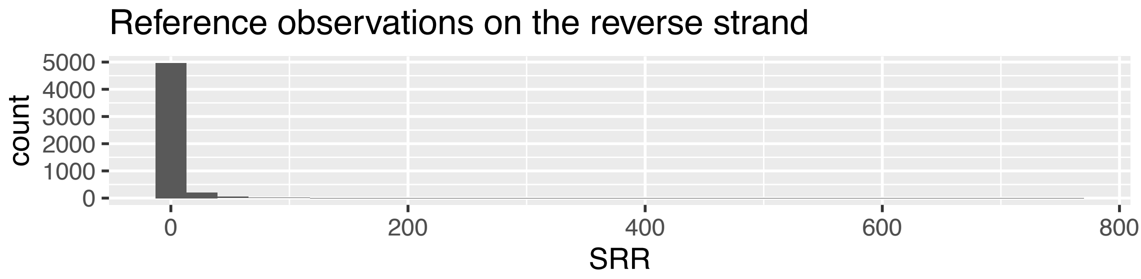
Reference observations on the reverse strand
count
5000
4000
3000
2000
1000
0
0
200
400
600
800
SRR

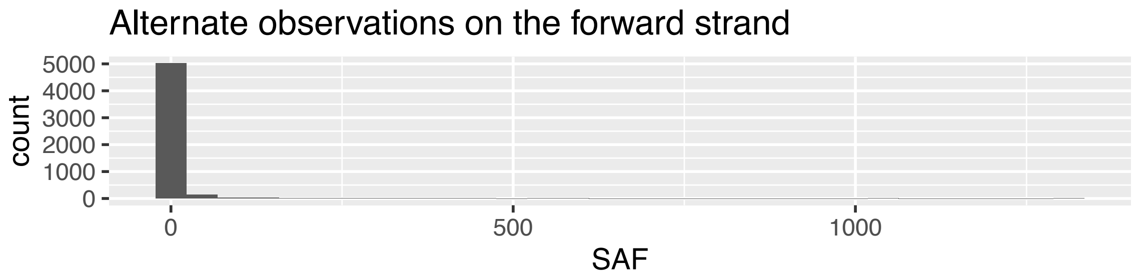
Alternate observations on the forward strand
count
5000
4000
3000
2000
1000
0
0
500
1000
SAF

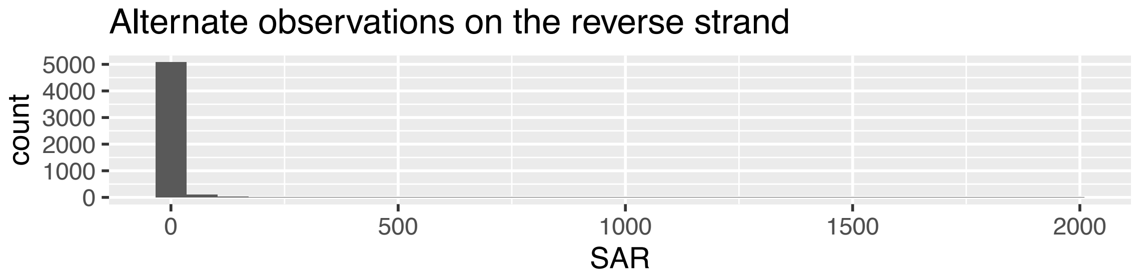
Alternate observations on the reverse strand
count
5000
4000
3000
2000
1000
0
0
500
1000
1500
2000
SAR

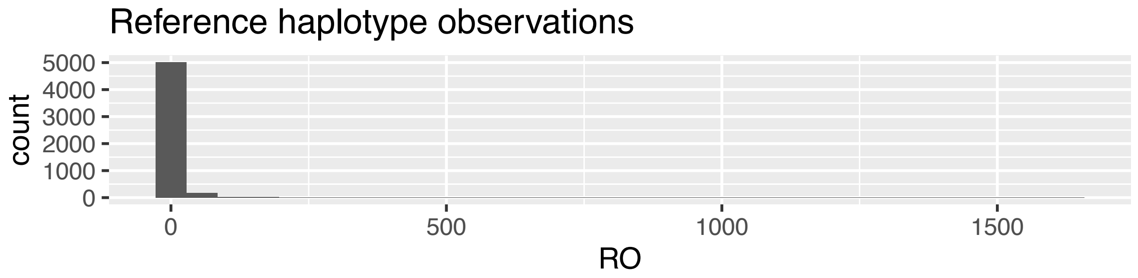
Reference haplotype observations
count
5000
4000
3000
2000
1000
0
0
500
1000
1500
RO

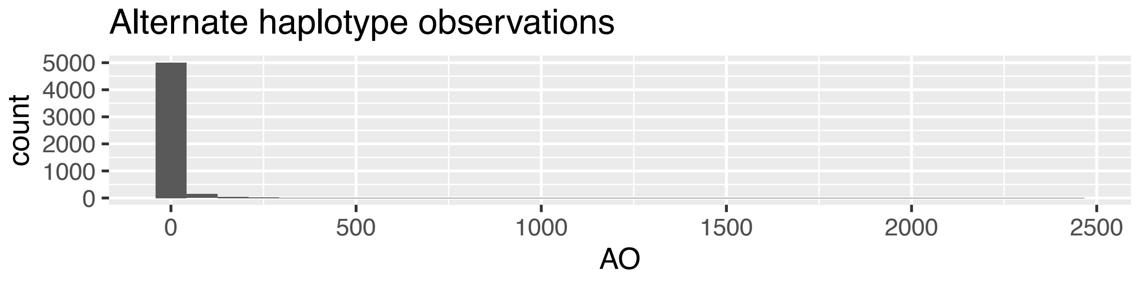
Alternate haplotype observations
count
5000
4000
3000
2000
1000
0
0
500
1000
1500
2000
2500
AO

Allele Balance by Variant Calls in Exons / 2023_14_34_ovovivi4dpf_10_plot2024−10−08

Mapping Quality >20, Readcounts >10

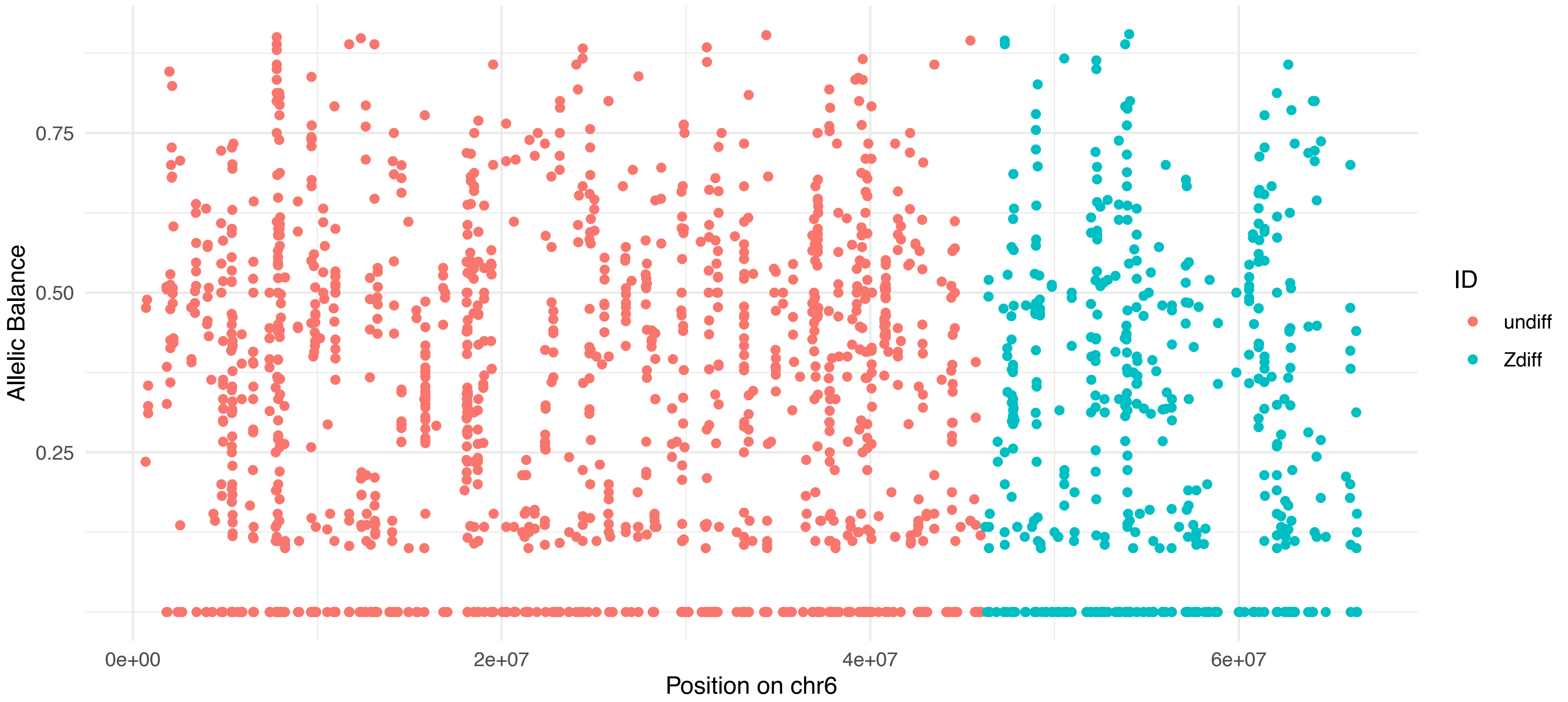


Allele Balance by Variant Calls in Exons / 2023_14_34_ovovivi4dpf_10_plot2024−10−08

Mapping Quality >20, Readcounts >10

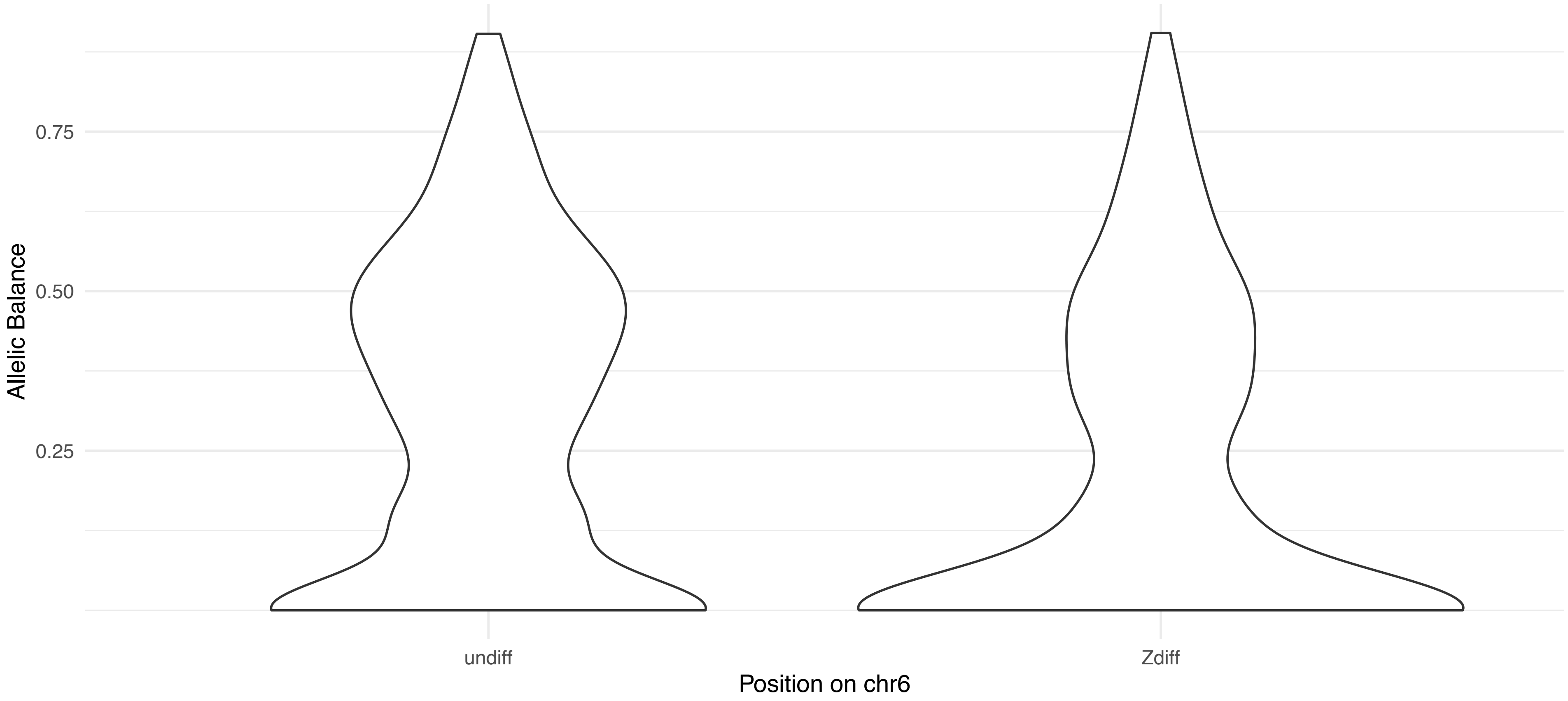


Allele Balance by Variant Calls in Exons / 2023_14_34_ovovivi4dpf_10_plot2024−10−08

Mapping Quality >20, Readcounts >10

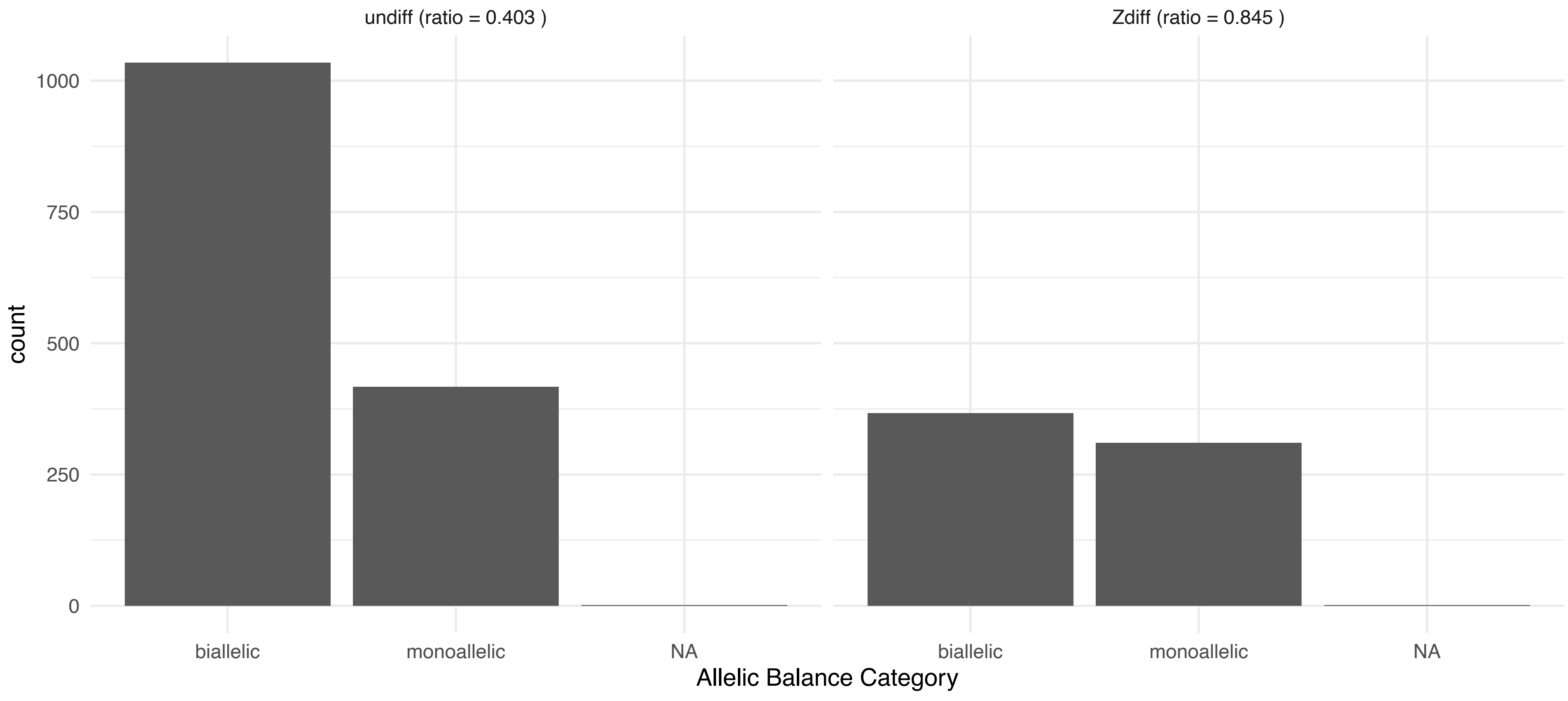

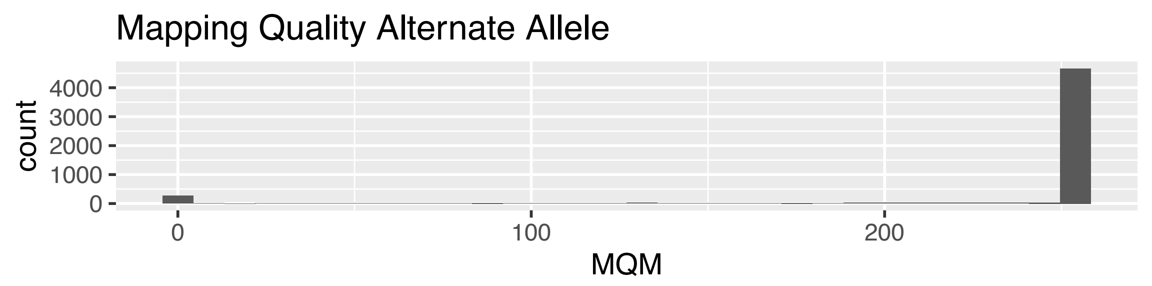
Mapping Quality Alternate Allele
count
0
1000
2000
3000
4000
0
100
200
MQM

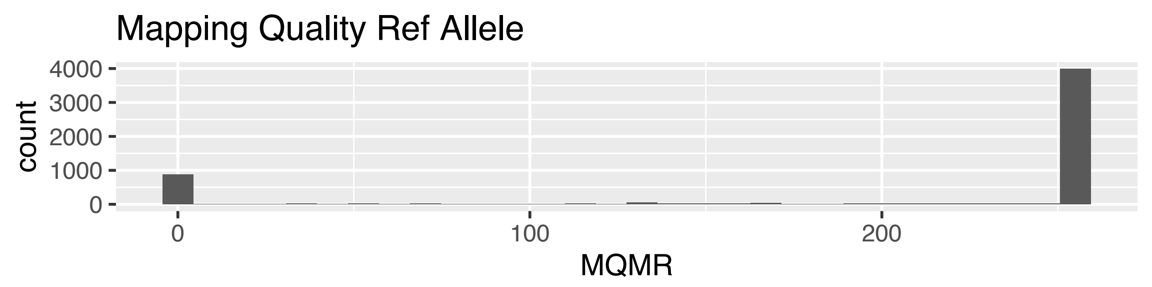
Mapping Quality Ref Allele
count
0
1000
2000
3000
4000
0
100
200
MQMR

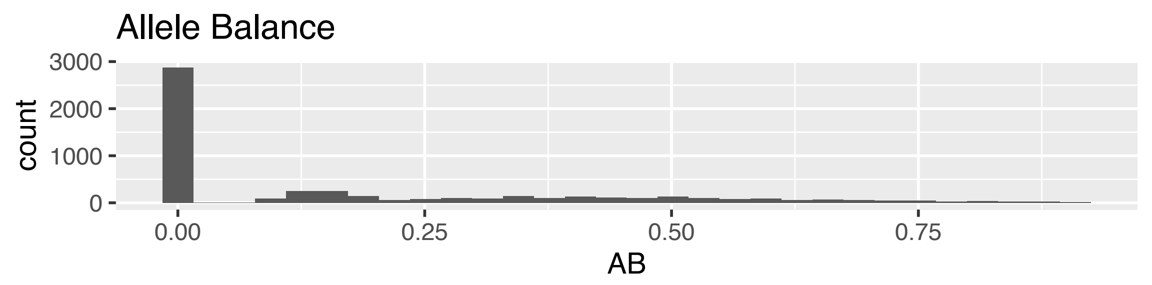
Allele Balance
count
0
1000
2000
3000
0.00
0.25
0.50
0.75
AB

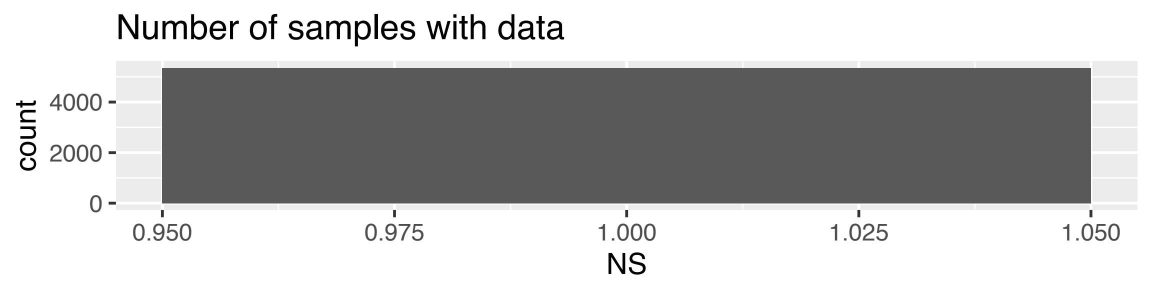
Number of samples with data
count
0
2000
4000
0.950
0.975
1.000
1.025
1.050
NS

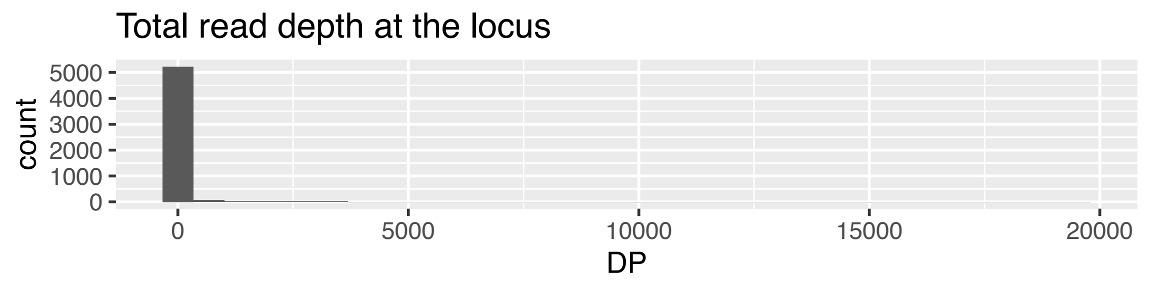
Total read depth at the locus
count
0
1000
2000
3000
4000
5000
0
5000
10000
15000
20000
DP

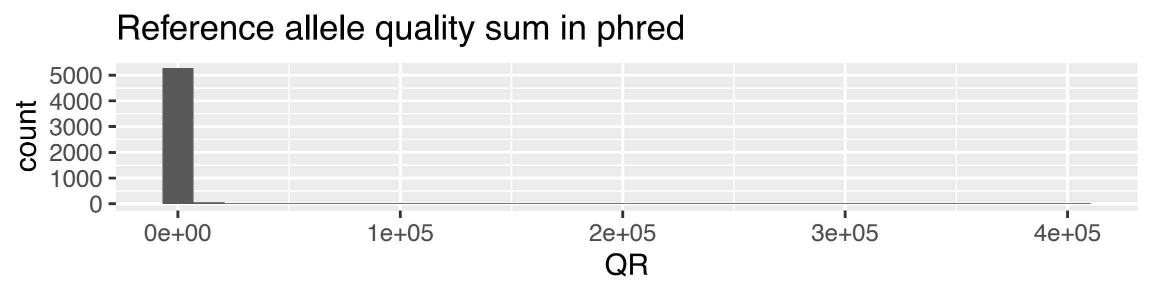
Reference allele quality sum in phred
count
0
1000
2000
3000
4000
5000
0e+00
1e+05
2e+05
3e+05
4e+05
QR

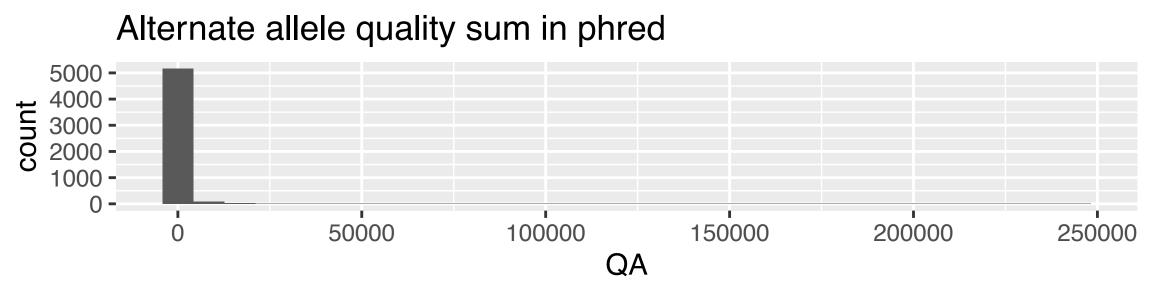
Alternate allele quality sum in phred
count
0
1000
2000
3000
4000
5000
0
50000
100000
150000
200000
250000
QA

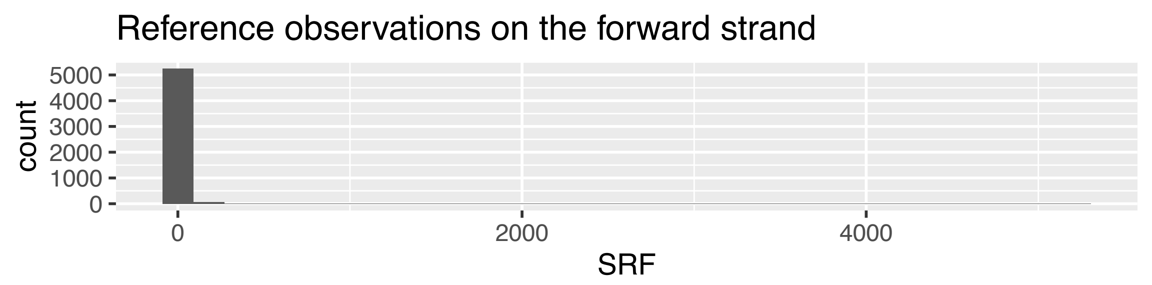
Reference observations on the forward strand
count
0
1000
2000
3000
4000
5000
0
2000
4000
SRF

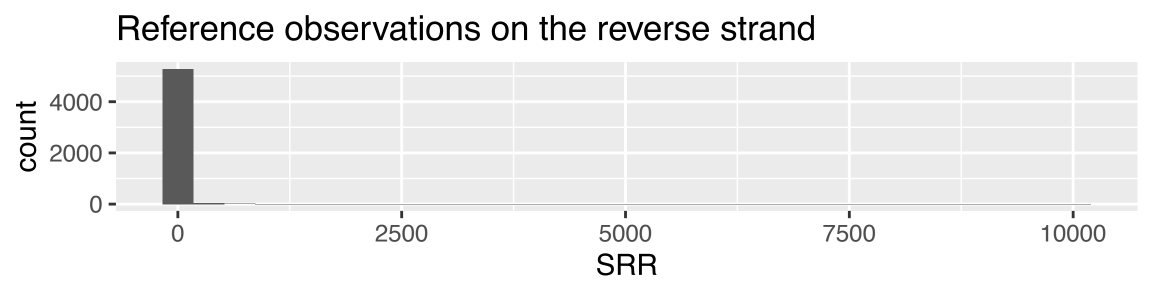
Reference observations on the reverse strand
count
0
2000
4000
0
2500
5000
7500
10000
SRR

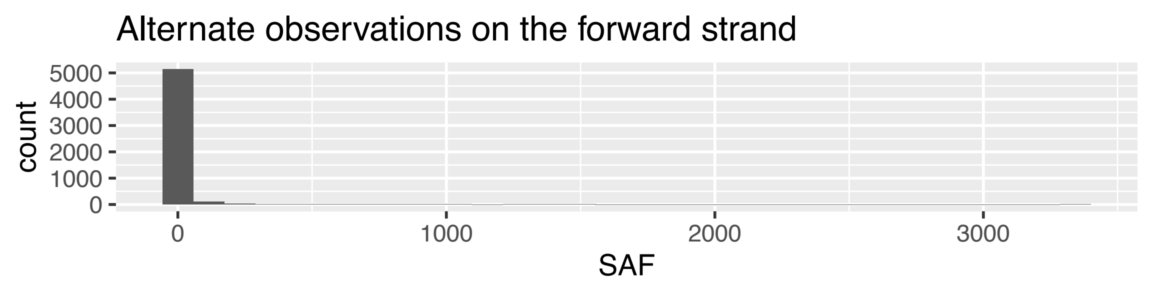
Alternate observations on the forward strand
count
0
1000
2000
3000
4000
5000
0
1000
2000
3000
SAF

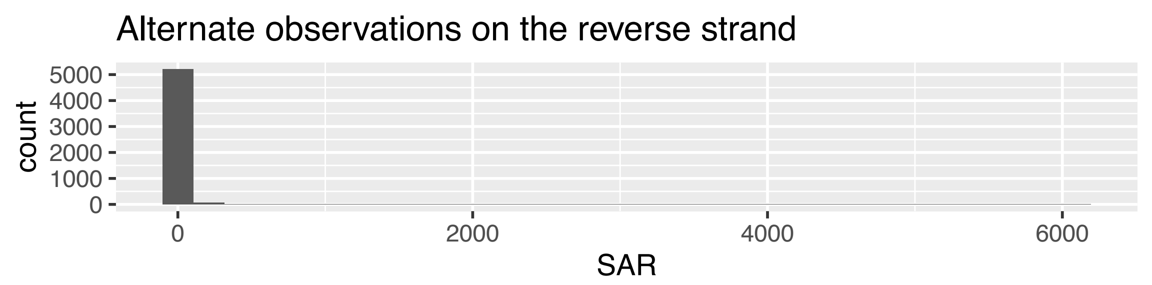
Alternate observations on the reverse strand
count
0
1000
2000
3000
4000
5000
0
2000
4000
6000
SAR

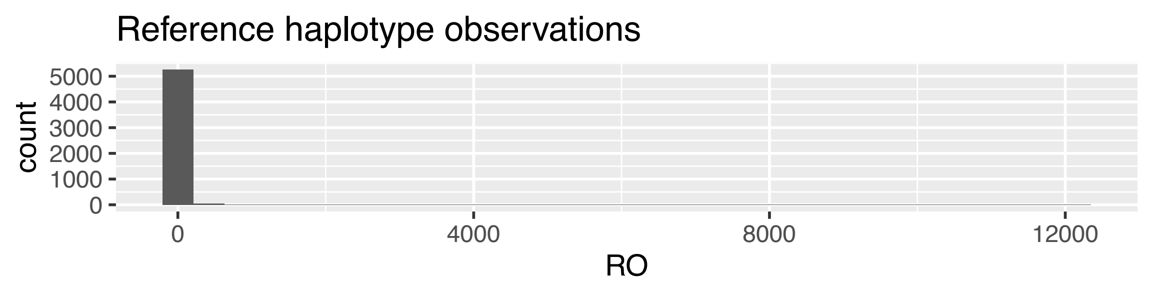
Reference haplotype observations
count
0
1000
2000
3000
4000
5000
0
4000
8000
12000
RO

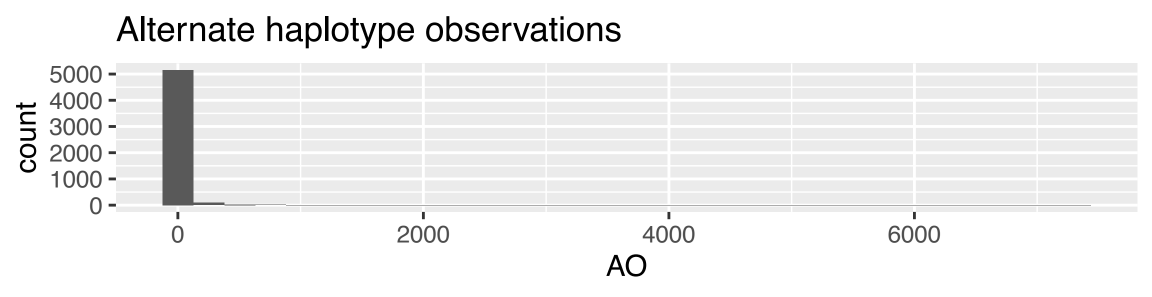
Alternate haplotype observations
count
0
1000
2000
3000
4000
5000
0
2000
4000
6000
AO

### Allele Balance by Variant Calls in Exons / 2023_14_35_ovovivi4dpf_11_plot2024−10−08

Mapping Quality >20, Readcounts >10

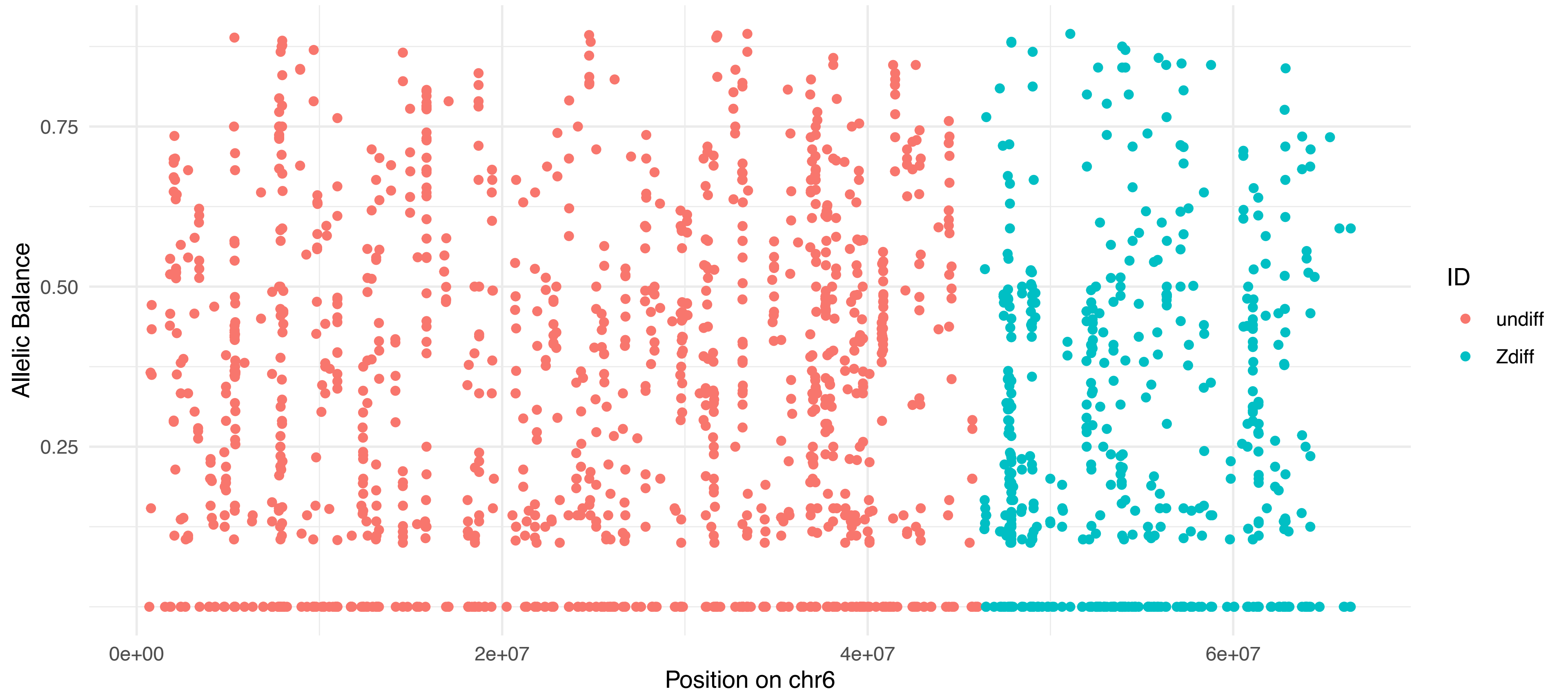


### Allele Balance by Variant Calls in Exons / 2023_14_35_ovovivi4dpf_11_plot2024−10−08

Mapping Quality >20, Readcounts >10

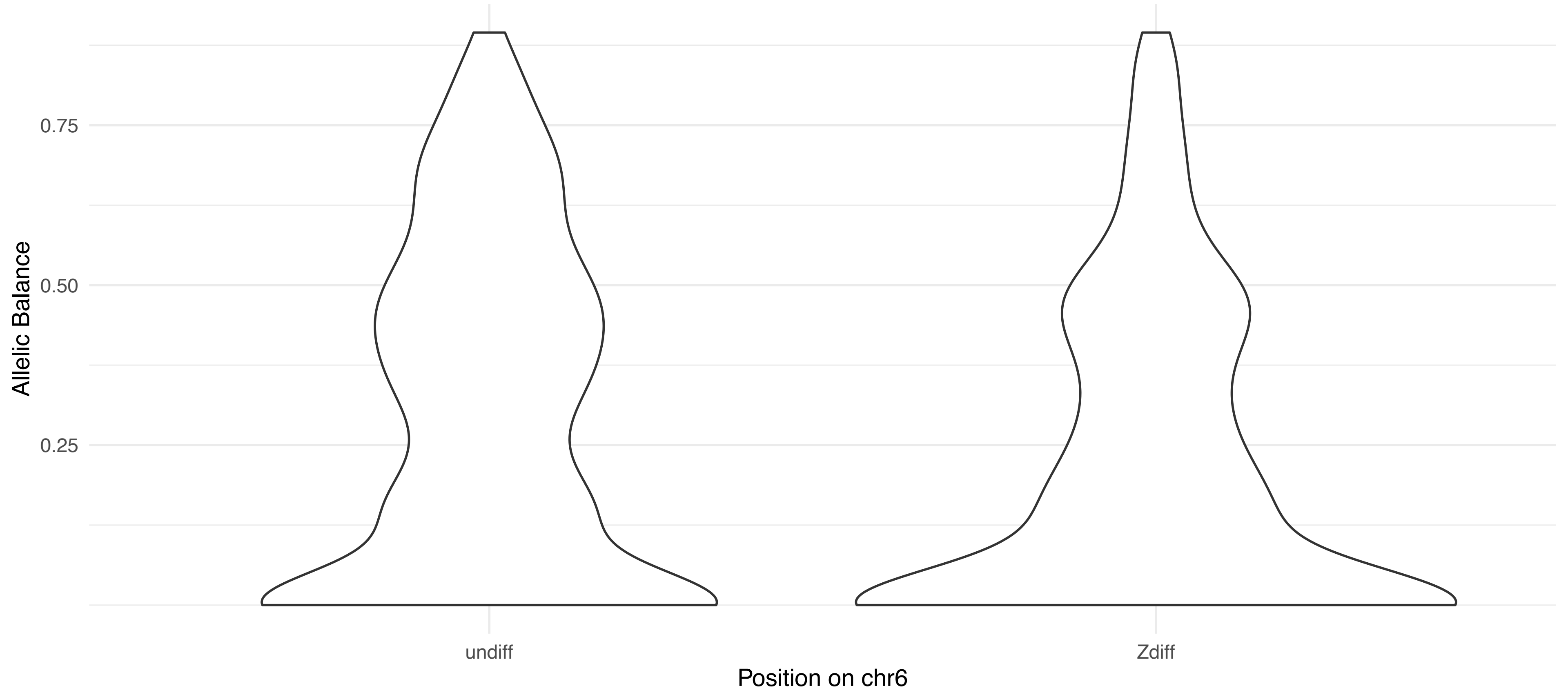


### Allele Balance by Variant Calls in Exons / 2023_14_35_ovovivi4dpf_11_plot2024−10−08

Mapping Quality >20, Readcounts >10

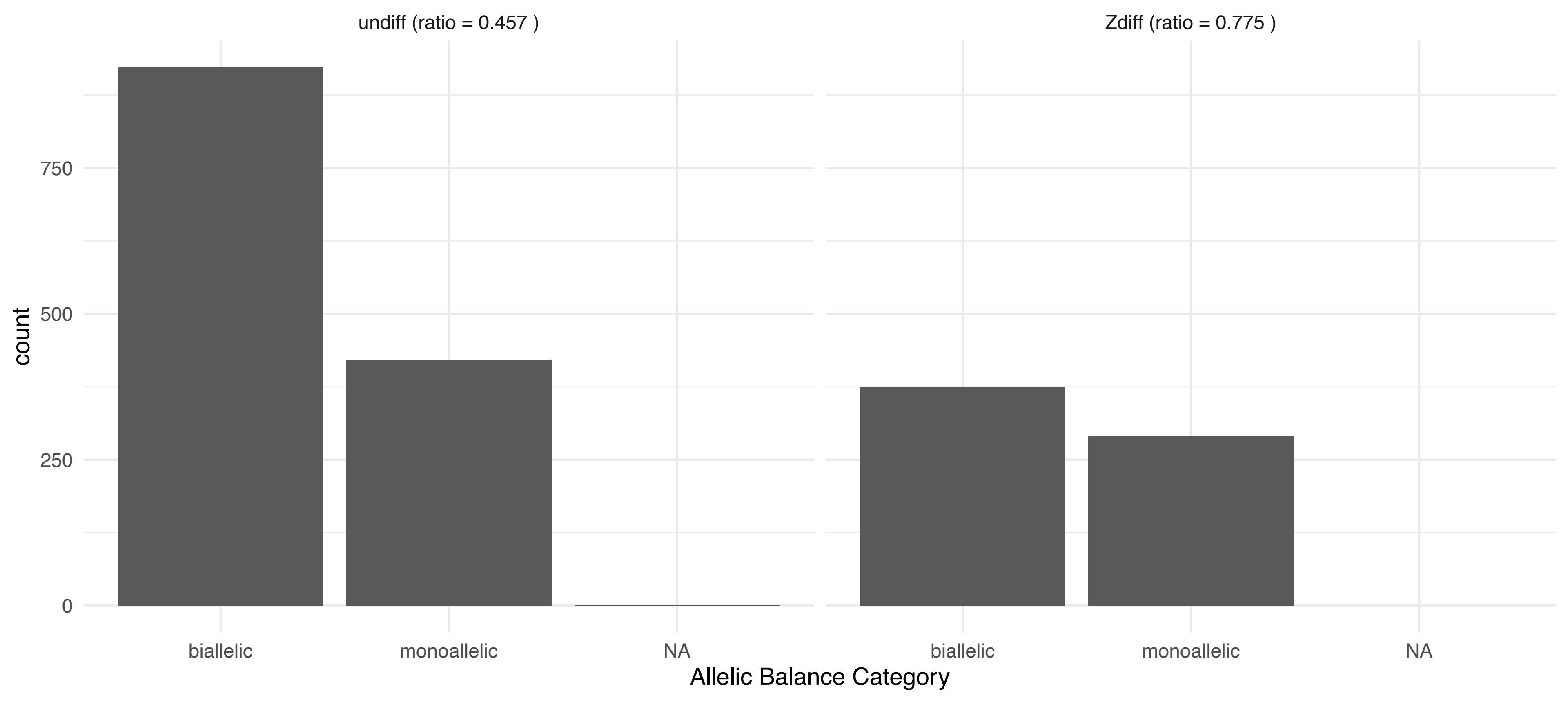

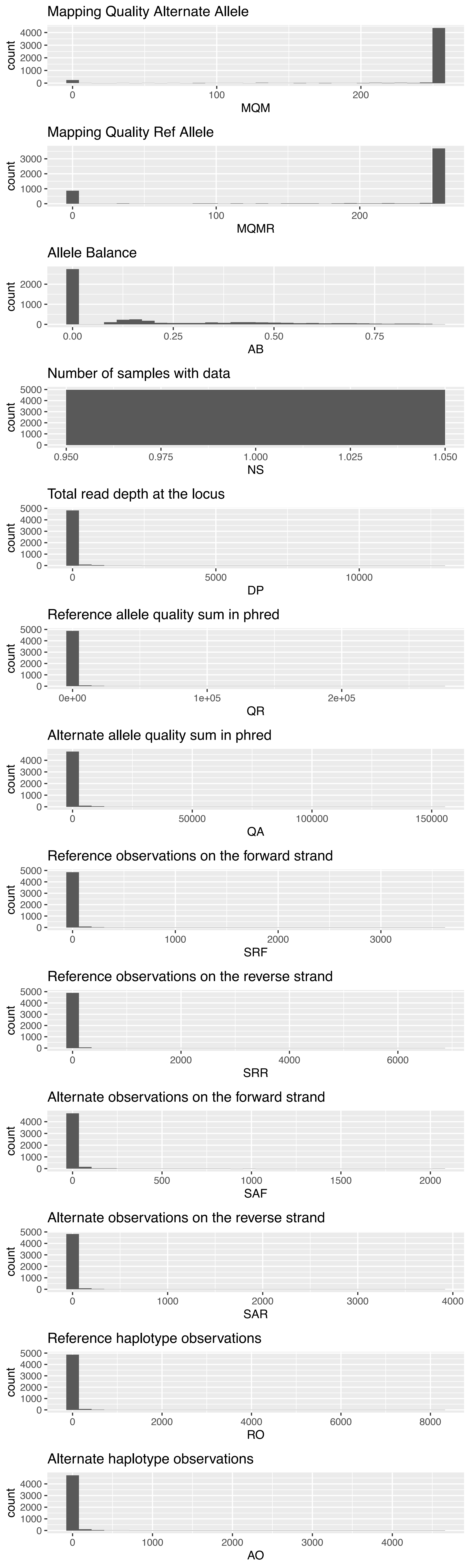
Mapping Quality Alternate Allele
count
0
1000
2000
3000
4000
0
100
200
MQM
Mapping Quality Ref Allele
count
0
1000
2000
3000
0
100
200
MQMR
Allele Balance
count
0
1000
2000
0.00
0.25
0.50
0.75
AB
Number of samples with data
count
0
1000
2000
3000
4000
5000
0.950
0.975
1.000
1.025
1.050
NS
Total read depth at the locus
count
0
1000
2000
3000
4000
5000
0
5000
10000
DP
Reference allele quality sum in phred
count
0
1000
2000
3000
4000
5000
0e+00
1e+05
2e+05
QR
Alternate allele quality sum in phred
count
0
1000
2000
3000
4000
0
50000
100000
150000
QA
Reference observations on the forward strand
count
0
1000
2000
3000
4000
5000
0
1000
2000
3000
SRF
Reference observations on the reverse strand
count
0
1000
2000
3000
4000
5000
0
2000
4000
6000
SRR
Alternate observations on the forward strand
count
0
1000
2000
3000
4000
0
500
1000
1500
2000
SAF
Alternate observations on the reverse strand
count
0
1000
2000
3000
4000
5000
0
1000
2000
3000
4000
SAR
Reference haplotype observations
count
0
1000
2000
3000
4000
5000
0
2000
4000
6000
8000
RO
Alternate haplotype observations
count
0
1000
2000
3000
4000
0
1000
2000
3000
4000
AO
